# Total workflows of the single-molecule imaging analysis in living cells: a tutorial guidance to the measurement of the drug effects on a GPCR

**DOI:** 10.1101/2020.06.08.141192

**Authors:** Masataka Yanagawa, Yasushi Sako

**Affiliations:** Cellular Informatics Laboratory, RIKEN Cluster for Pioneering Research, 2-1 Hirosawa, Wako, Saitama 351-0198, Japan

**Keywords:** single-molecule imaging, TIRFM, GPCR, ImageJ, single-molecule tracking analysis, VB-HMM, smDynamicsAnalyzer, diffusion dynamics, oligomer size distribution, colocalization analysis

## Abstract

Single-molecule imaging (SMI) is a powerful method to measure the dynamics of membrane proteins on the cell membrane. The single-molecule tracking (SMT) analysis provides information about the diffusion dynamics, the oligomer size distribution, and the particle density change. The affinity and on/off-rate of a protein—protein interaction can be estimated from the dual-color SMI analysis. However, it is difficult for trainees to determine quantitative information from the SMI movies. The present protocol guides the detailed workflows to measure the drug-activated dynamics of a G protein-coupled receptor (GPCR) and metabotropic glutamate receptor 3 (mGluR3), by using the total internal reflection fluorescence microscopy (TIRFM). This tutorial guidance comprises an open-source software named smDynamicsAnalyzer, with which one can easily analyze the SMT dataset by just following the workflows after building a designated folder structure (https://github.com/masataka-yanagawa/IgorPro8-smDynamicsAnalyzer).

## 1. Introduction

G protein-coupled receptors (GPCRs) are major drug targets, e.g., 34% of small molecule drugs target 6% of ~800 kinds of human GPCRs [1, 2]. Evaluation of the drug effects on a given GPCR often relies on monitoring cellular responses evoked by the drug-activated receptor [3]. However, such conventional approaches are difficult to investigate the drug effects on the majority of GPCRs whose signaling pathways are unclear, including ~80 non-olfactory orphan GPCRs [2].

We recently demonstrated that the diffusion dynamics of GPCR molecules in the plasma membrane (PM) is a versatile index to evaluate the drug effects on GPCRs [4]. We reported that commonly observed is agonist-dependent decrease of the average diffusion coefficient of various GPCRs, regardless of the receptor family and of the G protein coupling selectivity. We also revealed that the agonist-induced diffusion change of GPCR is partly related to the G protein coupling and the recruitment into the clathrin-coated pit (CCP) by dual-color single-molecule imaging (SMI) analysis. The activation-dependent decrease of the diffusion coefficient is also commonly observed with the epidermal growth factor receptor (EGFR) [5] and TRPV1 channel [6]. Therefore, the diffusion coefficient is a key index reflecting the functional states of various membrane receptors.

Here we describe the detailed protocols to determine the molecular dynamics of GPCRs in living cells with the total internal reflection fluorescence microscopy (TIRFM). TIRFM is a common SMI method [7–10], in which the fluorophore-labeled molecules within a limited depth ~200 nm above the coverslip are illuminated by the evanescent field on the glass—water interface (Fig. 1). This feature is suitable for observing various membrane proteins such as GPCRs and G-proteins, and has been used for the analysis of the dynamics of various GPCRs in the precedent studies [4, 11–16]. TIRFM allows us to selectively monitor the membrane proteins in the basal PM due to the reduction of the background fluorescence (FL) from the intracellular compartment and the apical membrane of the cell. When the density of the fluorophore-labeled molecule does not largely exceed 1/μm^2^, trajectories of the bright spots from single-molecules (or oligomeric particles) can be distinguished from one another.

**Fig. 1.**
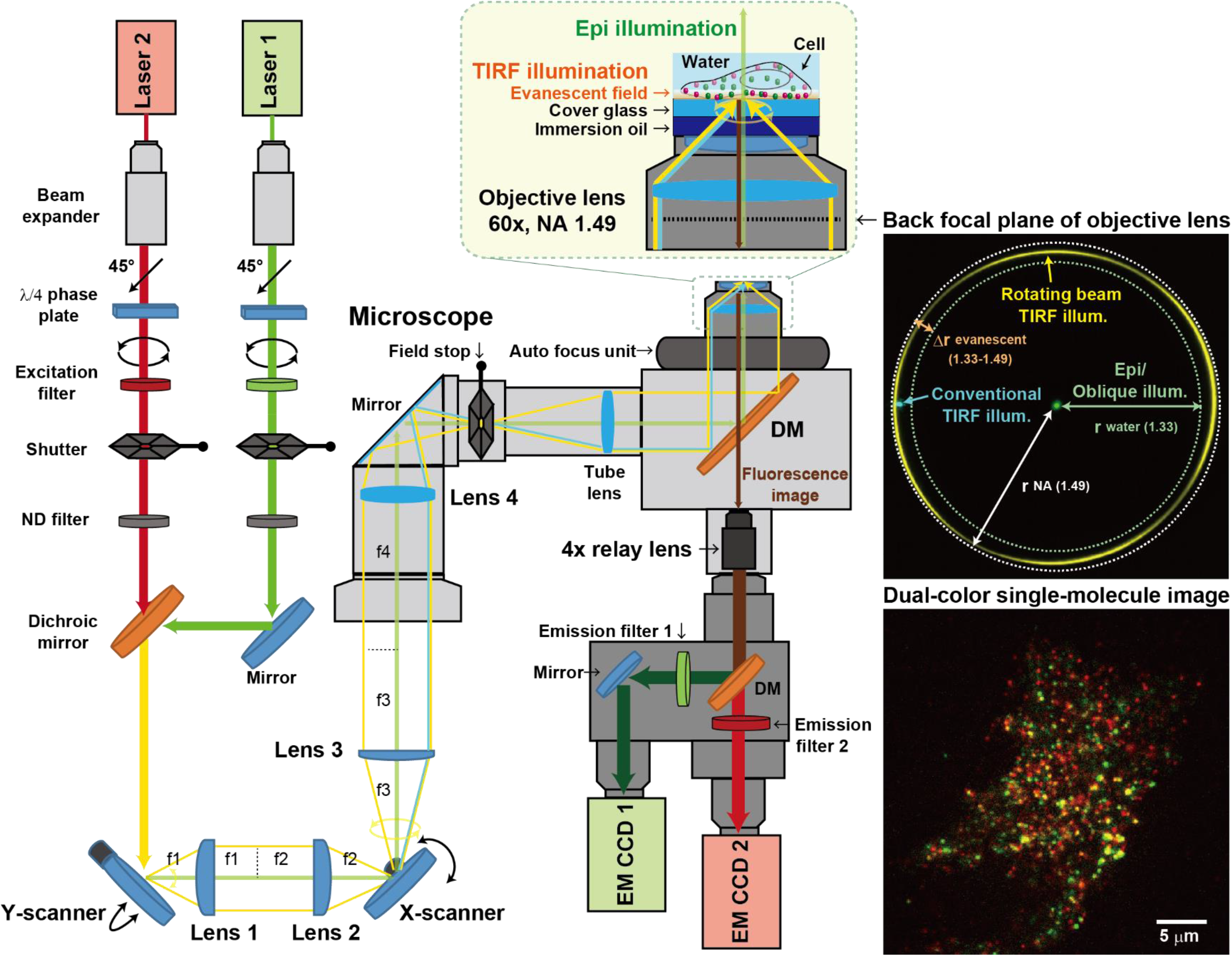
Optical configuration for the dual-color TIRFM. The images at the back focal plane of objective lens are taken by a CCD camera (Thorlab, 8051C-USB) through C-mount Camera Microscope adapter (MeCan, NYCZ). The dual-color single-molecule image is an example of TMR-labeled mGluR3 (red) and EGFP-labeled CLC (green).

The single-molecule tracking (SMT) analysis provides information on the localization, trajectory, and FL intensity of each particle. The particle density in the PM can be quantified from the localization data. The diffusion parameters can be obtained from the mean-squared displacement (MSD)-Δt plot of the trajectories and from the histogram analysis of step-size. The Variational Bayesian-Hidden Markov Model (VB-HMM) clustering analysis of the trajectories provides advanced parameters of diffusion, including the number of diffusion states, the diffusion coefficient and the fraction of each state in addition to the transition rate constants from a state to another. Statistics of the starting time and the length of trajectories provides information about on/off-rate constants of molecules. The distribution of oligomeric states of the molecules can be estimated from the FL intensity histogram analysis. Furthermore, the dual-color SMI analyses make it possible to quantify the diffusion parameters of two interacting molecules.

To date, various SMT analysis programs are available [17]; however, few programs have been published to calculate various kinetic parameters above from a large amount of the SMT dataset, and to perform a comparative analysis. This situation would make it difficult for trainees to perform single-molecule image analysis. Furthermore, the sample throughput of the SMI has been increased due to the development of the automated SMI system [5, 18], and the image analysis has become the rate-limiting phase of the experiments.

Considering these situations, the present protocol guides how to analyze the SMT dataset with our unique open-source software, named smDynamicsAnalyzer that can determine a series of kinetic parameters from the SMT dataset simply by following the designated steps after building an appropriate folder structure (https://github.com/masataka-yanagawa/IgorPro8-smDynamicsAnalyzer). As the tutorial guidance, we exemplify the analyses of metabotropic glutamate receptor 3 (mGluR3), a class C GPCR, and the clathrin-light chain (CLC) in detail.

## 2. Materials

### 2.1. PlasmidDNA (pDNA) vector encoding HaloTag/SNAP-tag/GFP-fusion

pDNA_HaloTag/SNAP-tag/GFP vectors encoding each target protein *(see* **Note 1**),which were inserted by In-Fusion HD Cloning Kit (Clontech) or Seamless Ligation Cloning Extract (SLiCE) from E.coli [19].

### 2.2. Cells

Human Embryonic Kidney cells 293 (HEK293 cells) (*see* **Note 2**)

### 2.3. Medium and buffer for HEK293 cells

1. Medium A (Culture medium): Dulbecco’s Modified Eagle Medium: Nutrient Mixture F-12 (DMEM/F12) (or DMEM) containing phenol red, supplemented with 15 mM HEPES (pH 7.3), 29 mM NaHCO_3_, and 10% fetal bovine serum (FBS).
2. Medium B (Medium for the FL-staining): DMEM/F12 (D2906-10L, Sigma) (or DMEM) without phenol red and NaHCO_3_ supplemented with 15 mM HEPES (pH 7.3) and 10% FBS.
3. Buffer A (Measurement buffer): Hanks’ balanced salt solution (HBSS, H1387-10L, Sigma) with 15 mM HEPES (pH 7.3) and 0.01% BSA, without NaHCO_3_.

### 2.4. FL dyes (see Note 3)

1. HaloTag ligands (Promega): HaloTag TMR, Janelia Fluor 549 (JF549), SaraFluor 650 (SF650, old product name: Stella Fluor 650) ligands, which are commercially available for the intracellular staining and the SMI.
2. SNAP-tag ligands (NEB): SNAP-Cell TMR-star, and 647-SiR ligands,which are commercially available for the intracellular staining and the SMI.

### 2.5. Equipment for TIRFM (Fig. 1)

1. An inverted fluorescence microscope (e.g. TE2000, TiE, Ti2E, Nikon, and IX81, IX83, Olympus).
2. An oil immersion objective with numerical aperture (NA) over 1.33 (e.g. PlanApo 60×, NA 1.49, Nikon, and Apo N 60×, NA 1.49, Olympus).
3. Lasers with an enough power and suitable wavelength to the fluorophores *(see* **Note 4**).
4. Optical shutter and its driver to turn the laser on and off (e.g. LS6 and VMM-D3, Uniblitz)
5. Optics to adjust the angle of the laser from epi illumination to TIRF illumination (*see* **Note 5**).
6. Dichroic and emission filters suitable for each laser and fluorophore (*see* **Note 6**).
7. Intermediate magnification lens (4× for 60× Objective) between the microscope and cameras.
8. EM-CCD camera (e.g. ImagEM, Hamamatsu) or sCMOS camera (e.g. ORCA-Flash4.0, Hamamatsu). Two cameras are used for the dual-color imaging.
9. A two-channel imaging system (W-view Gemini-2C, Hamamatsu) for the dual-color imaging.
10. A computer and a software (e.g. MetaMorph, Molecular Dvices) to control the optics, the microscope and cameras.

## 3. Methods

### 3.1. Preparation of coverslips for the SMI in living cells (see Note 7)

1. Coverslips (Matsunami, 25 mm round, No. 1; 0.13-0.17 mm thickness) for Attofluor Cell Chamber (Invitrogen).
2. Incubate the coverslips in a neutral detergent solution (e.g. 2% Clean-Ace S/water, AsOne) in a glass beaker with continuous sonication for 60-120 min.
3. Change the detergent solution to water several times followed by 5 min sonication.
4. Add coverslips into the concentrated sulfuric acid in a glass bottle with glass lid, and incubate over 16 h.
5. Remove the coverslips from the concentrated sulfuric acid, and put them into water in a glass beaker.
6. Change the water several times followed by 5 min sonication.
7. Repeat process 6 three times.
8. Autoclave the coverslips in the glass beaker with aluminum foil lid to avoid contaminations.
9. Incubate 2 coverslips in 3 mL medium A on a 60 mm dish over 16 h before seeding cells to increase the cell adhesion (coverslip dish).

### 3.2. Transfection by Lipofectamine 3000 (see Note 8)

1. Culture HEK293 cells to be 100 % confluence in a 10-cm dish on the day before the SMI.
2. Make transfection mixture A in a tube according to the following recipe:

Opti-MEM (Gibco): 60 μL
Lipofectamine 3000 reagent: 2.5 μL
3. Make transfection mixture B in a tube according to the following recipe:

Opti-MEM: 60 μL
pDNA vector: 0.1 μg for a vector carrying CMVd1promoter, (0.02 μg for a vector carrying CMV promoter) (*see* **Note 9**)
P3000 reagent: 0.2 μL (2x μL for x μg pDNA) (*see* **Note 10**)
4. Mix the mixtures A and B, and incubate over 15 min at room temperature (Mixture A+B).
5. Trypsinize the cells from the 10-cm dish with 1 mL 0.25% Trypsin-EDTA/PBS after 5 mL PBS rinse.
6. Remove the Trypsin-EDTA/PBS immediately after the treatment.
7. Suspend the cells into 4 mL medium A.
8. Remove the medium A from the coverslip dish (*see* Section 2.5.), and add 2.5 mL medium A.
9. Mix 500 μL of the suspended cells with the mixture A+B, and put it on the coverslip dish.
10. Incubate overnight at 37 °C under 5% CO_2_ (Day 0).

### 3.3. HaloTag/SNAP-tag staining (see Note 11)

1. Prepare the ligand solution by dissolving HaloTag or SNAP-tag ligands with DMSO to be 100 μM.
2. Freeze an aliquot of the ligand solution in different tubes before use to avoid repeated freeze-thawing.
3. Dilute the ligand solution to be 10~300 nM in 2 mL medium B per coverslip dish (staining solution).
4. Discard the medium A by suction from the coverslip dish, which was transfected Day 0, and add the staining solution carefully for avoiding detachment of the cells.
5. Incubate 15 min at 37 °C under 5% CO_2_.
6. Discard the staining solution by suction, and add 3 mL medium B carefully.
7. Prepare a new 6-cm dish containing 3 mL of medium B, and transfer the coverslips from the stained dish.
8. Discard the medium B from the dish by suction, and add 3 mL of fresh medium B carefully.
9. Incubate over 15 min at 37 °C under 5% CO_2_.
10. Repeat steps 8 and 9 three times.
11. Prepare each ligand for a target GPCR by dissolving it in buffer A to be a 5-fold higher concentration (i.e., 5x ligand solution).
12. Transfer a coverslip to the Attofluor Cell Chamber.
13. Wash the coverslip with 400 μL buffer A by a pipet three times carefully for avoiding detachment.
14. Add 400 μL of buffer A, and set the chamber on the microscope.
15. Add 100 μL of the ligand solution 15 min before imaging in the case of the measurement of a steady state of dynamics of GPCR.
16. In the case of the time-lapse imaging of the same cells, add 100 μL of the ligand solution between the time-points.

### 3.4. Single-molecule imaging (SMI)

1. Turn on all the equipment (microscope, camera, laser, shutter, and PC) and wait until the camera’s cooling temperature reaches to the appropriate set point (e.g. ImagEM: −65°C).
2. Set an objective to the microscope, and adjust the correction ring to the proper position.
3. Set the dichroic and emission filters to the proper position.
4. Activate an imaging software in the PC.
5. Put the immersion oil (e.g. IMMOIL-F30CC, Olympus) on the objective.
6. Put the sample on the oil. If any, eliminate bubbles between the objective and the coverslip.
7. Acquire images of multi-color beads (e.g. TetraSpeck, Thermo Fisher) for the dual-color imaging to calibrate and merge two channels. Adjust optics in the two-channel imaging system according to the manufacturer’s manual. (*see* **Note 12**)
8. Adjust the laser angle to an optimal position (*see* **Note 13**).
9. Adjust the camera settings to a high enough sensitivity to detect photons from a single fluorophore. (An example setting of ImagEM: exposure time: 30.5 ms, EM gain: 200, spot noise reduction: on)
10. Adjust the lens focus manually, and then keep it by using an auto focus system (e.g. Nikon PFS, Olympus ZDC)
11. Adjust the field stop to the center of the field of view of the camera, and widen the aperture to the minimum necessary.
12. Search cells with the density of single-molecules less than 1/μm^2^.
13. Acquire movies of ~20 cells for 100~400 frames (3~12 s with 30.5 ms exposure time) per cell during 15~30 min after ligand stimulation in the case of measuring a steady state of dynamics of GPCR.
14. Acquire time-lapse movies of ~20 cells for 100 frames per cell at each time point by using the Multi Dimensional Acquisition with the journal. (*see* **Note 14**).

### 3.5. Image processing by ImageJ

1. Run Background subtraction through following the path:

Process > subtract background > rolling ball radius: 25 pixels.
2. Run Running average through following the path: (optional processing)

Plugins > Stacks > Running_ZProjector > Size: 2, Type: average intensity (Running_ZProjector plugin can download from Vale Lab homepage, http://valelab.ucsf.edu/~nstuurman/ijplugins/)
3. Set Brightness and Contrast through following the path: (*see* **Note 15**)

Image > Adjust > Brightness/Contrast > Set (e.g. min: 0, max: 800)
4. Save as tif and/or avi files through following the path: (*see* **Note 15**)

File > Save As > Tif
File > Save As > Avi (Compression: None, Frame rate: 33 fps)

Alternatively,

1. Download the BatchImageProcessingSMT.ijm file (*see* **Note 16)** from the URL (https://github.com/masataka-yanagawa/ImageJ-macro-ImageProcessingSMT), and put it into the Macro folder of ImageJ.
2. Install the macro file through following the path: “Plugins > Macro > Install” and select the ijm file.
3. Run the macro “ImageProcessingSMT”.
4. Select a folder where multiple tif files are stored.
5. Check the check boxes for the required processing items, and enter the parameters.
6. Push OK button.
7. Processed tif and/or avi files with a suffix are saved in the same folder.

### 3.6. Additional image processing by ImageJ for the dual-color analysis

1. Split channel 1 and channel 2 if the format of tif files is 1024 × 512 pixel (pix). (e.g. the imaging data taken by two ImagEMs using the Metamorph)

i. Install the macro file as shown above.
ii. Run the macro “Split2ch”
iii. Select a folder where multiple tif files (1024 × 512 pix) are stored.
iv. Processed tif files with the suffix “-ch1” or “-ch2” are saved in the same folder.
v. Run the macro “ImageProcessingSMT” to the split images as shown above.
2. Composite 2 channels (*see* **Note 17**)

i. Install the macro file as shown above.
ii. Make 2 subfolders named “C1” and “C2 below the same folder.
iii. Move multiple tif files with the suffix “-ch1” and “-ch2” to “C1” and “C2” folders, respectively.
iv. Run the macro “Composite2ch”
v. Select the folder, where “C1” and “C2” are stored.
vi. Processed tif files with the suffix “_composite” are automatically saved in the selected folder
3. Align 2 channels by using affine transformation (*see* **Note 18**):

i. Open the multi-color bead image composited, and calculate parameters of affine transformation through following the path: Plugins > jars > GridAligner > Push “Calculate from Image” button
ii. Open the single-molecule image composited, and push “Apply” button.
iii. Save the aligned image through following the path, if necessary: File > Save As > tif and/or avi
iv. Split 2 channels of the aligned image through following the path: Image > Color > Split Channels
v. Save each channel of the aligned image through following the path: File > Save As > tif and/or avi

By using the AAS (Zido, https://zido.co.jp/en/), one may skip Steps 2 and 3 as follow:

i. Install and activate the AAS (User authentication is required).
ii. Push “Start this analysis” button on the “Affine Transformation” (Fig. 2a).
iii. Make “afine1”, “affine 2”, and “data” folders.
iv. Put the ch1 and the ch2 of the multi-color bead images in the “affine1” and “affine 2” folders, respectively.
v. Put the ch2 of the single-molecule images before affine transformation.
vi. Enter the parameters for the single-particle detection to analyze the localization of the bright spots of the ch1 and the ch2 of the multi-color bead images (*see* Section 3.6 for the detailed algorism). (e.g. Region of interest (ROI) size: 12 pix, scan length: 3 pix, connection distance: 8 pix, Light intensity threshold 1: 500, Light intensity threshold 2: 500)
vii. Click “Start analysis” button, then all the images in “data” folder will be output in the “data2” folder after affine transformation.

**Fig. 2.**
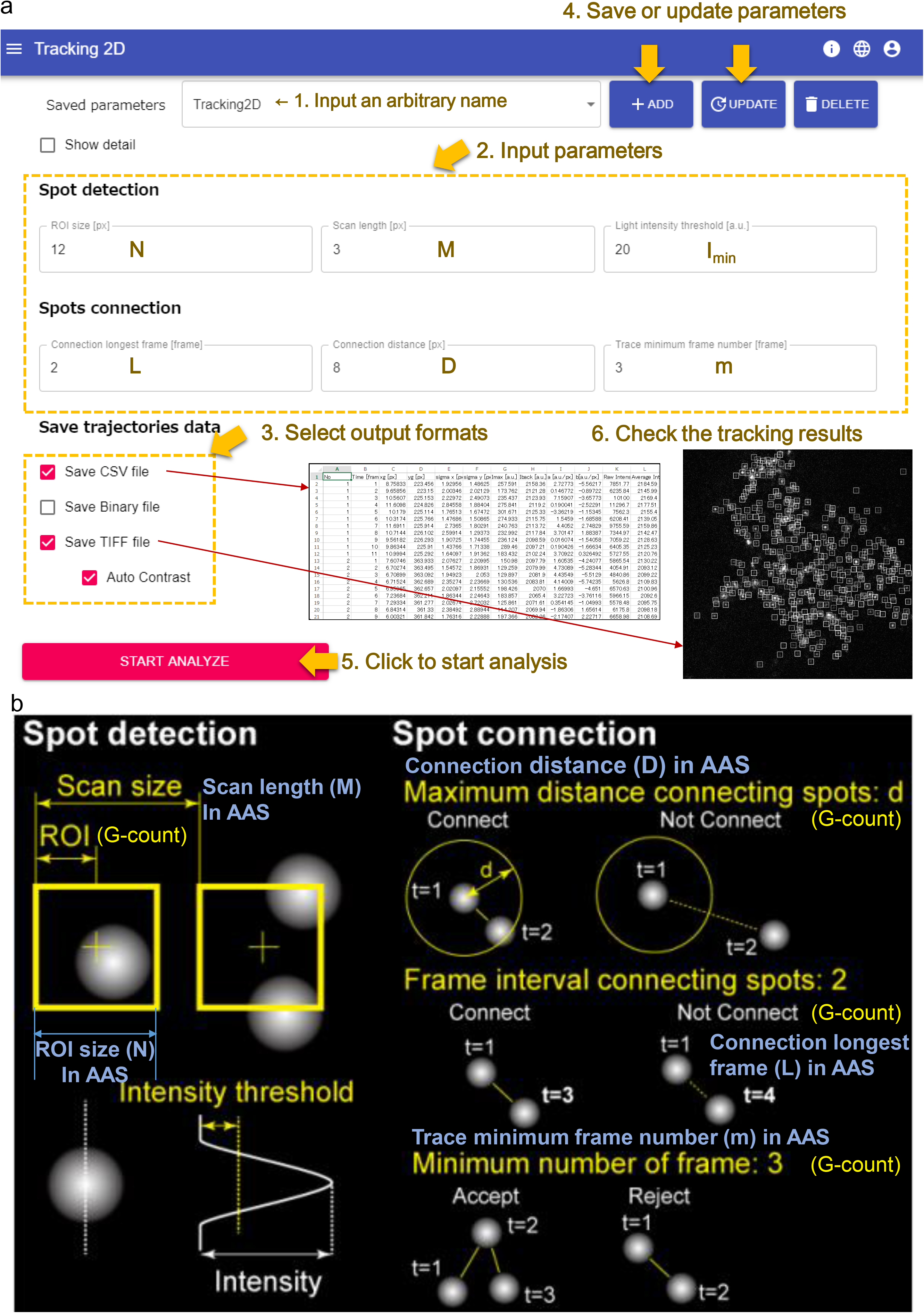
Screen shot of the AAS; (a) SMT analysis program (Tracking2D). (b) Illustration of SMT analysis algorithm.

### 3.7. Single-molecule tracking (SMT) analysis by AAS (see Note 19)

1. Put all the multiple tif (8 or 16 bit) files to be analyzed within a folder.
2. Install and activate the AAS (Zido, https://zido.co.jp/en/).
3. Push “Start This Analysis” button on the “Tracking2D”.
4. Enter the parameters for the SMT analysis (Fig. 2).
5. Select output options of AAS.
6. Push “Start Analyze” button, then the AAS output the analysis results (csv and tif files) in the same folder.
7. Perform the VB-HMM analysis of the diffusion dynamics (optional)

### 3.8. Analysis of the single-molecule dynamics of GPCRs by using smDynamicsAnalyzer

#### 3.8.1. Installation of Igor and building a folder structure to use smDynamicsAnalyzer (*see* Note 20)

1. Download and install Igor Pro 8 (abbreviated to Igor hereafter) (https://www.wavemetrics.com/products/igorpro).
2. Download the smDynamicsAnalyzer.pxp file (https://github.com/masataka-yanagawa/IgorPro8-smDynamicsAnalyzer). Open it by double click. Increase the resolution of the screen if the parameter and analysis button panels cannot fit into the screen (Fig. 3a).
3. Set parameters in the parameter panel (*see* **Note 21**).
4. Select an input format (Fig. 3a, *see* **Note 22**).
5. Store the data files taken under the same condition in a single folder (Fig. 3b). At least 3 files are required to calculate the statistics. Do not open the csv or xlsx files by excel when starting the analysis on the smDynamicsAnalyzer.
6. Build a folder structure as shown in Fig. 3b. In this example, five folders (“A1”~”A5”) each of which containing the 20 xlsx files with G-count format are built under “LY34” folder.
7. Check the input format checkbox in the “Measurements/SMT parameters” panel to fit the data format (Fig. 3a).
8. Push “Total Basic Analysis” button and select the folder containing the subfolders to be compared, then all the analyses of the selected folder are automatically performed (*see* 3.8.12, 3.8.21, and **Note 23**).

#### 3.8.2. Respective analysis of the AAS and G-count data formats: Total analysis of the data in a folder (Fig. 4a)

1. Make a folder containing at least three sample data files. In Fig. 4, we made a “test” folder containing three xlsx files (test1, test2, test3) with G-count format for the test run to set proper parameters.
2. Click the “Respective Analysis” button after setting the parameters in the parameter panel.
3. Input an arbitrary “Sample Name” (e.g. “test”) in the pop-up panel, then click the “continue” button.
4. Select the folder containing the data files. (e.g. the “test” folder)
5. Click the “Respective Analysis” button, which starts all the macros in the box below sequentially. Load → Trace → MSD-dt → Hist D → Intensity → Density → Off-rate → On-rate → stats (*see* Sections 3.8.3 ~ 3.8.11 regarding each macro.)
6. Temporarily ignore the error and wait until the macro finishes if you get an error message caused by the poor fitting of the graph during the process. Push “continue” button in case you set the debugger on.
7. Restart the macro, if a poor curve fitting error pops up, by clicking the appropriate button again after changing the parameters in the parameter panel. (e.g. Click “MSD-dt” button if a poor curve fitting error pops up in the process of MSD-Δt plot analysis.)
8. Perform the parameter optimization until no more error signs are posted in the fitting results for each item (*see* **Note 24**).

**Fig. 3.**
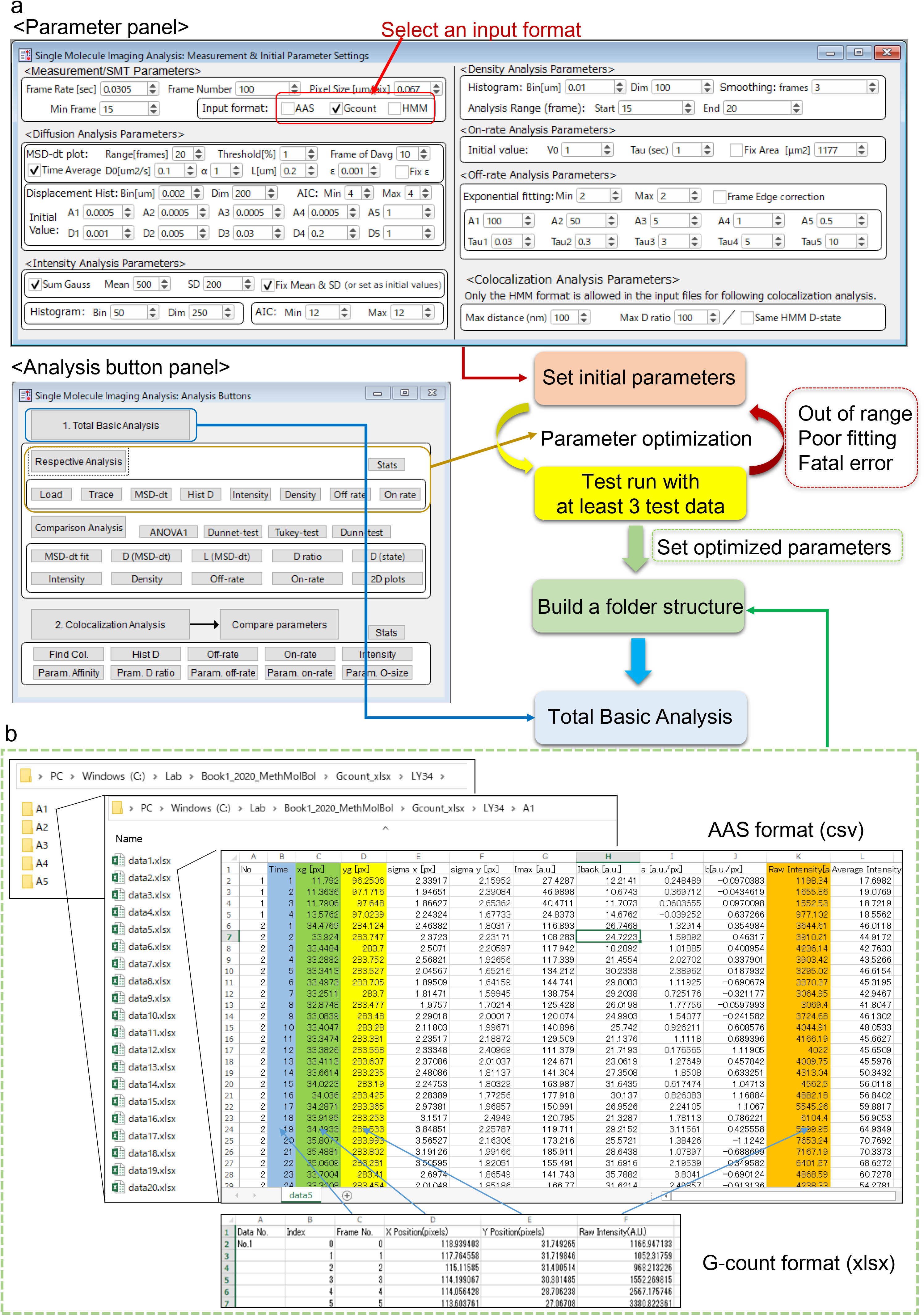
Summary of the use of the smDynamicsAnalyzer. (a) Screen shots of the parameter panel (top) and the analysis button panel (bottom). The basic workflow is depicted as a flow chart. (b) The folder structure containing the SMT data sets to load into the smDynamicsAnalyzer. The AAS (csv file) and the G-count (xlsx file) formats are exemplified.

#### 3.8.3. Load: Load input data (Fig. 4b)

1. Click the “Load” button after building a folder structure as shown in section 3.8.2.
2. Input an arbitrary “Sample Name” (e.g. “test”) in the pop-up panel, then click the “continue” button.
3. Select the folder containing the data files (e.g. the “test” folder), where the Macro automatically creates “Sample Name” folder under the root folder, then “Folder Name” folders under “Sample Name” folder in Igor, (i.e., “Folder Name” = “Sample Name” + file number). The macro also makes waves that are loaded from a data file into the “Folder Name” folder, and outputs tables loaded (*see* **Notes 25~29**).

**Fig. 4.**
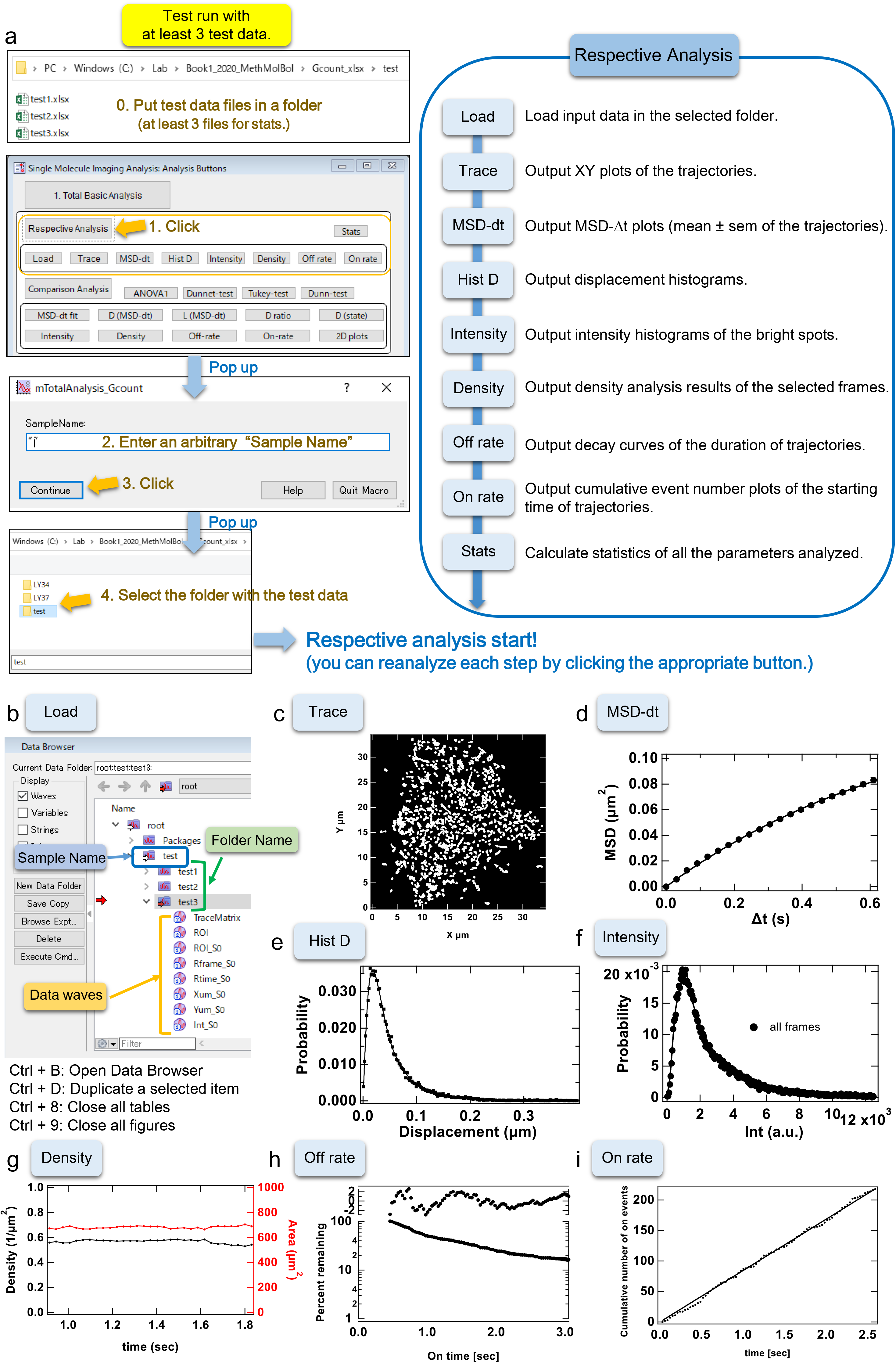
Guidance on “Respective Analysis” in the parameter optimization using practical test data (G-count format data). (a) The workflow of “Respective analysis”. (b) Structure of the Igor data folders built by “Load” macro (c) Trajectories of mGluR3 output generated by the “Trace” macro. (d) An MSD-Δt plot of mGluR3 output generated by the “MSD-dt” macro. (e) A displacement histogram of mGluR3 output generated by the “Hist D” macro. (f) An intensity histogram of mGluR3 output generated by the “Intensity” macro. (g) A density/area plot of the spots of mGluR3 in a cell output made by the “Density” macro. The estimated density (black, left axis) and area (red, right axis) are plotted versus time. (h) A decay curve of the duration of the mGluR3 trajectories output made by the “Off-rate” macro. (i) A cumulative event number plot output made by the “On-rate” macro.

#### 3.8.4. Trace: Output XY plots of trajectories (Fig. 4c)

1. Click the “Trace” button.
2. Input the “Sample Name” that corresponding a folder name under the root folder in the popup panel.
3. Click the “continue” button, where the macro automatically outputs graphs of the trajectories in each “Folder Name” folder (Yum_S0 vs Xum_S0 in Fig. 4b).
4. Select a ROI in the trace graph and right click to select “Expand” to visualize trajectories in the expanded area (*see* **Notes 28~31**).

#### 3.8.5. MSD-dt: Output MSD-Δt plots (Fig. 4d)

1. Click the “MSD-dt” button after setting the parameters in the parameter panel.
2. Input the “Sample Name” that corresponding a folder name under the root folder in the popup panel.
3. Click the “continue” button, where the macro automatically generates graphs of the MSD-Δt plots and the related tables (*see* **Note 32**).

#### 3.8.6. Hist D: Output displacement histograms (Fig. 4e)

1. Click the “Hist D” button after setting the parameters in the parameter panel.
2. Input the “Sample Name” that corresponding a folder name under the root folder in the popup panel.
3. Click the “continue” button, where the macro automatically produces graphs of the displacement histograms and the related tables (*see* **Note 33**).

#### 3.8.7. Intensity: Output intensity histograms (Fig. 4f)

1. Click the “Intensity” button after setting the parameters in the parameter panel.
2. Input the “Sample Name” that corresponding a folder name under the root folder in the popup panel.
3. Click the “continue” button, where the macro automatically outputs graphs of the intensity histograms and the related tables (*see* **Note 34**).

#### 3.8.8. Density: Output density analysis results. (Fig. 4g and Fig 5)

1. Click the “Density” button after setting the parameters in the parameter panel.
2. Input the “Sample Name” that corresponding a folder name under the root folder in the popup panel.
3. Click the “continue” button, where the macro automatically outputs XY-plots of the localization of spots, the mean local density plot, and the graphs and tables of the estimated density and area (*see* **Note 35**).

**Fig. 5.**
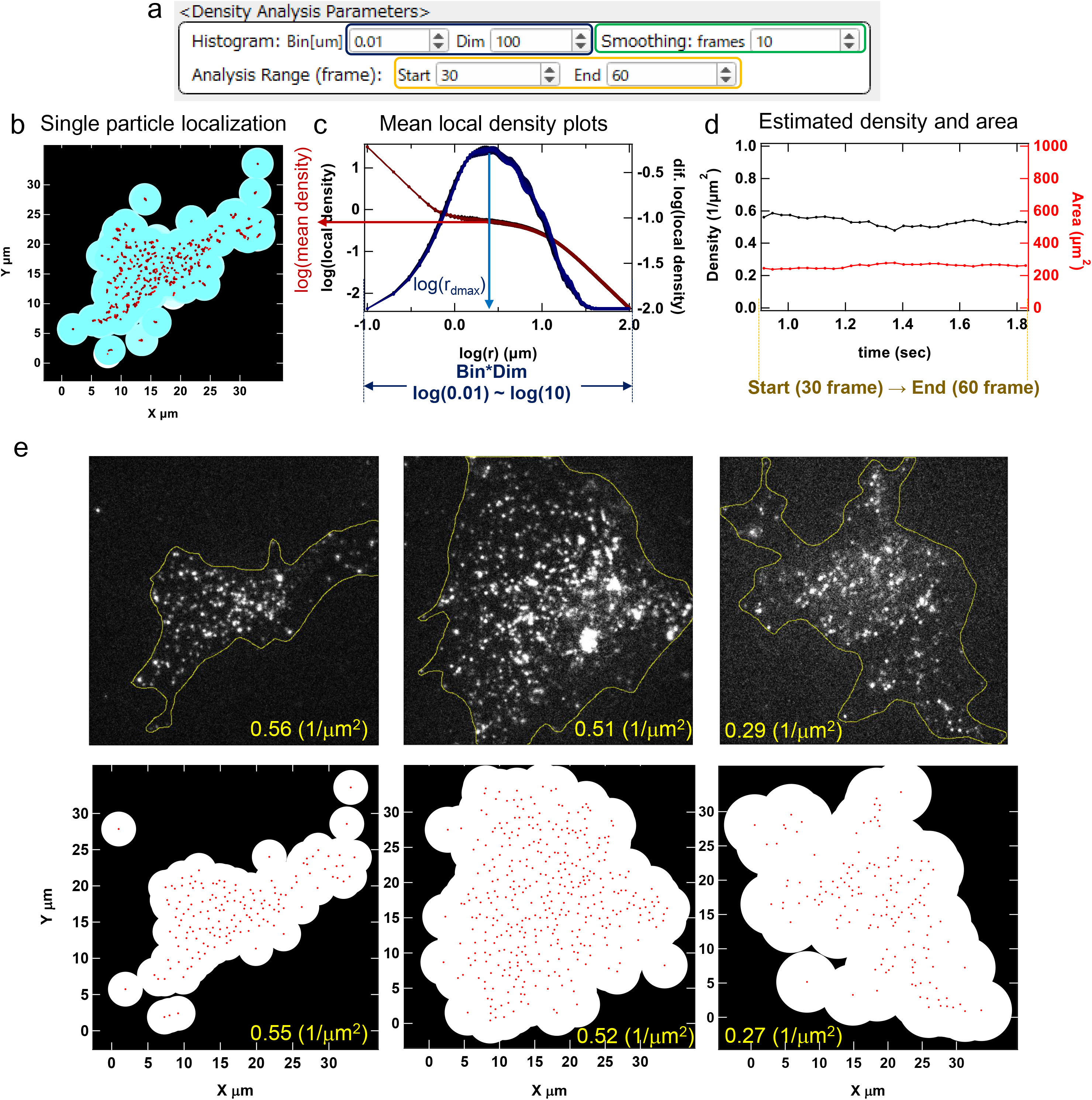
Guidance on “Density” analysis macro. (a) The part of the parameter panel related to the “Density” analysis. (b) The single-particle localization plot. The XY coordinates of spots were plotted by red dots. The local area around each spot given by the mean local density analysis was depicted by blue-white circles. (c) The mean local density (red, left axis) and its first order difference (blue, right axis) were plotted versus log(r). (d) The density (black, left axis) and area (red, right axis) estimated from (c) at each frame were plotted versus time. (e) Comparison between the mean densities estimated by hand and by the “Density” macro. Yellow lines in the upper panels were depicted by hand. The densities in the upper panels (yellow characters) were calculated as spot number/area of a frame. The densities in the lower panels were estimated by the mean local density analysis of the same frame.

#### 3.8.9. Off rate: Output decay curves of the duration of trajectories. (Figs. 4h and 6)

1. Click the “Off rate” button after setting the parameters in the parameter panel.
2. Input the “Sample Name” that corresponding a folder name under the root folder in the popup panel.
3. Click the “continue” button, where the macro automatically outputs decay curves of the duration of trajectories, and the related tables (*see* **Notes 36 and 37**).

**Fig. 6.**
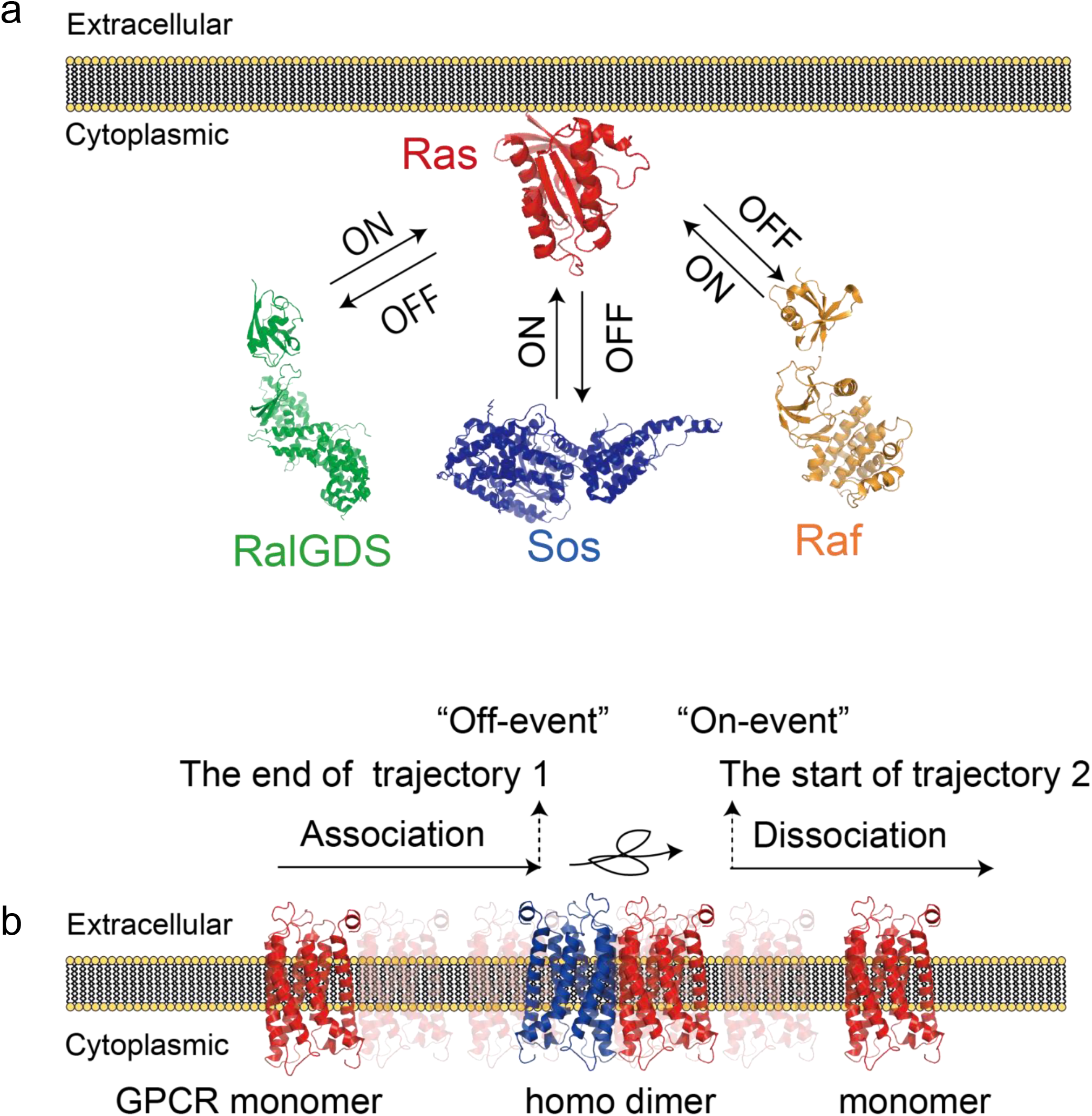
The interpretation of “On-rate” and “Off-rate” of the cytoplasmic proteins (a) and of the transmembrane proteins (b). Both figures are assumed a steady-state model.

#### 3.8.10. On rate: Output cumulative event number plots of the starting time of trajectories. (Figs. 4i and 6)

1. Click the “On rate” button after setting the parameters in the parameter panel.
2. Input the “Sample Name” that corresponding a folder name under the root folder in the popup panel.
3. Click the “continue” button, where the macro spontaneously generates cumulative event number plots of the starting time of trajectories, and the related tables (*see* **Notes 36 and 38**).

#### 3.8.11. Stats: Calculate statistics of all the parameters analyzed

1. Click the “Stats” button after setting the parameters in the parameter panel.
2. Input the “Sample Name” that corresponding a folder name under the root folder in the popup panel
3. Click the “continue” button, where the macro automatically outputs Matrix and Results folders under the “Sample Name” folder. (*see* **Note 39**).

#### 3.8.12. Total Basic Analysis of the AAS and G-count data formats: (Fig. 7)

1. Set the optimized parameters in the parameter panel (Fig. 3a).
2. Build the folder structure as shown in Fig. 3b and 7 (*see* **Note 40**).
3. Check again the input format checkbox in the “Measurements/SMT parameters” panel to fit the data format.
4. Click the “Total Basic Analysis” button.
5. Select the folder containing the subfolders to be compared (e.g. Select the “LY34” folder containing “A1” ~ “A5” subfolders in Fig. 7).
6. Click the “select folder” button in the pop-up panel, where the “Total Basic Analysis” automatically starts “Respective analysis” of each folder sequentially followed by “Comparison Analysis” (*see* **Note 41**).

**Fig. 7.**
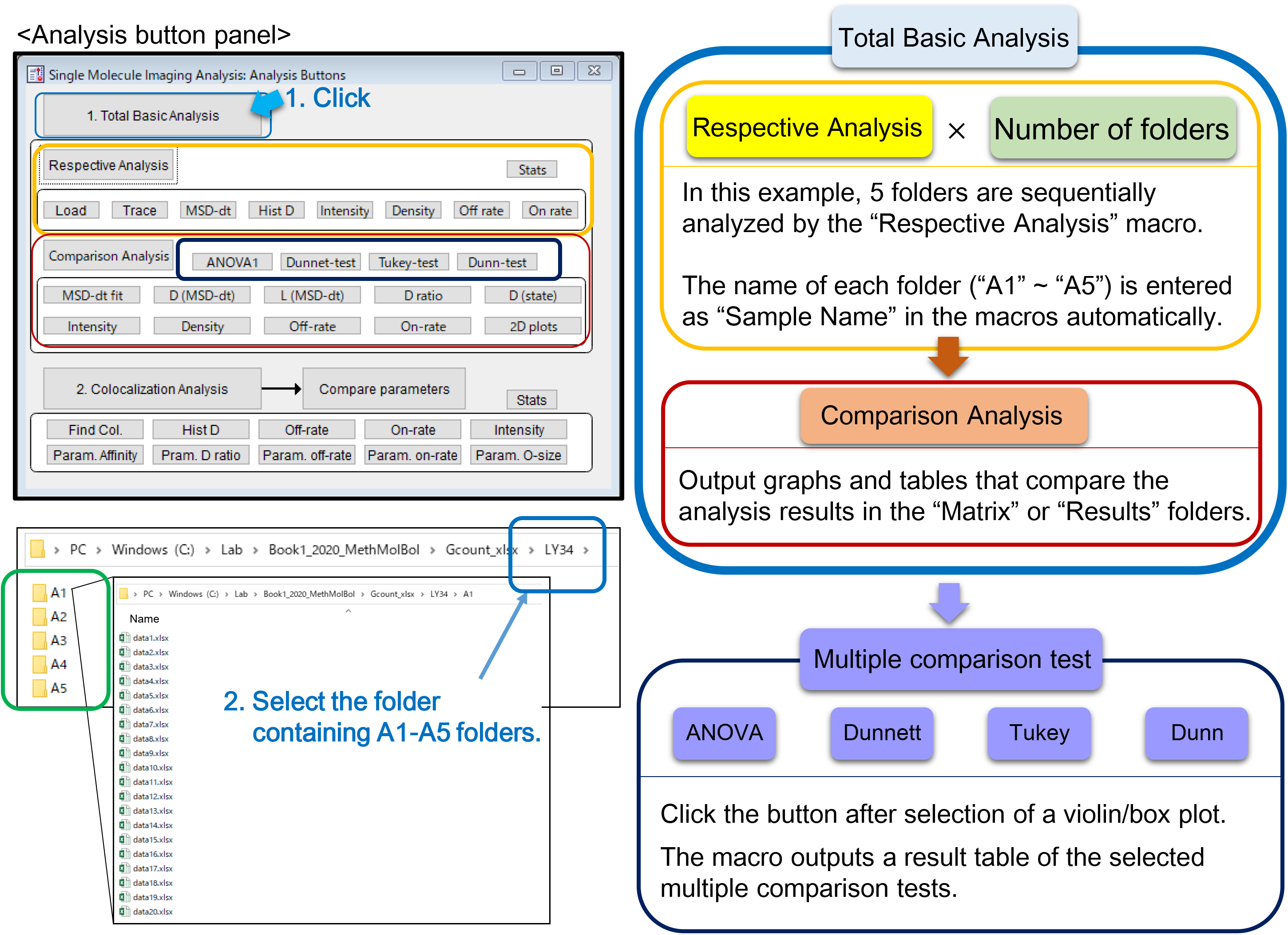
The work flow of “Total Basic Analysis” and “multiple comparison test” to analyze and compare the parameters of mGluR3 in five different ligand conditions (G-count format data). The inverse agonist LY341495 concentrations are A1: 0 nM, A2: 1 nM, A3: 10 nM, A4: 100 nM, A5: 1000 nM.

#### 3.8.13. Comparison Analysis of the AAS and G-count data formats: Total comparison analysis, and multiple comparison tests (Fig. 7 and 8)

1. Perform more than twice of “Respective Analysis” before starting “Comparison Analysis”. (*see* **Note 29**)
2. Click the “Comparison Analysis” button (*see* **Note 42**), where the click starts the “Comparative Analysis”, i.e., all the macros below sequentially. MSD-dt → D_MSD-dt_ → L_MSD-dt_ → D_ratio_ →D_state_ → Intensity → Density → Off-rate → On-rate → 2D plots. (*see* Sections 3.8.14 ~ 20 regarding each macro.)
3. Temporarily ignore the errors and wait until the macro finishes if you get an error message due to a poor fitting of the graph during the process. Push “continue” button in case you set the debugger on.
4. Restart the macro, where a poor curve fitting error pops up, by clicking the corresponding button after changing the parameters in the parameter panel.
5. Perform one-way ANOVA and multiple comparison tests among the groups on a violin/box plot by clicking the “corresponding” button after selecting a violin/box plot (Analysis button panel in Fig. 7, dark blue square). (*see* **Note 43**)

**Fig. 8.**
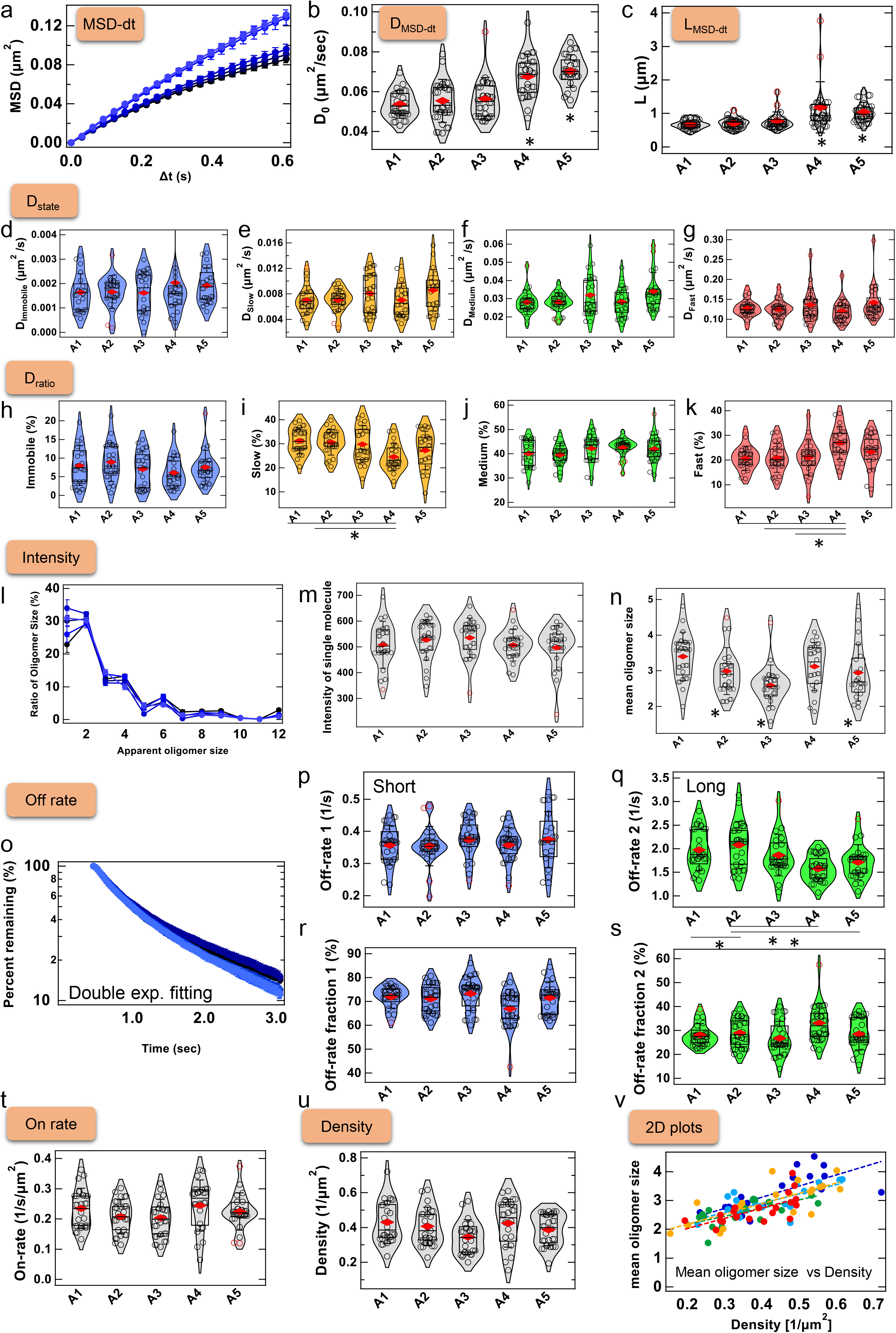
Guidance on “Comparison Analysis” macro. Example results of “MSD-dt” (a), “D_MSD-dt_” (b), and “L_MSD-dt_” macros (c). *p < 0.05 by the one-way ANOVA with the post-hoc Dunnett-test. Example results of “D_state_” (d-g) and “D_ratio_” (h-k) macros. *p < 0.05 by the one-way ANOVA with the post-hoc Tukey-test. Example results of “Intensity” macro (l-n). (l) The line-graphs of the oligomer size distribution (mean ± sem of the cells) estimated from the “An” values in equation 9 (*see* **Note 31**). The violin/box plots of the intensity of the single-molecule (m) and the mean oligomer size (n). *p < 0.05 by the one-way ANOVA with the post-hoc Dunnett-test. (o-s) Example results of “Off-rate” macros. (o) The decay curves of the duration of trajectories. The violin/box plots of the time constants (p: short, q: long) and of the fraction (r: short, s: long) estimated from the double-exponential fittings. *p < 0.05 by the one-way ANOVA with the post-hoc Tukey-test. Example results of “On-rate” (t), “Density” (u) and “2D plots” macros (v: mean oligomer size vs density). See **Note 37** for the violin/box plot settings.

#### 3.8.14. MSD-dt, D_MSD-dt_, L_MSD-dt_: parameter comparison of the MSD-Δt plot analysis (Fig. 8a-c)

1. Click the “MSD-dt fit” button to output the MSD-Δt plots (Fig. 8a, mean ± sem of the cells) and the violin/box plot of the MSD at the frame that was selected the parameter panel (“Frame of D_avg_” in Fig. 3a). (*see* **Notes 32, 42 and 44**)
2. Click the “D (MSD-dt)” button to output the violin/box plot of the diffusion coefficient (D_0_) estimated from the initial slope of the MSD-Δt plots (Fig. 8b).
3. Click the “L (MSD-dt)” button to output the violin/box plot of the confinement length (L) estimated from the MSD-Δt plots (Fig. 8c).
4. Click “ANOVA1” button to output the result table of the one-way ANOVA test after selecting the violin/box plot.
5. Click “Dunnett-test” button to output the result table of the Dunnett test after selecting the violin/box plot, where the inverse-agonist LY341495 significantly elevates the D_0_ and L of mGluR3 to a statistically meaningful extent with A4 (100 nM) and A5 (1000 nM) when A1 (0 nM) set as the control group as shown in Figs. 8b and 8c.

#### 3.8.15. D_state_ and D_ratio_: parameter comparison of the displacement histogram analysis (Figs. 8d-k)

1. Click the “D (state)” button to output violin/box plots of the diffusion coefficient of each state (Fig. 8d-g) estimated from the displacement histogram analysis (*see* **Notes 33, 42 and 44**).
2. Click the “D_ratio_” button to output violin/box plots of the diffusion state ratio (Fig. 8h-k) estimated from the displacement histogram analysis (*see* **Notes 33, 42 and 44**).
3. Click “Tukey-test” button to output the result table of the Tukey-test after selecting the violin/box plot, where the LY341495 significantly elevate the change of the fast state fraction with A4 (100 nM), compared with A1 (0 nM), A2 (1 nM), and A3 (10 nM) in Fig 8k.

#### 3.8.16. Intensity: Output the intensity analysis results (Fig. 8l-n)

1. Click the “Intensity” button to output line-graphs of the oligomer size distribution (Fig. 8l), violin/box plots of the single-molecule intensity (Fig. 8m) and the mean oligomer size (Fig. 8n) estimated from the intensity histogram analysis (*see* **Notes 34, 42 and 44**).
2. Click “Dunnett-test” button to output the result table of the Dunnett test after selecting the violin/box plot, where the inverse agonist LY341495 induces statistically significant decrease of the mean oligomer size with A2 (1 nM), A3 (10 nM) and A5 (1000 nM) when A1 (0 nM) set as the control group in the Dunnett-test (Figs. 8n).

#### 3.8.17. Off-rate: Output mean decay curves of the duration of trajectories and violin/box plots of the off rate parameters. (Fig. 8o-s)

1. Click the “Off-rate” button to output mean decay curves of the duration of trajectories (Fig. 8o), violin/box plots of the time constant (Fig. 8p, q) and the fractions of “Off-rate” (Fig. 8r, s) estimated from the exponential fitting (*see* **Notes 36, 37, 42 and 44**).
2. Click “Tukey-test” button to output the result table of the Tukey-test after selecting the violin/box plot, where the inverse-agonist LY341495-induced-changes of the long time constant “Off-rate 2” of the decay curve are observed between A1 and A2, between A2 and A4, and between A2 and A5 by Tukey-test (Fig. 8q).

#### 3.8.18. On-rate: Output violin/box plot of the on rate parameters (Fig. 8t)

1. Click the “On-rate” button to output violin/box plot of the “On-rate” estimated from the exponential fitting (*see* **Notes 36, 38, 42 and 44**).
2. Click “ANOVA1” button to output the result table of the one-way ANOVA test after selecting the violin/box plot, where no significant change is observed by the one-way ANOVA.

#### 3.8.19. Density: Output violin/box plot of the density analysis parameters (Fig. 8u)

1. Click the “Density” button to output violin/box plots of the density and area estimated from the density analysis (*see* **Notes 35, 42 and 44**).
2. Click “ANOVA1” button to output the result table of the one-way ANOVA test after selecting the violin/box plot, where no significant change is observed by the one-way ANOVA.

#### 3.8.20. 2D plots: Output 2D plots of the diffusion, intensity, and density parameters of the single-cells (Fig. 8v)

1. Click the “2D plots” button to output scatter plots that compared between D_avg_ and mean oligomer size, between D_avg_ and density, and between mean oligomer size and density (*see* **Notes 32, 34, 35, and 45**).

#### 3.8.21. Total Basic Analysis of the HMM data format: (Fig. 9)

1. Set the optimized parameters in the parameter panel. The parameter settings are basically the same as the AAS and G-count data format (*see* Section 3.8.2).
2. Build the folder structure as shown in Fig. 3b and 7. In the case of “Total Basic Analysis”, the name of each folder (e.g. “A1” ~ “A5”) is entered as “Sample Name” in the macros automatically. Do not set too long folder names or folder names with space.
3. Check the input format checkbox in the “Measurements/SMT parameters” panel to fit the HMM data format (Fig. 9a) (*see* **Note 22**).
4. Click the “Total Basic Analysis” button (Fig. 7).
5. Select the folder containing the subfolders to be compared (e.g. Select the “LY34” folder containing “A1” ~ “A5” subfolders in Fig. 7).
6. Click the “select folder” button in the pop-up panel, where the “Total Basic Analysis” starts “Respective analysis” of each folder sequentially. (*see* Sections 3.8.22 ~ 3.8.25 regarding each macro that outputs different results from the Sections 3.8.3 ~ 3.8.11.)

**Fig. 9.**
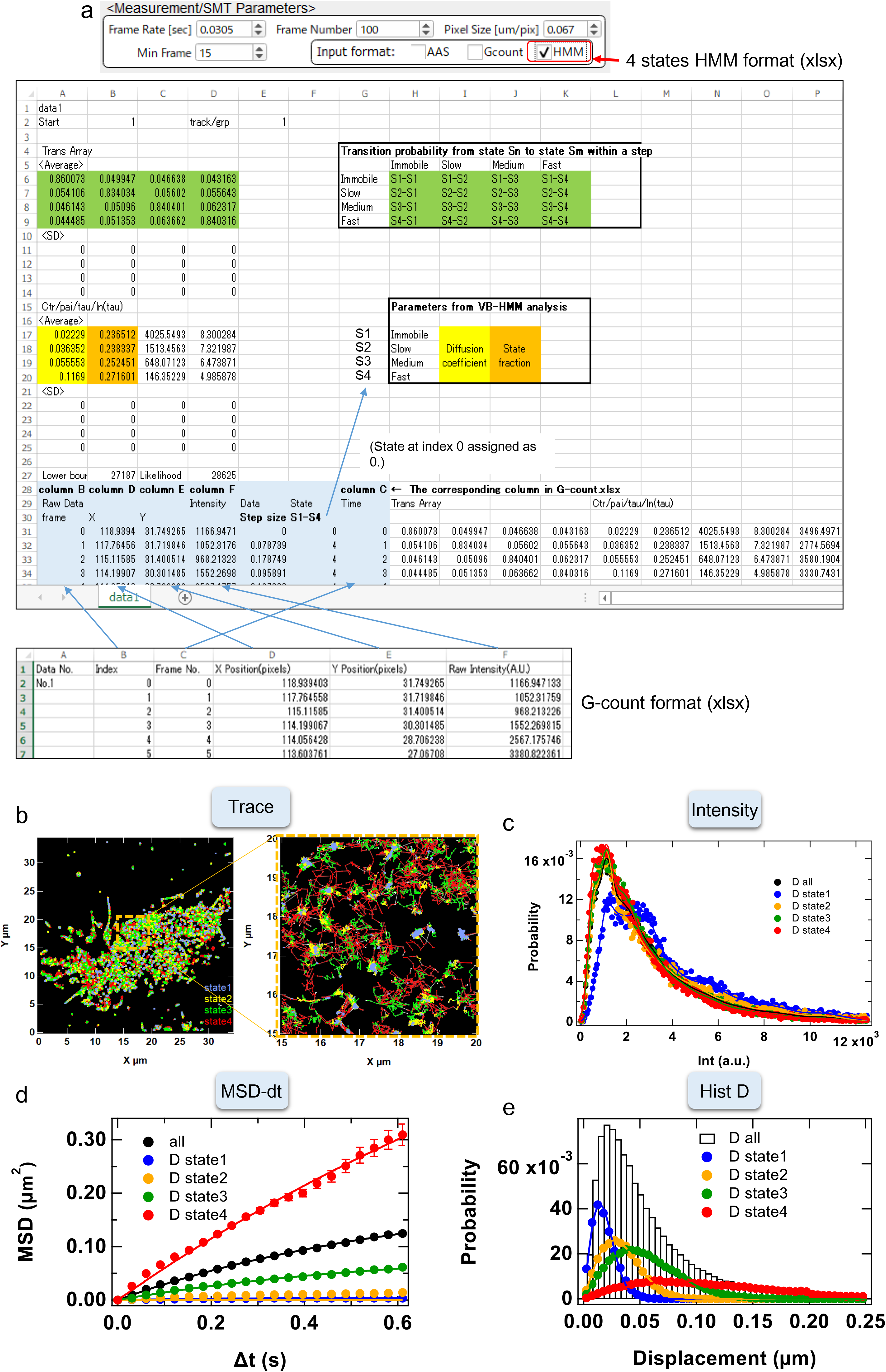
“Respective Analysis” of the HMM format data of mGluR3. (a) The 4 state HMM and G-count formats (xlsx files) were exemplified. (b) Trajectories of mGluR3 in the 4 diffusion states output by the “Trace” macro. (c) Intensity histograms of the total trajectories and the 4 diffusion states of mGluR3 output by the “Intensity” macro. (d) MSD-Δt plots of the total trajectories and the 4 diffusion states of mGluR3 output by the “MSD-dt” macro. (e) Displacement histograms of the total trajectories and the 4 diffusion states of mGluR3 output by the “Hist D” macro. The colors in (b-e) are as follows: [black (or white in b): total, blue: state1 (immobile), yellow: state2 (slow), green: state3 (medium), red: state4 (fast)]

#### 3.8.22. Trace: Output XY plots of the trajectories (Fig. 9b)

1. Click the “Trace” button (Fig. 3a).
2. Input the “Sample Name” corresponding a folder name under the root folder in the pop-up panel.
3. Click the “continue” button, where the macro automatically outputs graphs of the trajectories of each diffusion state (Total: Yum_S0 vs Xum_S0, Immobile: Yum_S1 vs Xum_S1, Slow: Yum_S2 vs Xum_S2, Medium: Yum_S3 vs Xum_S3, Fast: Yum_S4 vs Xum_S4) in each “Folder Name” folder in different colors (Total: white, S1: blue, S2: yellow, S3: green, S4: red).
4. Select a ROI in the trace graph and right click to select “Expand” to visualize trajectories in the expanded area.

#### 3.8.23. Intensity: Output intensity histograms (Fig. 9c)

1. Click the “Intensity” button after setting the parameters in the parameter panel (Fig. 3a).
2. Input the “Sample Name” that corresponding a folder name under the root folder in the popup panel.
3. Click the “continue” button, where the macro automatically outputs graphs of the intensity histograms of the total trajectories (D all (black)), and of 4 diffusion states of HMM analysis (Immobile: D state 1 (blue), Slow: D state 2 (yellow), Medium: D state 3 (green), Fast: D state 4 (red)). (*see* **Note 34**).

#### 3.8.24. MSD-dt: Output MSD-Δt plots (Fig. 9d)

1. Click the “MSD-dt” button after setting the parameters in the parameter panel (Fig. 3a).
2. Input the “Sample Name” that corresponding a folder name under the root folder in the popup panel.
3. Click the “continue” button, where the macro automatically outputs graphs of the MSD-Δt plots of 4 diffusion state of HMM analysis (Immobile: D state 1 in blue, Slow: D state 2 in yellow, Medium: D state 3 in green, Fast: D state 4 in red) in addition to the MSD-Dt plot of the total trajectories (all: black). (*see* **Note 32**).

#### 3.8.25. Hist D: Output displacement histograms (Fig. 9e)

1. Click the “Hist D” button after setting the parameters in the parameter panel (Fig. 7).
2. Input the “Sample Name” that corresponding a folder name under the root folder in the popup panel.
3. Click the “continue” button, where the macro automatically outputs graphs of the displacement histograms of the total trajectories (D all (black)), and of four diffusion states of HMM analysis (Immobile: D state 1 (blue), Slow: D state 2 (yellow), Medium: D state 3 (green), Fast: D state 4 (red)). (*see* **Note 33**).

#### 3.8.26. Comparison Analysis of the HMM data formats: Total comparison analysis, and multiple comparison tests are carried out (Fig. 7 and 10)

1. Perform more than twice of “Respective Analysis” before start “Comparison Analysis”. (*see* **Note 29**)
2. Click the “Comparison Analysis” button to start the macros sequentially. (*see* **Note 46**) (see Sections 3.8.27 ~ 3.8.29 regarding each macro that outputs different results from the Sections 3.8.14 ~ 3.8.20.)
3. Click the multiple comparison test buttons (*see* Section 3.8.13) to perform one-way ANOVA and multiple comparison tests among groups on a violin/box plot by clicking the corresponding button after selecting a violin/box plot. (*see* **Note 43**)

**Fig. 10.**
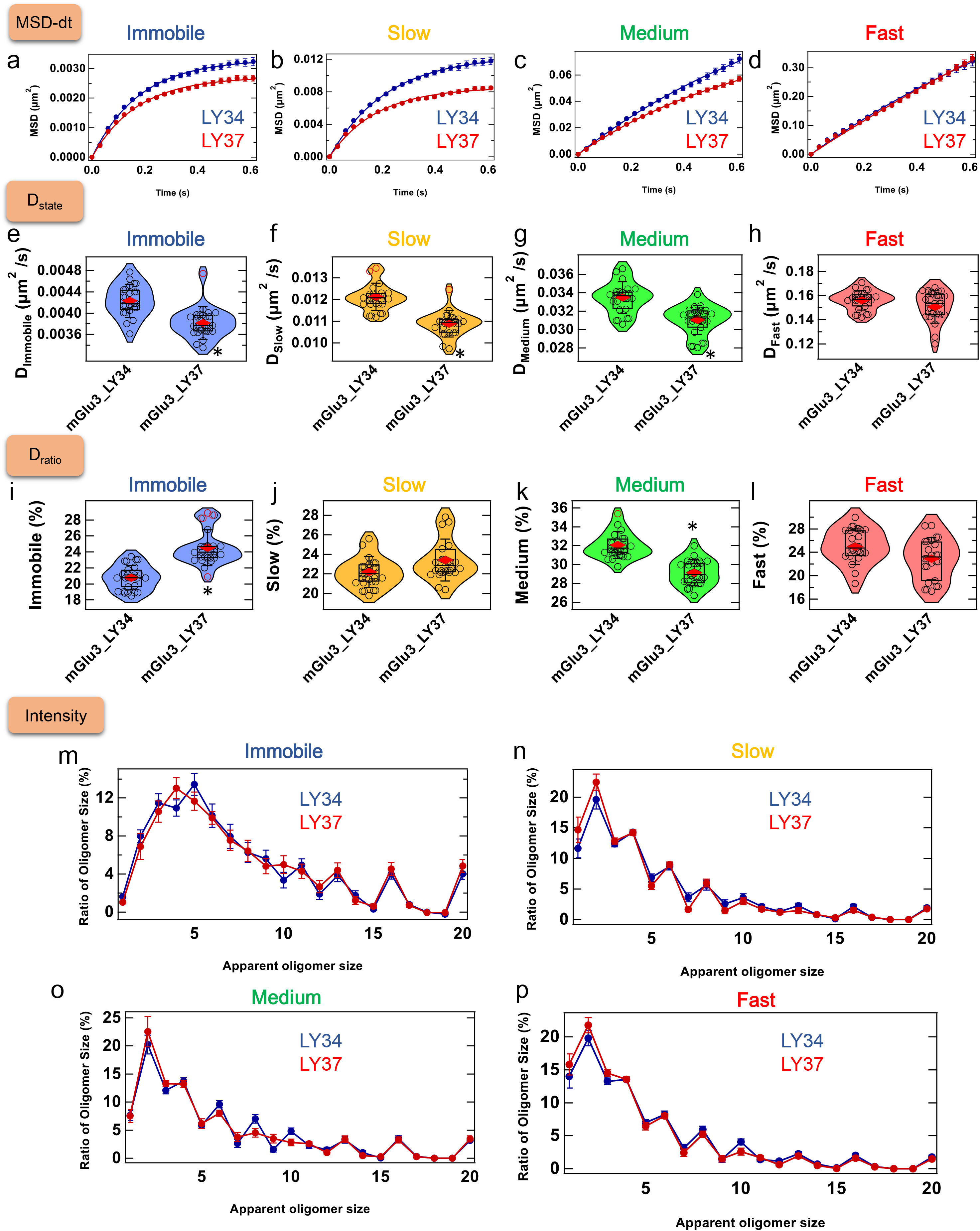
“Comparison Analysis” to compare the parameters of mGluR3 in two different ligand conditions (HMM format data). The ligand conditions are “mGlu3_LY34”: 100 nM LY341495 and “mGlu3_LY37: 10 μM LY379268. These data correspond to the red channel data of the dual-color imaging with EGFP-labeled CLC (see Fig. 1, 11, and 12). (a-d) Example results of “MSD-dt” plot analysis macro. (a: immobile, b: slow, c: medium, d: fast) (e-h) Example results of “D_state_” macro (e: immobile, f: slow, g: medium, h: fast). (i-l) Example results of “D_ratio_” macro. (i: immobile, j: slow, k: medium, l: fast). (m-p) Example results of “Intensity” macro (i: immobile, j: slow, k: medium, l: fast). *p < 0.05 by the Welch’s t-test (two-tailed) in all the panels.

#### 3.8.27. MSD-dt: parameter comparison of the MSD-Δt plot analysis (Fig. 10a-d)

1. Click the “MSD-dt fit” button to output the MSD-Δt plots (Fig. 10a-d, mean ± sem of the cells), where the MSD of mGluR3 molecules in the immobile (a), slow (b), and medium (c) states were decreased upon activation. (*see* **Notes 32, 42, 44**)

#### 3.8.28. D_state_ and D_ratio_: parameter comparison of the VB-HMM analysis (Fig. 10e-l)

1. Click the “D (state)” button to output a category plot and violin/box plots of the diffusion coefficient of each state estimated from the VB-HMM analysis (*see* **Notes 22, 42, 44 and 47**).
2. Click the “D_ratio_” button to output a category plot and violin/box plots of the diffusion state ratio estimated from the VB-HMM analysis (*see* **Notes 22, 42, 44, and 48**).

#### 3.8.29. Intensity: Output the intensity analysis results. (Fig. 10m-p)

1. Click the “Intensity” button to output line-graphs of the oligomer size distribution of each diffusion state (Fig. 8d), where the apparent oligomer size distribution was normalized with the total number of oligomers in each state (*see* **Notes 34 and 49**).

#### 3.8.30. Colocalization Analysis of the HMM data formats: (Fig. 11)

1. Perform “Respective Analysis” of the HMM data format to analyze the data for the corresponding two channels before start “Colocalization Analysis”. (*see* **Notes 29 and 50**).
2. Set parameters in the parameter panel to define colocalization (Fig. 11a) (*see* **Note 51**).
3. Click “Colocalization Analysis” button to start.
4. Input “SampleName1” (e.g. “mGlu3_LY34”) and “SampleName2” (e.g. “CLC_LY34”) that corresponding the folder names under the root folder to be analyzed colocalization in the pop-up panel.
5. Click the “continue” button where, the “Colocalization Analysis” automatically starts all the macros below sequentially. Find Col. → Hist D → Off-rate → On-rate → D ratio →stats (*see* Sections 3.8.31 ~ 3.8.35 regarding each macro.)

**Fig. 11.**
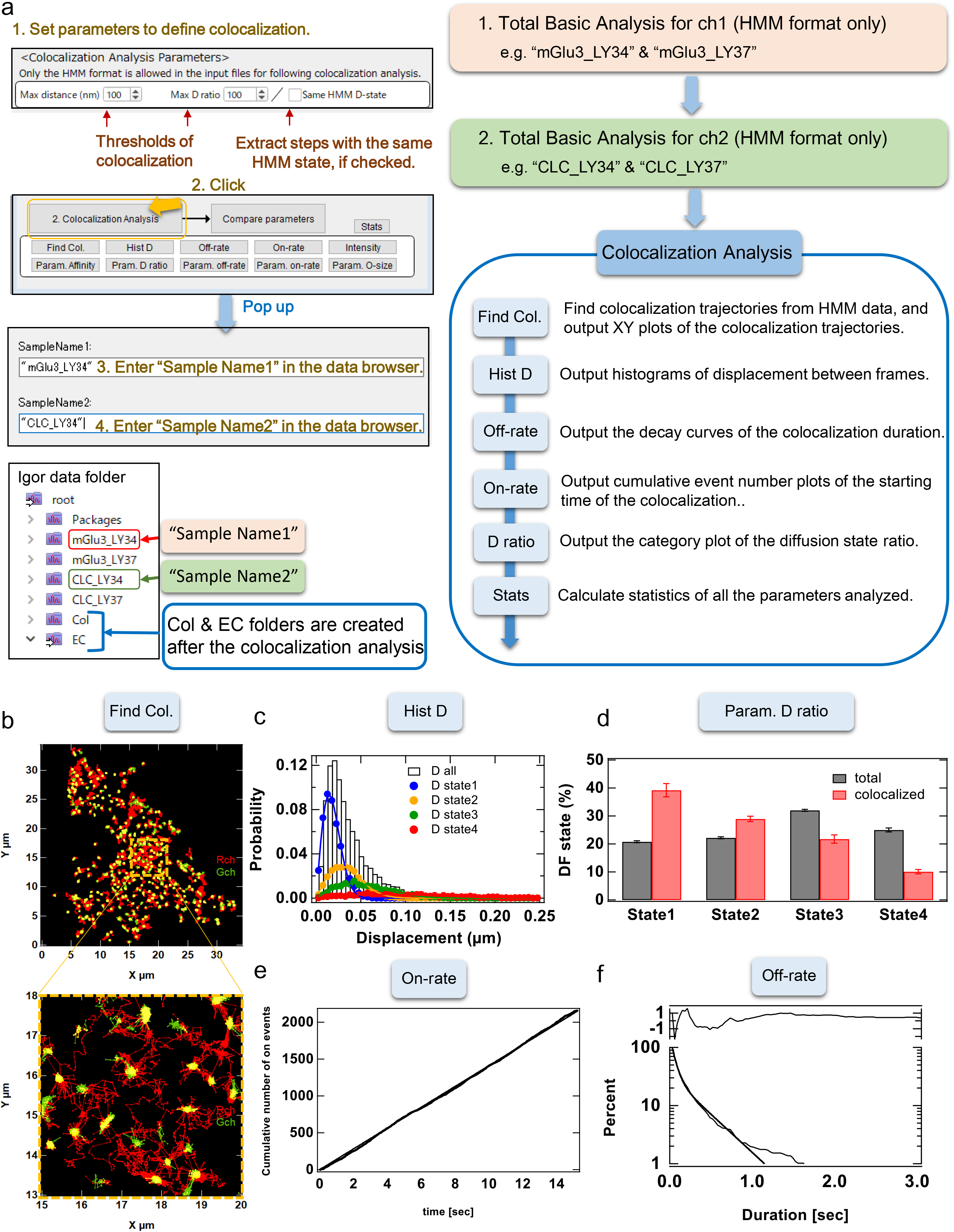
Guidance on “Colocalization Analysis” between TMR-labeled mGluR3 and EGFP-labeled CLC. (a) The work flow of “Colocalization Analysis”. (b) Example results of “Find Col.”. Trajectories with more than one colocalize step (yellow) were plotted (red: mGluR3, green: CLC). (c) Example results of “Hist D” macro for the colocalize steps. (d) Example results of “Parm. D ratio” macro. The diffusion state fractions of the total trajectories (gray) and the colocalize steps (red) of mGluR3 are shown in the category plots (mean ± sem of the cells). (e) Example result of “On-rate” macro. A cumulative event number plot of the association between mGluR3 and CLC. (f) Example result of “Off-rate” macro. A decay curve of the colocalization duration were fitted with the double-exponential function.

#### 3.8.31. Find Col.: Find colocalization trajectories from HMM data (Fig. 11a, b)

1. Click “Find Col.” button to start.
2. Input “SampleName1” and “SampleName2” that corresponding the folder names under the root folder to be analyzed colocalization in the pop-up panel.
3. Click the “continue” button, where the macro automatically finds colocalization trajectories from HMM data, and outputs XY plots of the colocalization trajectories (Fig. 11b, *see* **Note 52**).

#### 3.8.32. Hist D: Output histograms of displacement between frames. (Fig. 11c)

1. Click “Hist D” button to start.
2. Input “SampleName1” and “SampleName2” that analyzed in advance by “Find Col” in the pop-up panel.
3. Click the “continue” button, where the macro automatically outputs histograms of the displacement between frames of the total colocalization trajectories and that of the 4 diffusion states of HMM analysis (Fig. 11c, *see* **Note 53**).

#### 3.8.33. Param. D ratio: Output the category plot of the diffusion state ratio (Fig. 11d)

1. Click “Param. D ratio” button to start.
2. Input “SampleName1” and “SampleName2” that analyzed in advance by “Find Col” in the pop-up panel.
3. Click the “continue” button, where the macro automatically outputs the category plot that compared the diffusion state fractions of the total trajectories and of the colocalization frames in each channel (Fig. 11d, mean ± sem of the cells).

#### 3.8.34. On-rate: Output cumulative event number plots of the starting time of the colocalization. (Fig. 11e)

1. Click “On-rate” button to start.
2. Input “SampleName1” and “SampleName2” that analyzed in advance by “Find Col” in the pop-up panel.
3. Click the “continue” button, where the macro automatically outputs the cumulative event number plots of the starting time of trajectories (Fig. 11e, *see* **Note 54**)

#### 3.8.35. Off-rate: Output the decay curves of the colocalization duration (Fig. 11f)

1. Click “Off-rate” button to start.
2. Input “SampleName1” and “SampleName2” that analyzed in advance by “Find Col” in the pop-up panel.
3. Click the “continue” button, where the macro automatically outputs the decay curve of the colocalization duration (Fig. 11f, *see* **Note 55**).

#### 3.8.36. Param. Off-rate: Output the category plots of the parameters of the off-rate analysis (*see* Note 56)

1. Click “Param. Off-rate” button to start.
2. Input “SampleName1” and “SampleName2” that analyzed in advance by “Find Col” in the pop-up panel.
3. Click the “continue” button, where the macro automatically outputs the category plots of the parameters of the off-rate analysis (mean ± sem of the cells).

#### 3.8.37. Intensity: Output the intensity histograms at the colocalization frames (*see* Note 56)

1. Click “Intensity” button to start.
2. Input “SampleName1” and “SampleName2” that analyzed in advance by “Find Col” in the pop-up panel.
3. Click the “continue” button, where the macro automatically outputs the intensity histograms of the colocalization frames of the each channel. (*see* **Note 57**)

#### 3.8.38. Compare Parameters of the Colocalization Analysis: (Fig. 12)

1. Perform more than twice of “Colocalization Analysis” before start “Compare Parameters”. The macro compares the parameters among those in the subfolders in the “EC” folder.
2. Remove the subfolders that you do not compare from the “EC” folder since the subfolders of the both channels exist in the EC folder just after the “Colocalization Analysis”. (*see* **Note 58**) Import data from other Igor files by using “Brows Expt.” in the Igor data browser if you compare the data analyzed by the “Colocalization Analysis” in the smDynamicsAnalyzer beforehand. Select the Igor file, and drag and drop the data folders in the “EC” folder from the browsing file to the current file.
3. Click the “Compare Parameters” button to start (*see* **Note 59**), where the macro automatically outputs the violin/box plots of all the parameters below sequentially. P_col_ (*see* **Note 60**) → On-rate (*see* **Note 61**) → Off-rate (*see* **Note 62**) → D ratio (*see* **Note 63**)
4. Click the multiple comparison test buttons (*see* 3.8.4) to perform one-way ANOVA and multiple comparison tests among groups on a violin/box plot by clicking the corresponding button after selecting a violin/box plot (*see* **Note 43**).

**Fig. 12.**
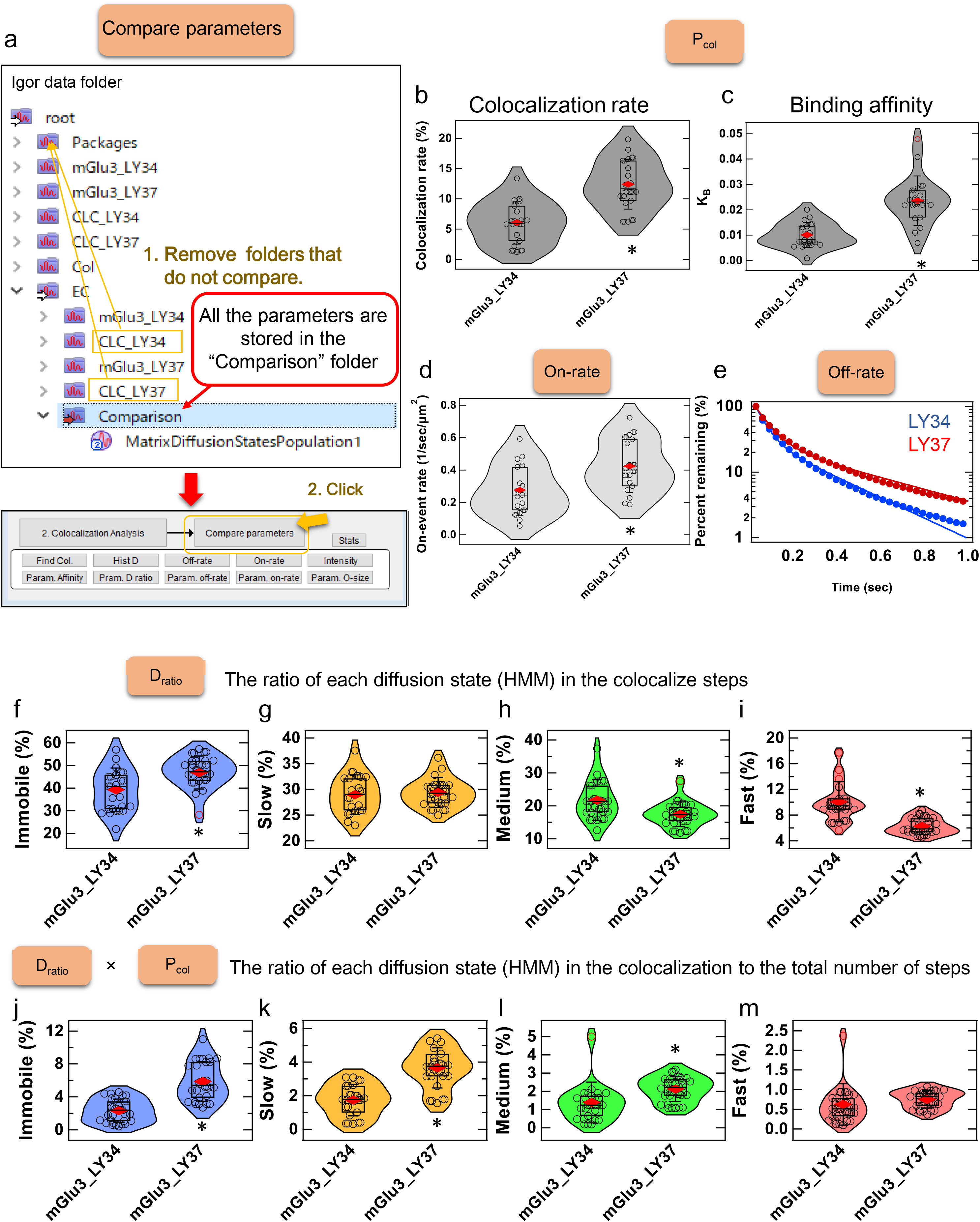
Guidance on “Compare Parameters” to compare the parameters of the colocalization between mGluR3 and CLC in two different ligand conditions. The ligand conditions are “mGlu3_LY34”: 100 nM LY341495 and “mGlu3_LY37: 10 μM LY379268. (a) How to use “Compare Parameters” macro that integrates all the macros that output all the panels (b-m). Example results of “Pcol” macro (b, c). The violin/box plots of the colocalization rate (b) and the binding affinity KB (c). (d) The violin/box plots that output by the “On-rate” macro. (e) Example results of “Off-rate” macro. The decay curves the colocalization duration were compared (blue: 100 nM LY341495, red: 10 μM LY379268). Example results of “D_ratio_” (f-i) and “D_ratio_ × Pcol” (j-m) macros. The violin/box plots of the immobile (f, j), slow (g, k), medium (h, l), and fast (i, m).*p < 0.05 by the Welch’s t-test (two-tailed) in all the panels. See **Note 37** for the violin/box plot settings.

## 4. Notes

1. Listed below is the mammalian expression vectors containing HaloTag/SNAP-tag/GFP-tag which are commercially available. In the case of the HaloTag, Promega provides vectors with low level expression promoters, instead of the CMV promoter. These vectors are useful for the SMI. We recommend pFC15 (CMVd1) vector for the first choice for GPCR expression. Expression of the N-terminally HaloTag-fusion GPCR in pFN21A vector, which can be purchased from Promega as Flexi ORF Clone, often caused a N-terminal half-fragment of HaloTag-fusion protein in the cytoplasmic region (In our experience, ADRB2, HTR2A, ADORA2A are well expressed, but CXCR4, FFAR4, F2R, GCGR, GRM3 are not.) In the case of Gα-proteins, the tags should be inserted into the loop region as previously reported [4, 16].

i. HaloTag vectors (Promega) pFC14A/K (CMV), pFC15A/K (CMVd1), pFC16A/K (CMVd2), pFC17A/K (CMVd3) for C-terminal fusion pFN21A/K (CMV), pFN22A/K (CMVd1), pFN23A/K (CMVd2), pFN24A/K (CMVd3) for N-terminal fusion
ii. SNAP-tag vector (NEB) pSNAPf Vector
iii. GFP-tag vector (Clonetech) pEGFP-N1 Fluorescent proteins including mEGFP can be used for SMI, but the brightness and the photostability are much lower than FL dyes.
2. Any adherent cells are applicable for the present study (e. g. HEK293, HeLa, A549, CHO, MEF, MDF cells). In general, the stronger cell adhesion to glass bottom is better for TIRFM. Here we exemplify HEK293 cell lines, whose adhesive strength is not so high but are used frequently for pharmacological studies of GPCRs. HEK293A cell line is better than other variants such as HEK293S and HEK293T cell lines in the view point of cell adhesion.
3. In our measurement settings, the HaloTag JF549 and SF650 ligands are the best membrane permeable fluorophores for the Cy3 and Cy5 channels, respectively, with respect to brightness, photostability, and affinity to the tags. The affinities of SNAP-tag ligands are one-order lower than those of HaloTag ligands [4]. The SNAP-Cell ligands are useful for the dual-color SMI with HaloTag.
4. Glass-bottom dishs similarly work, but the washed coverslips are much recommended to reduce the FL from a dust and the non-specific binding of FL dyes.
5. The required laser power depends on the optical system including the irradiation area. The following is the examples of lasers (Coherent) used in our laboratory. Sapphire 488-200: 488 nm, 200 mW laser for GFP. Compass 315M-100: 532 nm, 100 mW laser for TMR or JF549. OBIS 561 nm: 100 mW laser for TMR or JF549. OBIS 637 nm: 140 mW laser for SF650 or SNAP-647-SiR.
6. Optics for TIRF illumination is provided by each microscope company (e. g. Ti2-LAPP, Nikon, and cellTIRF-4Line, Olympus), but homemade optics are preferred with respect to customization (Fig. 1). One can achieve uniform illumination of the laser by the rotating beam illumination using xy galvano scanner controlled by a function generator (*see* **Note 13**).
7. The optimal filter set depends on the optical properties of the lasers and the fluorophores. Here we describe an example filter set for dual-color imaging.

A. Dual-color imaging with 488 nm and 561 nm lasers Dichroic mirror in the microscope: ZT488/561rpc (Chroma) Dichroic mirror in the two-channel imaging system: T560lpxr (Chroma) Emission filters: ET525/50m (Chroma) for GFP ET605/70m (Chroma) for TMR or JF549
B. Dual-color imaging with 532 nm and 637 nm lasers Dichroic mirror in the microscope: ZT532/640rpc (Chroma) Dichroic mirror in the two-channel imaging system: FF640-FDi01 (Semrock) Emission filters: FF01-585/40 (Semrock) for TMR or JF549 FF01-676/29 (Semrock) for SF650 or SNAP-647-SiR. An excitation filter (e.g. ZET642/20x, Chroma) is required for OBIS 637 nm to cut off the short wavelength light that is leaked from the laser.
8. Any transfection methods may work. However, it is difficult to achieve an appropriate level of expression of the fluorophore-labeled GPCRs in the PM (less than ~1/μm^2^) if one follows the manufacturer’s protocols, because most of the protocols are customized for overexpression. We can achieve relatively higher transfection efficiency and less expression levels than others through transfecting the cells with a low amount of pDNA per Lipofectamine 3000 level. If the expression level is still high with this protocol, one may further reduce the amount of pDNA in the transfection reagent, and refresh the medium 3 h after transfection. The plasmid vectors with CMVd2 or CMVd3 promoter are also useful for reducing the expression level.
9. One may exert a dual-color imaging simply using a mixture of two kinds of pDNA vectors.
10. P3000 has to be added at the end so as to inhibit the aggregation with ethylenediaminetetraacetic acid (EDTA).
11. The concentration of HaloTag/SNAP-tag ligands in cell staining depends on the purpose of the measurement. Higher the staining rate is recommended in order to quantify the oligomer size distribution. However, the ligand concentration levels recommended by the manufacturers (1~5 μM) are generally too high to distinguish the specific staining from non-specific ones. We previously characterized the concentration-dependent non-specific binding of HaloTag/SNAP-tag ligands, and we recommend to set the ligand concentration at most 300 nM [4]. Bosch et al. also demonstrated that the non-specific binding of 400 nM SNAP-tag ligand highly depends on a physical property of the fluorophores. Actually, many fluorophores cannot be used because of the severe non-specific binding to the cellular components and/or the glass surface [20]. In case one simply quantifies the diffusion dynamics of GPCRs, we recommend the use of 30 nM HaloTag ligands, where the staining rate are preferred to set about 70% and 20% for SF650 and TMR ligands, respectively [4]. The staining and wash steps with medium B containing 10% FBS reduce the non-specific binding. The longer incubation time in step 4 causes the higher non-specific binding of the dyes. The step 6 also reduces the non-specific binding due to reduced free fluorophores trapped on the outside of the coverslip.
12. When using Metamorph, the split view (Display Menu - Split View - Dialog Box Options - Align Tab) is useful to adjust two channels.
13. Keep in mind the following key points upon adjustment of the TIRF optics (Fig. 1).

i. Set the focus on the fluorescent bead sample on a glass-bottom dish, and adjust the optical system so that the parallel light passes straight up through the objective lens when the collimated laser beam is injected into the tube lens of the microscope (epi-illumination).
ii. In the case of the Nikon TiE or Ti2E, the position of the laser beam at the back focal plane of objective lens can be checked by inserting a belt run lens in the lens-barrel (upper image in Fig. 1). In the case of Olympus IX81 or IX83, centering telescope is useful. Adjust the angle of the mirror at the image conjugate position so that the laser beam passes outside of NA1.33 at the objective aperture (cyan lines of Fig. 1).
iii. In order to achieve uniform illumination, we have set up a rotating beam illumination system by placing XY galvanometer scanners in the image conjugate positions and sending 90-degree phase-shifted sine waves from the function generator (yellow lines of Fig. 1). This system enables us to switch the illumination modes among epi, oblique and TIRF by changing the angle of the mirrors by voltage control from PC.
14. When using Metamorph, “Multi Dimensional Acquisition” mode is useful. Metamorph can remember XY positions of the cells, which enables us to do the time-lapse stream acquisition of the same cells by using a home-made journal.
15. The acceptable image format depends on the SMT analysis software (e.g. G-count: 8-bit avi, AAS: 16-bit or 8-bit tif). To keep the single-molecule intensity constant across the images, dynamic range of the signal intensity should be kept constant among movies and frames in a movie (e.g. minimum: 0, maximum: 1000 in 16-bit tif images) followed by image conversion from 16-bit to 8-bit. Be careful not to make a saturated pixel (255 in 8-bit image), which is produced if the maximum intensity value in the conversion is lower than the maximum brightness in 16-bit images.
16. “BatchImageProcessingSMT.ijm” is a macro file for batch processing of the series of processes in section 3.5. to all the multiple tif files within a single folder. One may freely customize and use the macro with suitable citations of this chapter. Install Running_ZProjector and GridAligner plugins before using the BatchImageProcessingSMT. One can download the Plugins from Vale Lab homepage: http://valelab.ucsf.edu/~nstuurman/ijplugins/
17. In case of the users of the GridAligner plugin to align two channels, this process is necessary beforehand. This process may be skipped for the users of the AAS, although the macro “Composite2ch” is useful for the movie presentation of the dual color image. The users can change the color of each channel of the composite image by ImageJ through following the path: Image > Color > Arrange Channels.
18. The GridAligner automatically generates the image with the suffix “aligned” after affine transformation. This process is time consuming because the GridAligner does not support the batch processing. We recommend an alternative affine transformation program in a shareware package for the SMI analysis: Auto Analysis System (AAS), Zido (https://zido.co.jp/en/).
19. We previously used G-count software (G-angstrom) for SMT analysis [4], but the G-count does not support the batch processing. One may use the AAS (Zido, https://zido.co.jp/en/) too for SMT analysis with a similar algorism to G-count. The VB-HMM analysis can be performed with a LabView-based homemade program as previously described in detail [4]. The VB-HMM analysis enables to assign a diffusion state to each step of the trajectories. The clustering algorism is based on the following assumptions: (i) the diffusion states of molecules are in a steady state among several states with different diffusion coefficients, (ii) the transition from one state to the other follows a Markov process. The number of the diffusion states can be estimated from a comparison of the lower bound values of the likelihood that calculated based on the VB expectation-maximization algorithm [4]. Here we provide an example of raw data after VB-HMM analysis of TMR-labeled mGluR3 and GFP-labeled clathrin light chain (CLC). (https://github.com/masataka-yanagawa/IgorPro8-smDynamicsAnalyzer). The latest AAS after version 2.4.5 incorporates the VB-HMM analysis option. The vbSPT is a similar program that is available as an open source software working in MATLAB (http://vbspt.sourceforge.net/) [21]. The parameter settings in the AAS are as follows:

i. Parameters for single-molecule detection within a frame of the multiple tif image. As shown in Fig. 2b, the image of each frame is scanned with a square ROI (N × N pix) sliding with a scan size (M pix). In each ROI, the following “2D Gaussian function on a primary plane” is used to fit and detect the bright spots with an intensity over the threshold (I_min_).

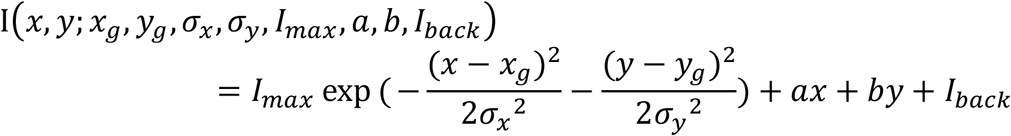

where (x_g_ y_g_) is the centroid of a bright spot that have a peak intensity I_max_. The 2D Gaussian fitting can be done by the Levenberg-Marquardt method. The bright spot is assumed to be on an inclined plane described by ax+by+I_back_. We found that M should be lower than N/4 to reduce detection errors at ROI boundaries. I_min_ depends on the experimental and image processing conditions. Increase the I_min_ if the noise spots are over detected. Decrease the I_min_ if the bright spots, which can be detected by eye, are not detected.
ii. Parameters for connecting coordinates of a spot over frames to make trajectories. When two spots detected in successive frames are within the “connection distance (D)” and within the “connection longest frame (L)”, they are connected as a trajectory of a singlemolecule. If there are more than two spots meeting the criteria above, the closest spots are connected. If the two spots in frame i and frame i+L are connected as a trajectory, and if there are no spots corresponding the trajectory in frames between i and i+L, the localization and the intensity of the spot in frames between i and i+L are linearly complemented by the data of spots in the frame i and frame i+L. The trajectories shorter than minimum trajectory length (m) are deleted to reduce the false positive detection of noise spots. e.g.) connection longest frame (L): 2 frames connection distance (D): 8 pix trace minimum frame number (m): 3 frames (over 15 frames are recommended for VB-HMM analysis) The higher L and D values are set, the more misconnected the trajectories of different molecules. In contrast, the lower L and D values are set, the less detected the trajectories with a high diffusion coefficient. AAS can output the following files when the check boxes for the required items are checked:

i. Trajectory files: the csv (Fig. 2b) and/or binary files of trajectories
ii. Image files to check the result: 8-bit tif files with ROIs on the spots detected (Fig. 2b).
iii. Statistics of each multiple tif file:

MSD within frames (Δt = frame rate*frames)
Displacement histogram within a frame time
Intensity histogram of all the spots We do not output (iii) because the more advanced analysis can be performed by using homemade program “smDynamicsAnalyzer” in Igor Pro 8.0 as a platform (*See* 3.7). Many SMT programs in various platforms are available now as alternatives to the AAS and G-count (e.g. TrackMate in ImageJ/Fiji, Spot Tracking in Icy, SMTracker in MatLab). Every programs have their own unique merits. Chenouard et al. compared the performances of the 14 methods of the SMT analysis to simulation data with various signal-to-noise (S/N) ratios [17]. The performance of each program is influenced by the S/N ratios of the simulation data. Moreover, noise properties in the experimental data are even more complicated than ones in the simulation data. No matter what programs, it is important to make sure that the output trajectories from a sample movie match the trajectories of the spots followed by your naked eyes. If there is an obvious difference between them, consider changing the parameters or the program to use. We considers that the differences in the diffusion dynamics of GPCRs proposed in the previous reports [4, 11–16] may be partly due to the differences in these parameters or the algorisms of SMT analysis. The comprehensive SMI analysis of GPCRs by the same platform is required to reveal the similarity and diversity of the dynamics of GPCRs. The advantages of AAS are that it supports batch processing and parallel computing modes to speed up the SMT analysis, which is even advantageous for handling such large data.
20. One may freely customize and use the macro with suitable citations of this chapter. For more detailed instructions, refer to the supplementary pdf file uploaded to the following URL (https://github.com/masataka-yanagawa/IgorPro8-smDynamicsAnalyzer).
21. In the “Measurement/SMT Parameters” section, input the proper values regarding the measurement setting and SMT analysis. (e.g. Set “Frame rate [sec]” as 0.0305 when the exposure time was 30.5 ms. Set “Frame number” as 100 when the files analyzed by SMT were 100 frames. Set “Pixel Size [um/pix] as 0.067 when the pixel size was 67 nm/pixel. Set “Min frame” as 15 when the minimum frame of trajectories in the SMT analysis was 15 frames, which corresponds to “trace minimum frame number” in the AAS.) For other sections, a parameter optimization by using test data is required to get proper fitting results before starting “Total Basic Analysis” (*see* Section 3.8.21).
22. Three input formats are available (AAS.csv, G-count.xlsx, and HMM.xlsx). If you use other SMT or clustering program, refer to the sample files (https://github.com/masataka-yanagawa/IgorPro8-smDynamicsAnalyzer), and modify the file format to one of the formats. The smDynamicsAnalyzer loads information about the time point of spots detected [frame] (column B in AAS.csv, column C in G-count.xlsx, and column G in HMM.xlsx), the X-coordinate [pix] (column C in AAS.csv, column D in G-count.xlsx, and column B in HMM.xlsx), the Y-coordinate [pix] (column E in AAS.csv, column F in G-count.xlsx, and column C in HMM.xlsx), and the Raw Intensity after subtracting the background plane (column K in AAS.csv, column F in G-count.xlsx, and column D in HMM.xlsx) (Fig. 3b and Fig. 9a). In addition, the smDynamicsAnalyzer loads the diffusion state parameters estimated form the VB-HMM clustering analysis. The transition array highlighted by green is related to the probability of the state transition within a step. The area highlighted by yellow and orange corresponds to the diffusion coefficient [μm^2^/s] and the fraction of each state, respectively (Fig. 9a).
23. The “Total Basic Analysis” starts “Respective analysis” of each folder (e.g. A1 to A5) in the selected folder (e.g.) “LY34” folder) followed by “Comparison Analysis”. (see Sections 3.8.13 ~ 3.8.20) The experimental file is automatically saved after “Respective analysis” and “Comparison Analysis”. Note that the demo version of Igor is not able to save, so an error will appear.
24. You can skip the parameter optimization process if you do not use the item, although you will get an error message.
25. Refer to the Fig. S1 in the supplementary pdf file uploaded to the following URL as well (https://github.com/masataka-yanagawa/IgorPro8-smDynamicsAnalyzer).The filename of the csv/xlsx file is irrelevant to the “Folder Name” in Igor.
26. Push “Ctrl + B” to open the Igor data browser. You can delete an item by right click and select “delete object”. You cannot delete a folder from the data browser if you are opening tables or graphs containing a wave in the folder.
27. Push “Ctrl + 8” to close all the tables in Igor before you delete the folder.
28. Push “Ctrl + 9” to close all the graphs in Igor before you delete the folder.
29. To import data from other Igor files that are analyzed by the smDynamicsAnalyzer beforehand, click “Brows Expt.” button in the Igor data browser and select the Igor file. You can drag and drop the data folders from the browsing file to the current file.
30. Push “Ctrl + D” to duplicate a selected item.
31. Refer to the Figs. S2-3 in the supplementary pdf file uploaded to the following URL as well (https://github.com/masataka-yanagawa/IgorPro8-smDynamicsAnalyzer). This macro assumes 512×512 pix images. Change the display range by using “Modify Axis” panel, which is pop-up after double-clicking the X or Y axis of a graph, if required.
32. The parameter settings for the MSD-Δt plot analysis are as below. Refer to Figs. S4-6 in the supplementary pdf file uploaded to the following URL as well (https://github.com/masataka-yanagawa/IgorPro8-smDynamicsAnalyzer).

i. “Range [frames]” and “Threshold [%]” in Fig. 3a determine the maximum value of Δt.
ii. If the checkbox “Time Average” in Fig. 3a is checked, the macro calculates the time- and ensemble-averaged MSD of all the trajectories in each “Folder Name” folder; otherwise, the macro calculate the ensemble-averaged MSD. The time-averaged MSD within time nΔt of each trajectory is calculated by following function;

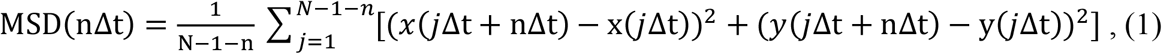

where n is the length of frames, Δt is the frame rate, and N is the total frame number of the trajectory. The non-time-averaged MSD within time nΔt of each trajectory is calculated by following function;

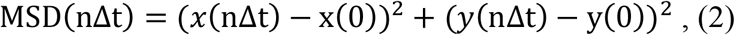 Then, the ensemble-averaged MSD (<MSD>) is calculated by following function:

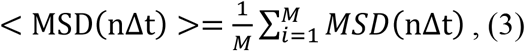

where M is the total number of trajectories.
iii. “Frame of D_avg_” in Fig. 3a specifies the point of Δt to calculate the MSD and D_avg_ to be compared in “Comparison Analysis” (*see* Sections 3.8.13, 3.8.14, and 3.8.20). The D_avg_ is calculated based on the two-dimensional diffusion equation as follows:

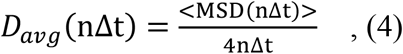
iv. The MSD-Δt plots are fitted using the following two equations.

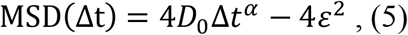

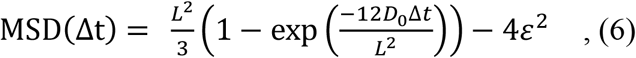 Only the fitting results of equation 6 are used in “Comparative Analysis”. The fitting results of equation 5 can be checked in the “fit_D_Alph_m” wave in the “Matrix” folder in each “Folder Name” folder.
v. If the fitting results are of less quality, change the initial parameters (D_0_, α, L, ε), and click the “MSD-dt” button again to reanalyze them. Repeat it until there are no more errors messages.
vi. D_0_ is related to the initial slope of the MSD-Δt plot. (4D_0_ is the initial slope of equation 1 when α = 1, and of equation 5),
vii. α is an index reflecting a diffusion mode. If 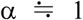, the molecules exhibit the simple Brownian diffusion in the average. If α > 1, the molecules exhibit the directed diffusion mode in the average. If α < 1, the molecules exhibit the confined diffusion mode in the average.
viii. L is the confinement length of the confined diffusion.
ix. ε is an error term that depends on the positional accuracy of the SMT analysis. If the checkbox “Fix ε” is checked, the input value is calculated as a fixed constant in the curve fitting; otherwise, the input value is calculated as an initial value.
33. The parameter settings for the displacement histogram analysis are as below. Refer to Figs. S7-9 in the supplementary pdf file uploaded to the following URL as well (https://github.com/masataka-yanagawa/IgorPro8-smDynamicsAnalyzer).

i. “Bin [um]” and “Dim” determines the X-axis range of the displacement histogram.
ii. The displacement histograms are fitted using the following equation.

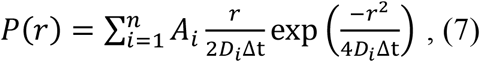
iii. “r” is the mean displacement 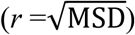 within a frame (Δt).
iv. “D_i_” is the diffusion coefficient of the i-th state.
v. “A_i_” is a coefficient that related to the fraction of the i-th state.
vi. “n” is the number of the states.
vii. “Min” and “Max” specify the range of “n” in the fitting function. Enter the natural number between 1 and 5. When you set the “Min” and “Max” values as “2” and “4”, the displacement histogram is fitted with the two-, three-, and four-state models. Then, the model with the lowest Akaike information criterion (AIC) value is selected. AIC is defined as follows:

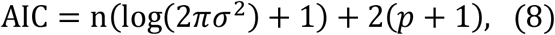

where n is the number of data points for the curve fitting, σ^2^ is the residual sum of squares (RSS), and p is the number of free parameters.
viii. To compare the fitting results among data from different cells, the fitting model should be fixed. It is recommended to select the model first by changing the number of states, then to do all the analysis with the selected number of states. Set the “Min” and “Max” values to the same value to fix the state number.
ix. The fitting result is sensitive to the initial values. If the fitting results are of less quality, change the initial values (A1-A5, D1-D5), and click the “Hist D” button to reanalyze them. Repeat it until there are no more errors messages.
x. In the HMM format analysis, each displacement histogram of the diffusion state is fitted with the function of the one state model of equation 7, where n=1. In this step, it is possible to visualize what kinds of step size distribution are assigned to each diffusion state by VB-HMM analysis.
34. The parameter settings for the intensity histogram analysis are as below. Refer to the Fig. S10-12 in the supplementary pdf file uploaded to the following URL as well (https://github.com/masataka-yanagawa/IgorPro8-smDynamicsAnalyzer).

i. “Bin” and “Dim” determine the X-axis range of the intensity histogram.
ii. The intensity histograms are fitted using the following equation.

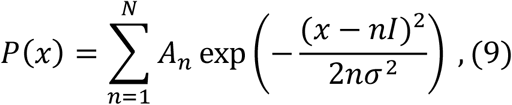
iii. “I” and “σ” in the equation are the mean and standard deviation (SD) of the intensity of a single fluorophore, respectively. “Mean” and “SD” in the parameter panel correspond to I and σ, respectively, in the curve fitting. When the checkbox “Fix Mean & SD” is activated, I and σ are fixed to the input values; otherwise, the input values are calculated as the initial values of the curve fitting.
iv. A_n_” is a coefficient that is connected to the fraction of the n-th state.
v. “n” is the oligomer size
vi. “N” is the maximum oligomer size to be considered in the model.
vii. “AIC: Min” and “Max” specify the range of “N” in the fitting function. Enter the natural number between 1 and 20. When you set the “Min” and “Max” values as “2” and “16”, the displacement histogram is fitted with the 2~16-state models. Then, the model with the lowest AIC value is selected (equation 8).
viii. To compare the fitting results among data from different cells, they should be taken from the same fitting model. It is recommended to select an appropriate model first by changing the number of states, then to do all the analysis with the selected number of states. Set the “Min” and “Max” values to the same value as the appropriate number of states in order to fix the state number.
ix. Check the fitting results to confirm that the Mean and SD values are within the appropriate range for a single-molecule intensity. In the cases of (i) the number of states in the fitting model is too large or (ii) the initial value is too small from the single-molecule intensity, the Mean and SD values are often estimated as less than half of the single-molecule intensity. It is usually difficult to estimate the number of states just from the AIC comparison in that situation. To determine the single-molecule intensity precisely, a control experiment is required, where a monomeric protein such as CD86 with the same FL label is measured in the same settings.
x. In the HMM format analysis, all the intensity histograms in the graph are globally fitted using the equation 9, where the I and σ are set as global variables
35. The parameter settings for the density analysis are as below. Refer to the Figs. S13-14 in the supplementary pdf file uploaded to the following URL as well for more details (https://github.com/masataka-yanagawa/IgorPro8-smDynamicsAnalyzer).

i. “Bin [um]” and “Dim” in Fig. 5a determine the X-axis range of the local density plots.
ii. “Start” and “End” in Fig. 5a define the range of the frames to be analyzed. The particle localization within the selected frames is plotted in the XY-plot as red dots (Fig. 5b, Start: 30 ~ End: 60). The “Start” frame should be more than “Min frame”, and the “End” frame should be less than “Frame Number - Min frame”, where the “Min frame” is the parameter in the “Measurement/SMT parameters” section on the parameter panel (Fig. 3a). In the SMT analysis, the lengths of the trajectories are biased at around both ends of the movies because the trajectories are broken off at both ends. To avoid such an edge effect in the SMT analysis, it is better to exclude the edge frames of the movies from the range of the density analysis. Because the density analysis of long frames is time-consuming, we recommend for “Start” to set “Min frame” and for “End” to set “Min frame + 5”.
iii. The mean local density plots are calculated from the distribution of the localization coordinates in each frame based on the following function,

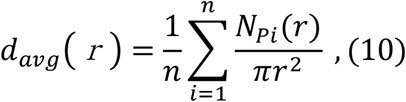

where “n” is the number of localization within the frame, and N_Pi_(r) is the number of points *N* within a distance *r* of point *i*. Namely, the red curves in Fig. 5b correspond to the mean local density around in the vicinity of r from each localization within a frame are plotted against r.
iv. When r is less than the distance from the most adjacent spot, the mean density function is defined as follows; d_avg_(r) = 1/ πr^2^. When r is more than that, the mean local density becomes closer to the mean density of spots in the cell. If the region of the cell were infinitely extended, the limit of the mean local density function would converge to the mean density. In reality, because the cell has the edge, the mean local density function reaches a plateau followed by the convergence to 0. The mean local density function decrease inversely proportional to πr^2^ when r becomes larger than the long axis radius of the cell.
v. Consequently, the mean density of the spots in the cell within each frame can be approximately estimated as the plateau value of the mean local density function. The plateau value is estimated from the first-order difference of the mean local density curves (the blue curves in Fig. 5c). The mean local density at the peak of the blue curves, is adopted as the mean density of each frame. r_dmax_ is the distance which gives the mean density.
vi. “Smoothing: frames” in Fig. 5a is a parameter to find peak (r_dmax_) in the middle panel. The blue curves are plotted after smoothing by a moving average with the window width of the input value to find the peak properly. Increase it if the peak detection was improper due to the noise.
vii. The blue circles with r_dmax_ are plotted around each localization in Fig. 5b. The region of the circles does not cover the cell region if the peak detection is improper.
viii. The estimated density and area surrounding the spots (cell area) in each frame are plotted in Fig. 5d. The cell area is estimated as “number of spots/density”. The average values of all the analyzed frames are used as the representative values of the density and area of the cell in the “Comparison analysis”. (*see* Section 3.8.19)
ix. The mean density of spots is often calculated by dividing the number of spots by the area of the cell region that manually enclosed around the spots (Fig. 5e). However, the “by hand method” is time consuming and difficult to reproduce. The density analysis macro automatically provides the mean density of spots comparable to that estimated from the conventional “by hand method” (Fig. 5e). Even the analysis of a single-frame can provide estimation results that are comparable to those of manual analysis (Fig. 5e).
36. The interpretation of “On-rate” and “Off-rate” The terms “On-rate” and “Off-rate” are intended to analyze the on-and off-rates of a cytoplasmic protein to the PM in a steady state, or of a protein in solution to the coverslip *in vitro.* For example, we previously analyzed the changes of the on- and off-rates of wild-type and mutants of Sos [22], Raf [23], and RalGDS [24] proteins to the PM upon EGFR activation, which suggested the role of each domain of the proteins in the interaction with signaling molecules such as Ras (Fig. 6a) In the case of the analysis of a membrane protein including GPCR, the terms “On-rate” and “Off-rate” are considered to be inverse to the reality in a steady state. For example, one of the trajectories ends when two molecules associate to form a dimer, which is counted as an event regarding “Off-rate”. In contrast, when a new trajectory starts when the dimer dissociates into two monomers, which is counted as an event regarding “On-rate” (Fig. 6b).
37. The parameter settings for the off-rate analysis are as below. Refer to the Figs. S15-16 in the supplementary pdf file uploaded to the following URL as well (https://github.com/masataka-yanagawa/IgorPro8-smDynamicsAnalyzer).

i. The decay curves are fitted using the following equation,

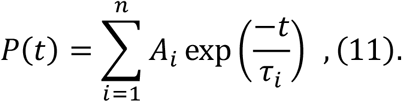
ii. “A_i_” is a coefficient that related to the fraction of the i-th state. The input values “A1”~”A5” in the parameter panel are calculated as the initial values of the curve fitting.
iii. “τ_i_” is the exponential decay time constant of the i-th state. The input values “Tau1”~”Tau5” in the parameter panel are calculated as the initial values of the curve fitting.
iv. “n” is the maximum number of exponential to be considered in the model.
v. “Min” and “Max” specify the range of “n” in the fitting function. Enter the natural number between 1 and 5. When you set the “Min” and “Max” values as “1” and “3”, the displacement histogram is fitted with the single, double, and triple exponential functions. Then, the model with the lowest AIC value is selected.
vi. If the check box “Frame Edge Correction” is checked, the decay curve plot is created except for the trajectories that start in the first frame or ends in the last frame to avoid the edge effect of SMT analysis.
38. The parameter settings for the on-rate analysis are as below. Refer to the Figs. S17-18 in the supplementary pdf file uploaded to the following URL as well (https://github.com/masataka-yanagawa/IgorPro8-smDynamicsAnalyzer).

i. The cumulative event number plots are fitted using the following equation.

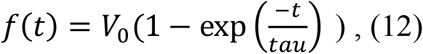
ii. “V_0_” is the initial slope of the cumulative event number plot. The input value in the parameter panel is calculated as the initial value of the curve fitting.
iii. “Tau” is the exponential decay time constant, which is required when the cumulative event number plot is not linear due to the photo bleaching. The input value in the parameter panel is calculated as the initial value of the curve fitting.
iv. The on-rate is calculated as V_0_/Area_cell_. If the check box “Fix Area [μm2]” is unchecked, the area of cell estimated from the density analysis is assigned to Areacell (*see* **Note 35**); otherwise Area_cell_ was fixed as the input value. The latter is useful for *in vitro* single-molecule analysis or for the analysis of a cell region cropped to a certain size.
39. Refer to the Figs. S19-20 in the supplementary pdf file uploaded to the following URL as well (https://github.com/masataka-yanagawa/IgorPro8-smDynamicsAnalyzer). Matrix folder contains a side-by-side matrix wave of each analysis results in the “Folder Name” folders. Results folder contains the stats waves with suffix “mean”, “sd”, “sem”, and “n”. At least three data are required to calculation of the stats waves.
40. In the case of “Total Basic Analysis”, the name of each folder (e.g. “A1” ~ “A5”) is entered as “Sample Name” in the macros automatically. Do not set too long folder names and/or folder names with space.
41. In the example in Fig. 7, the five folders “A1” ~ “A5” are sequentially analyzed by the “Respective Analysis” as shown in the sections 3.8.2 ~ 3.8.11. Then, “Comparison Analysis” that output graphs and tables that compare the analysis results in the Matrix or Results folders (*see* 3.8.13 ~ 3.8.20).
42. Refer to the Figs. S21-22 in the supplementary pdf file uploaded to the following URL as well (https://github.com/masataka-yanagawa/IgorPro8-smDynamicsAnalyzer). Preexistence of “Comparison” folder in the Igor data browser sometimes causes an error. If you got an error message, delete the “Comparison” folder under the root folder. [Open data browser (Ctrl + B), right click the “Comparison” folder, and select “delete object”.] (*see* **Notes 26~28**)
43. The multiple comparison macros run the program built into Igor as shown below; ANOVA1: Statistics > One-way ANOVA test (Welch test, Alpha: 0.05) Dunnett-test: Statistics > Multi-comparison test > Dunnett (Alpha: 0.05, tail: mean1 = mean2). The left most group in the violin/box plot is selected as the control wave. Tukey-test: Statistics > Multi-comparison test > Tukey (Alpha: 0.05) Dunn-test: Statistics > Multi-comparison test > Non-parametric multiple contrasts > Dunn-Holland-Wolfe test (Alpha: 0.05, tail: mean1 = mean2)
44. The violin/box plot is a graph that the violin and box plots are merged. One can remove any of the plots by right clicking the graph and selecting “Remove from Graph”. The format of the plots can be changed through following the path: “Graph > Modify Box Plot” or “Graph > Modify Violin Plot”. The default settings of the plots are as follows: Violin plot: Bandwidth Method: Silverman, Kernel: Gaussian, Show Data points as open circle, Show Mean as closed diamond. Box plot: Whisker Method: One standard deviation, Quartile Method: Tukey, Show Median as line, Show data as open circle, Show outliers as open red circle, Show mean as red diamond.
45. In the 2D plot of Fig. 8v, the mean oligomer size of mGluR3 shows a positive correlation to the particle density as previously described in [4]. Each point corresponds to the data from the cell. The dashed lines show linear fitting of the points in the same “Sample Name” folder.
46. The example data in Fig. 10 are the TMR channel data in the dual color SMI of mGluR3-TMR and the CLC-EGFP (Fig. 1). The VB-HMM data in two ligand conditions (Inactive: 100 nM LY341495, and Active: 100 μM LY379268) are compared.
47. The agonist induces decrease of the diffusion coefficient of mGluR3 in the immobile, slow, and medium states to a statistically significant extent as observed (Fig. 10e-g), which is consistent with the MSD-Δt plots (Fig. 10a-c).
48. The agonist induces decrease of the medium state fraction of mGluR3, and increase of the immobile state fraction to a statistically significant extent as observed (Fig. 10i and k).
49. The macro also generates the line-graphs of the partial oligomer size distribution of each diffusion state that is normalized with the total sum of all the 4 states. Namely, the plotted values are the product of “percentile of each oligomer size in each state” and “ fraction of each state”.
50. The example data in Figs. 11 and 12 show the dual color SMI of mGluR3-TMR and the CLC-EGFP (Fig. 1). The VB-HMM data were compared in two ligand conditions (Inactive: 100 nM LY341495, and Active: 100 μM LY379268). In advance, we performed “Total Basic Analysis” of TMR channel (“mGlu3_LY34” & “mGlu3_LY37”) as shown in 3.8.21, and of GFP channel (“CLC_LY34” & “CLC_LY37”).
51. The parameter settings for the colocalization analysis are as below. The parameters, “Max distance [nm]” and “Max D ratio”, are the thresholds to define colocalization (Fig. 11a). The colocalization of two particles is detected if the two particles are within the “Max distance [nm]” in the same frame, and if the diffusion coefficient ratio of the two particles is less than “Max D ratio”. When judging the ratio of the diffusion coefficient of two particles (D_1_/D_2_) are less than “Max D ratio” or not, the D_1_/D_2_ is calculated from the step size (r_1_ and r_2_) between two frames, where D_1_/D_2_ = (r_1_/ r_2_)^2^ and D_1_ > D_2_. If you do not want to set the “Max D ratio” condition, set the value large enough to ignore it. Here we set the “Max distance [nm]” as 100 nm based on the localization accuracy (20~30 nm in each channel) and “Max D ratio” as 100 to reduce the detection of random colocalization. If the checkbox “Same HMM D-state” is checked (Fig. 11a), the colocalization is defined as the two particles meet the criteria: (i) the distance between two particles is within the “Max distance [nm]” in the same frame, and (ii) the two particles are in the same diffusion state. In this case, the macro ignores the “Max D ratio” condition. To avoid the fragmentation of the colocalization trajectory due to the photoblinking of the fluorophores and/or an analytical error, a non-colocalization frame sandwiched between two colocalization frames is defined as a colocalization frame in the macro.
52. The “Find Col.” macro automatically creates the “Col” and “EC” folders under the root folder (Fig. 11a). The trajectories where any length of colocalization is detected are displayed in red (SampleName1) and green (SampleName2) in the XY plots. These trajectories are stored in the “SampleName1” and “SampleName2” subfolders in the “Col” folder. Trajectories in the colocalization frames, which were extracted into the “SampleName1” and “SampleName2” subfolders in the “EC” folder, are displaced in yellow in the XY plots (Fig. 11b). All the colocalization analysis results is stored in the “EC” folder.
53. The “Hist D” macro in the “Colocalization Analysis” automatically performs the analysis as described in the section 3.8.25 (see **Note 33**) using the waves in the “SampleName1” and “SampleName2” subfolders inside the “EC” folder. The small number of the colocalization frames often causes a curve fitting error. Ignore the error and skip this process in that case.
54. The “On-rate” macro in the “Colocalization Analysis” automatically performs the analysis as described in the section 3.8.10 (see **Note 38**) using the waves in the “SampleName1” and “SampleName2” subfolders inside the “EC” folder.
55. The “Off-rate” macro in the “Colocalization Analysis” automatically performs the analysis as described in the section 3.8.9 (see **Note 37**) using the waves in the “SampleName1” and “SampleName2” folders inside the “EC” folder.
56. These macros are not included in the “Colocalization Analysis”.
57. In the “Intensity” analysis, the same analysis in the section 3.8.23 is performed in the “SampleName1” and “SampleName2” subfolders inside the “EC” folder.
58. In the example in Fig. 12, the “CLC_LY34” and “CLC_LY37” folders are moved to “Package” folder” by drag and drop to compare the “mGlu3_LY34” and “mGlu3_LY37” folders (Fig. 12a).
59. In the way of the “Comparison Parameters” analysis after “Colocalization Analysis”, error signs may pop up if “Comparison” folder exists inside the “EC” folder. If you get an error message, delete the “Comparison” folder in the “EC” folder. [Open data browser (Ctrl +B), right click the “Comparison” folder, and select “delete object”.]
60. “P_col_” macro outputs the violin/box plots of the colocalization rate (percent total steps), and of the binding affinity (K_B_), which were calculated in the “Find Col.” analysis (*see* 3.8.31). In the example, the P_col_ and K_B_ were significantly increased upon activation of mGluR3 (Fig. 12b, c). The P_col_ and K_B_ were defined as follows;

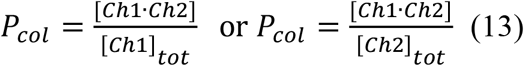

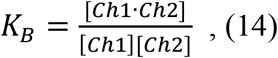

where [ch1 · ch2] = (the number of the colocalization frames)/Area_cell_, [ch1]_tot_ = (the total step number of ch1)/Area_cell_, [ch2]_tot_ = (the total step number of ch2)/Area_cell_, [ch1] = [ch1]_tot_ - [ch1 · ch2], and [ch2] = [ch2]_tot_ - [ch1 · ch2]. The Area_cell_ was estimated from the density analysis (*see* **Note 35**. and Fig. 5).
61. “On-rate” macro outputs the violin/box plots of the on-event rate parameter. In the example, the association rate between mGluR3 and CLC is increased to a statistically significant extent upon activation (Fig. 12d).
62. “Off-rate” macro outputs the mean decay curves of the colocalization duration (mean ± sem of the cells), and the violin/box plots of the off rate parameters. In the example, the decay time constants of the mGluR3/CLC complex is increased to a statistically significant extent upon activation (Fig. 12e).
63. “D ratio” macro outputs the violin/box plots of the fractions of the each diffusion state in the colocalize steps, where the sum of the ratio of 4 states should be 100%. In the example, significantly observed are the increase in the immobile state fraction and decreases in the medium and the fast state fractions in the colocalization frames of mGluR3 upon its activation (Fig. 12f-i). Then, the macro also outputs the violin/box plots of the product of “D ratio” and “ Pcol”, where the sum of the colocalization ratio of 4 states is “Pcol”. In the example, the increases of immobile, slow, and medium state fractions are observed to a statistically significant extent due to the increase of mGluR3/CLC complex upon activation (Fig. 12j-m).

## Acknowledgments

We thank J. Nathans (Johns Hopkins University) for providing us with the HEK 293S cell line, M. Murata for the cDNA of EGFP-tagged CLC, and R. D. Vale (University of California San Francisco) for plugins of ImageJ. This work was supported by the Ministry of Education, Culture, Sports, Science and Technology (MEXT), Japan [Grants-in-Aid for Scientific Research 19H05647 for Y. S and 16K18533, 20K05760 for M.Y], and RIKEN Pioneering Project, Integrated Lipidology, and Glycolipidologue Initiative for Y. S.

