## Supplementary figures for "Total workflows of the single-molecule imaging analysis in living cells: a tutorial guidance to the measurement of the drug effects on a GPCR"

Fig. S1 How to use the "Load" macro

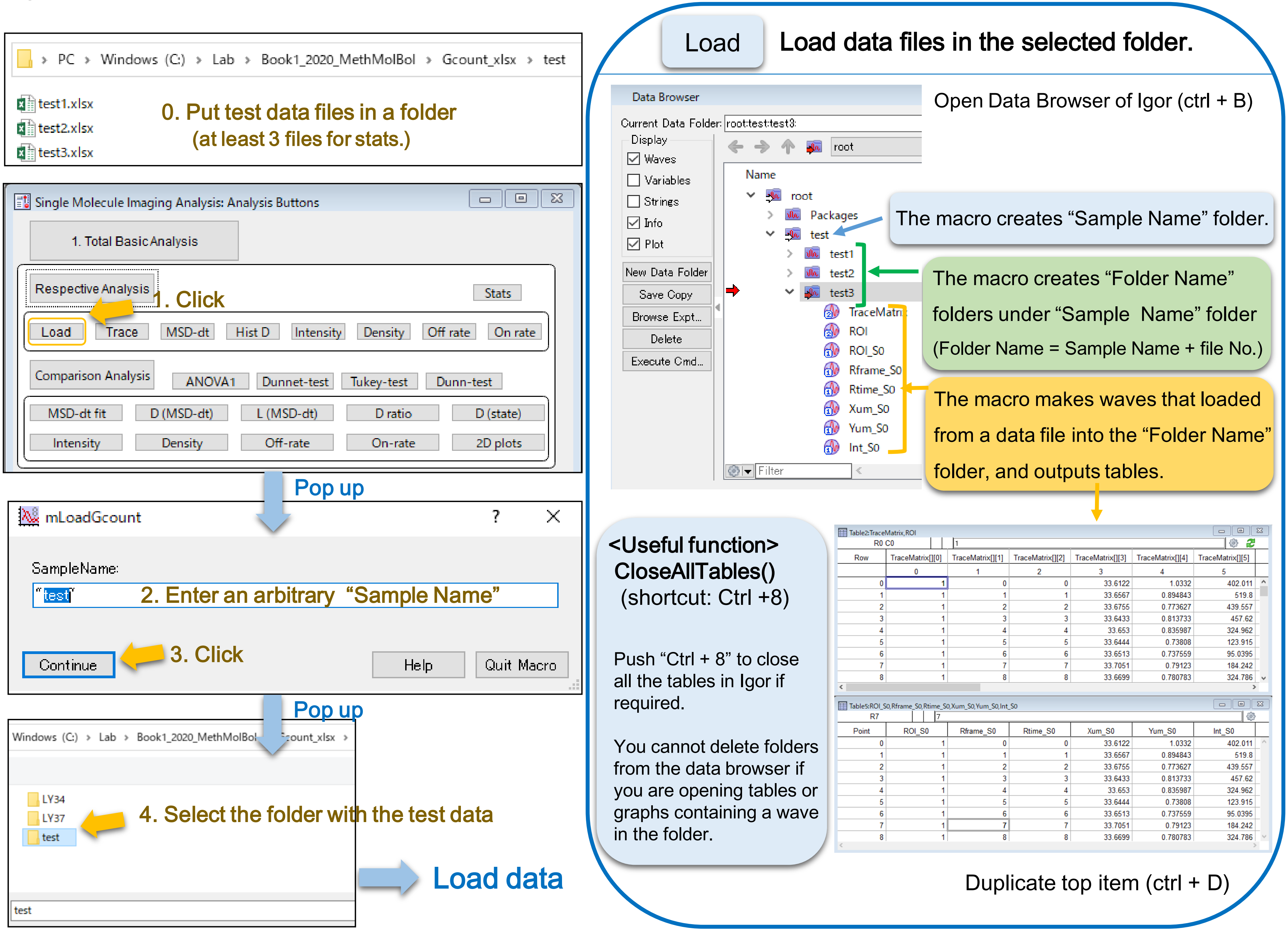

### Fig. S2 How to use the “Trace” macro

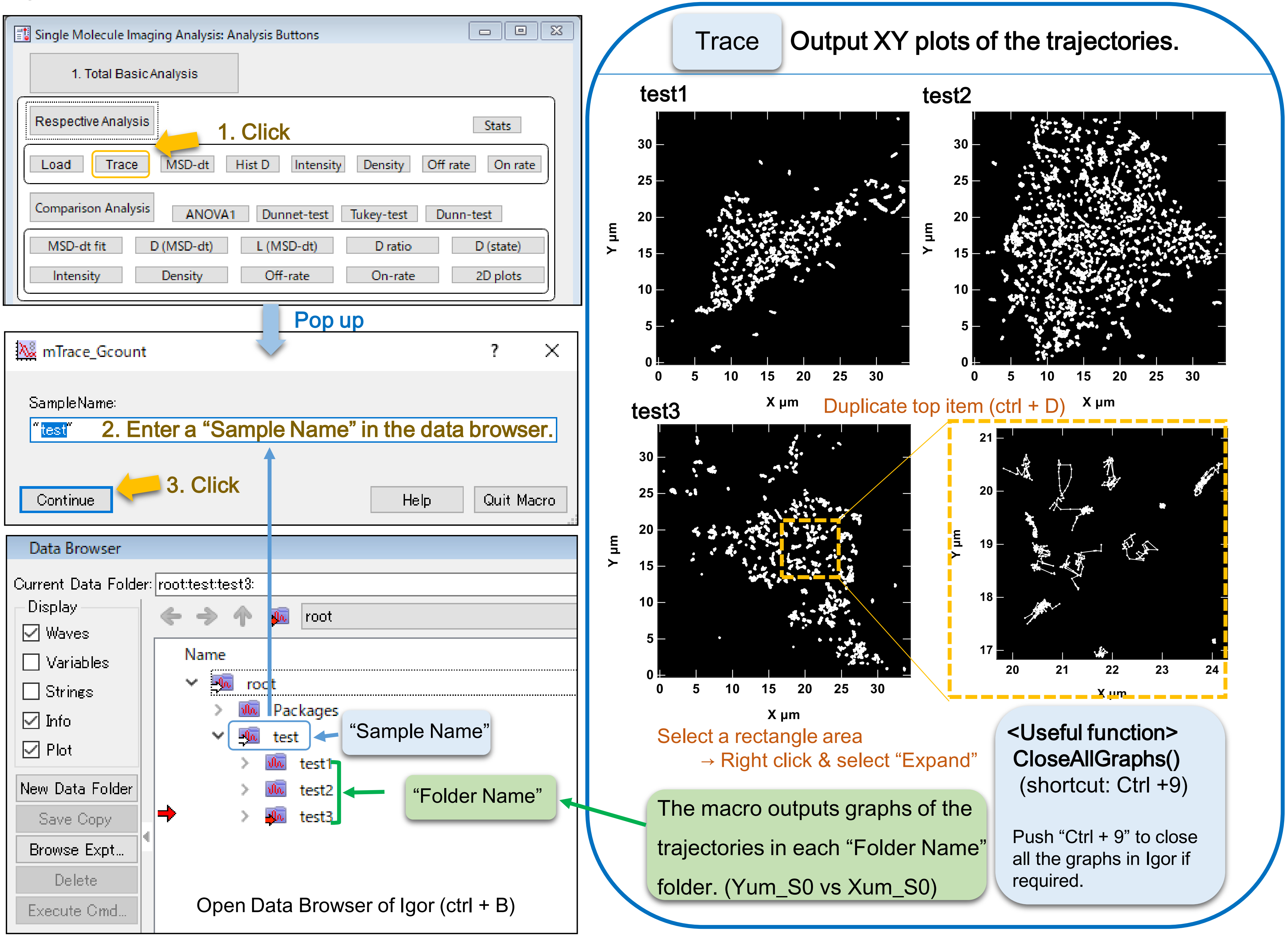

Fig. S3 How to use the “Trace” macro (HMM format)

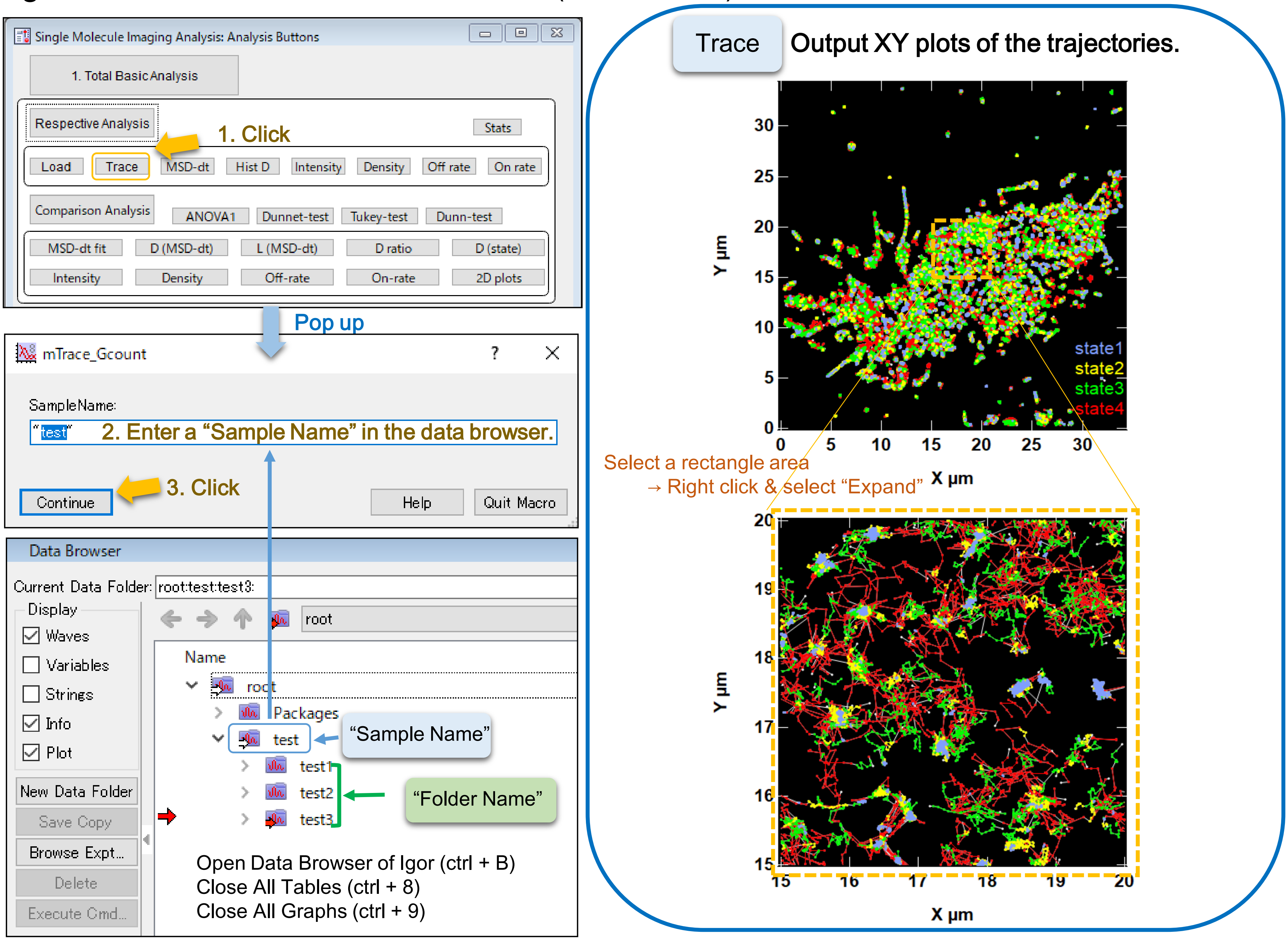

### Fig. S4 How to use the “MSD-dt” macro

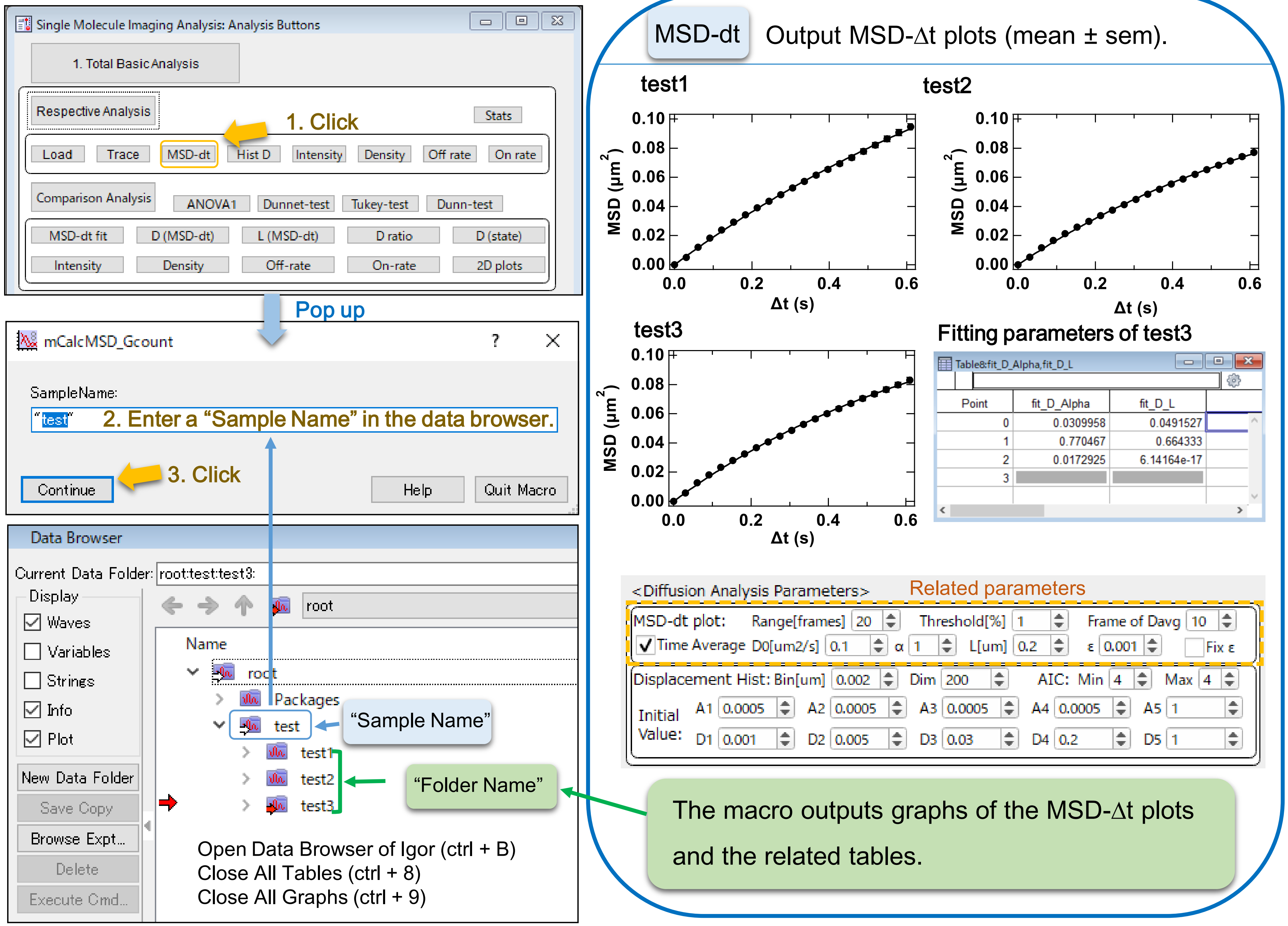

Fig. S5 Parameter settings of the “MSD-dt” macro

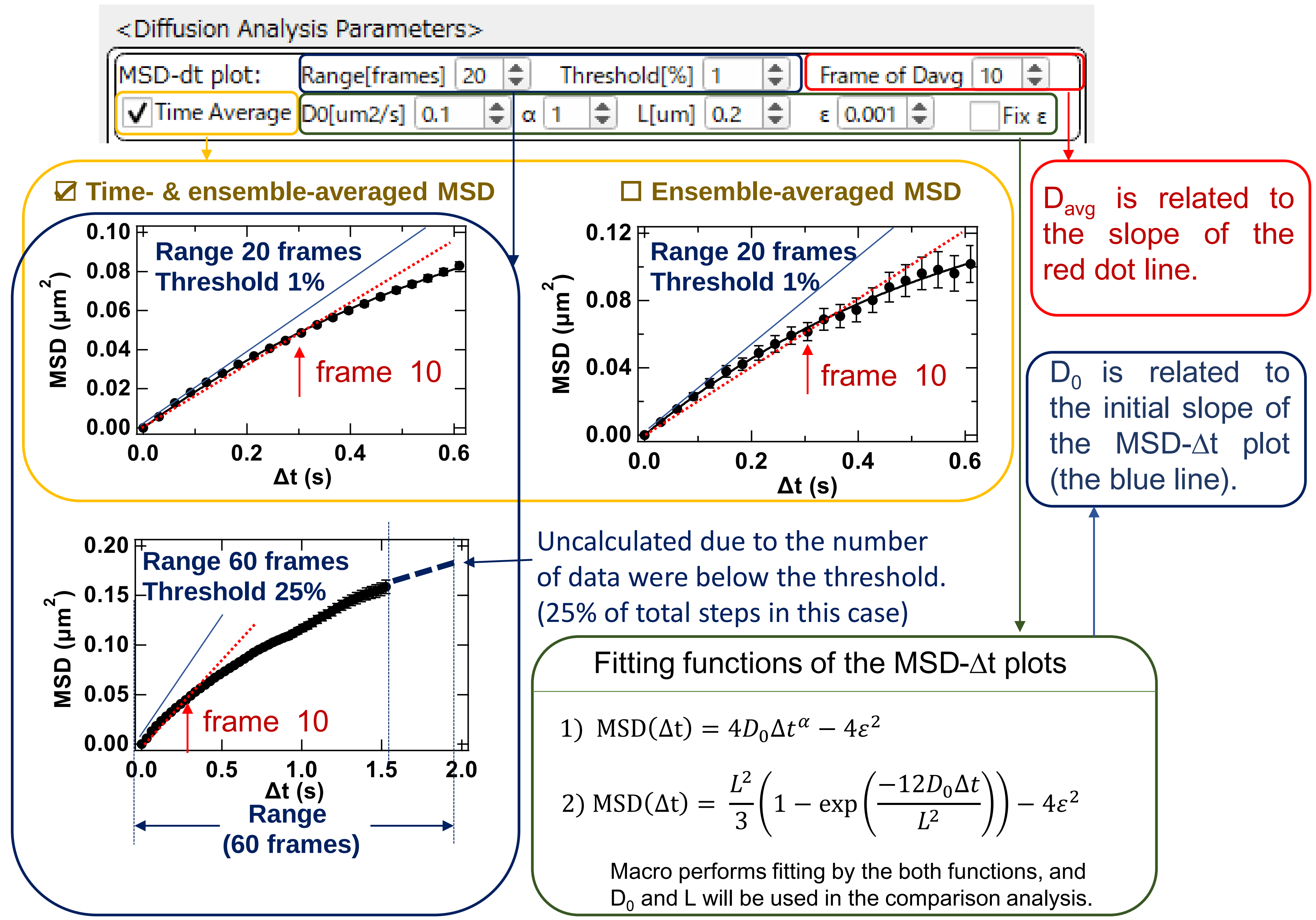

Fig. S6 How to use the “MSD-dt” macro (HMM format)

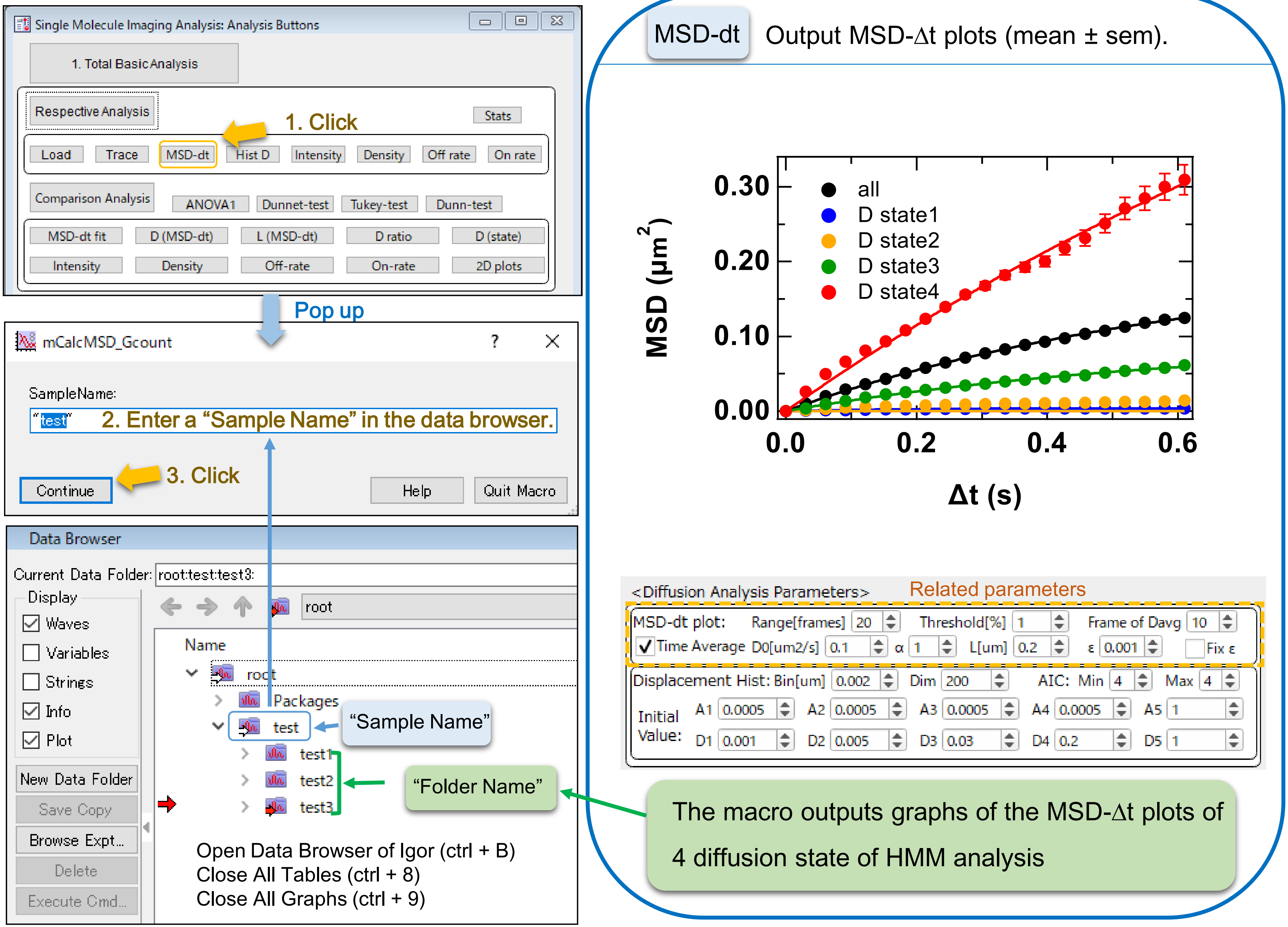

### Fig. S7 How to use the “Hist D” macro

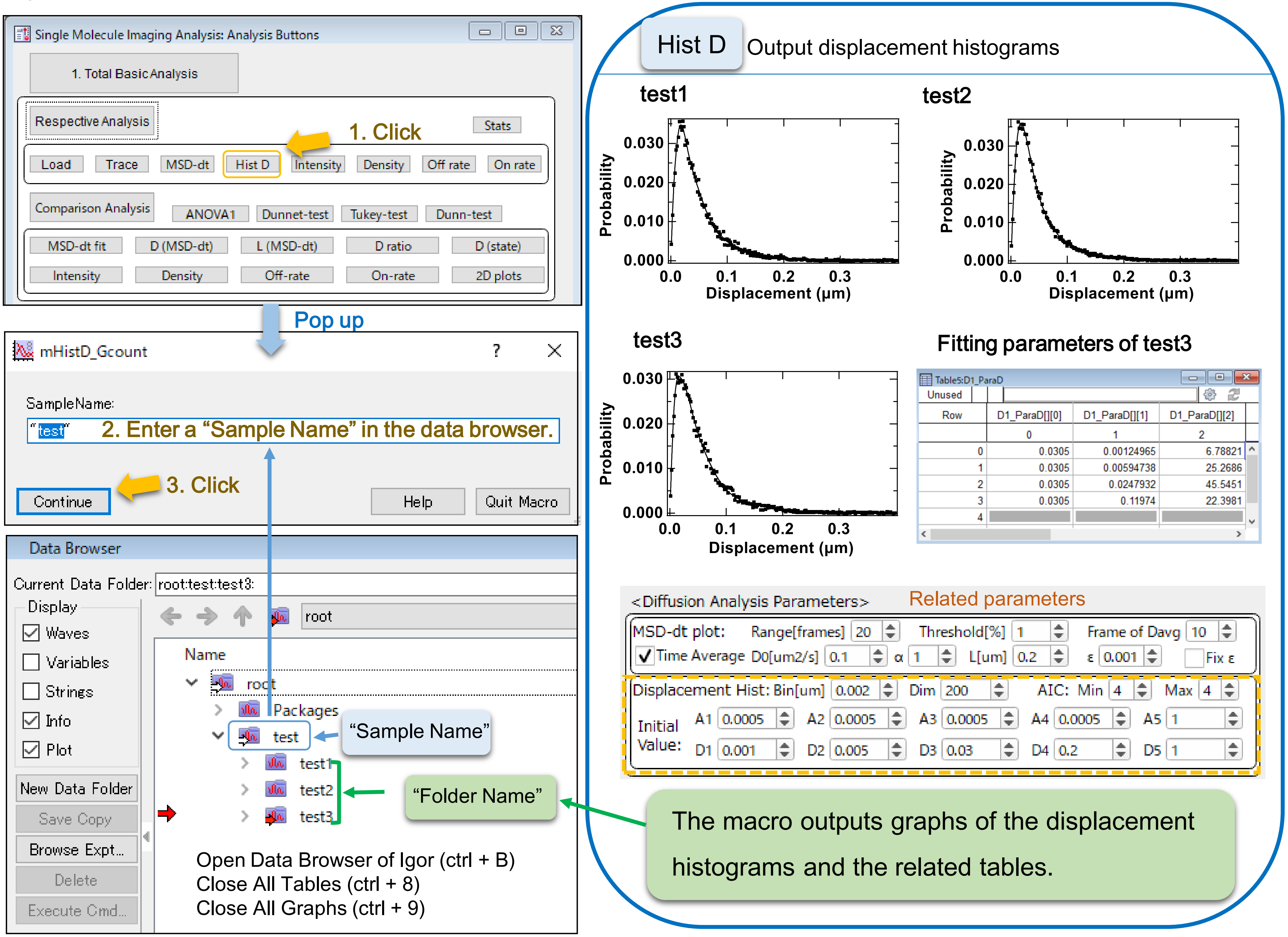

Fig. S8 Parameter settings of the "Hist D" macro

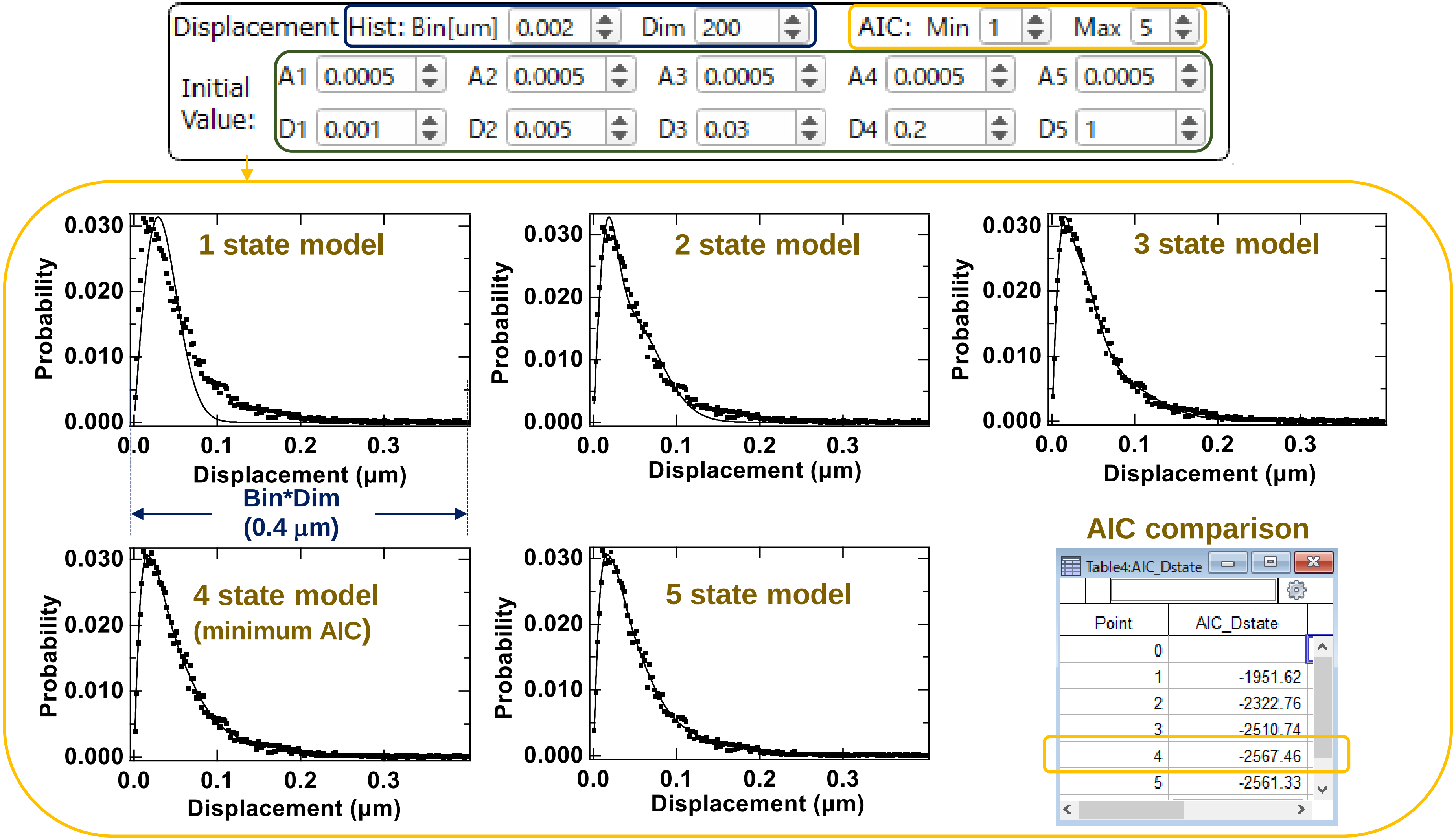


#### Fitting function of the displacement histogram

$$P(r) = \sum_{i=1}^{n} A_i \frac{r}{2D_i\Delta t} \exp\left(\frac{-r^2}{4D_i\Delta t}\right)$$

You can compare Min (1) to Max (5) state fitting models by AIC, but the fitting result is sensitive to the initial values. Check the fitting result of each model if the error message outputs as below .

```
error: singular matrix or other numeric error
```

### Fig. S9 How to use the “Hist D” macro (HMM format)

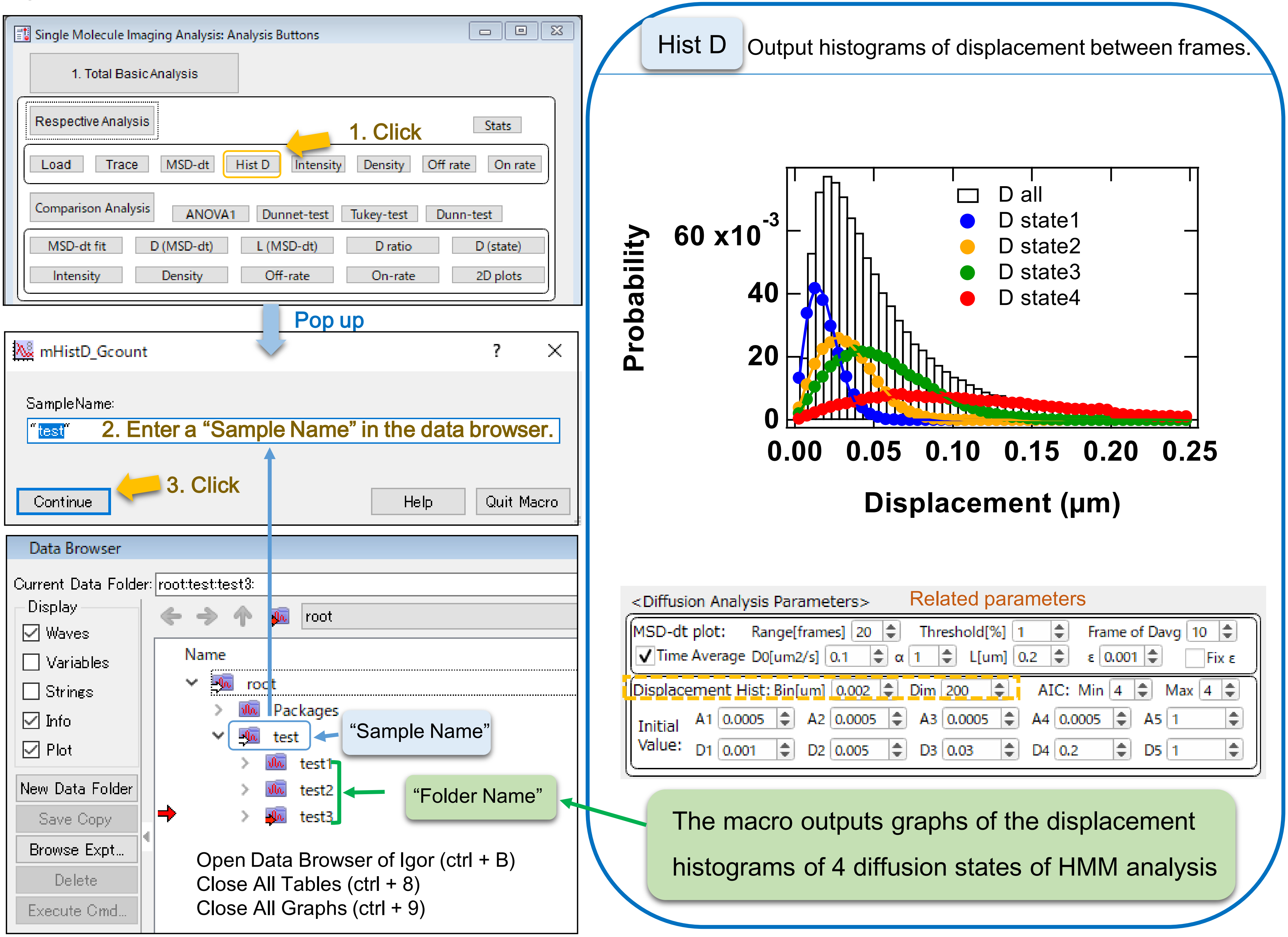

### Fig. S10 How to use the “Intensity” macro

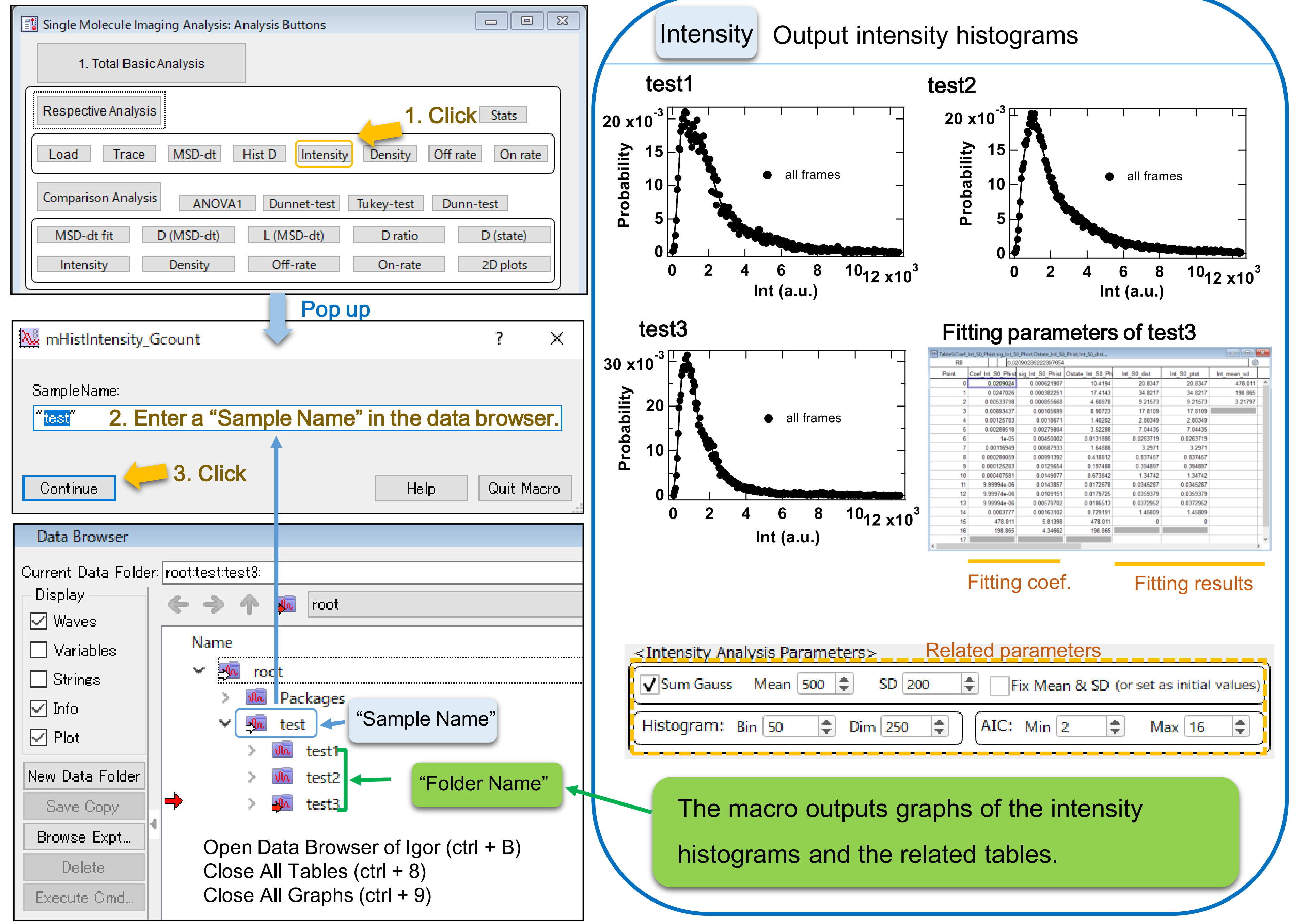

Fig. S11 Parameter settings of the “Intensity” macro

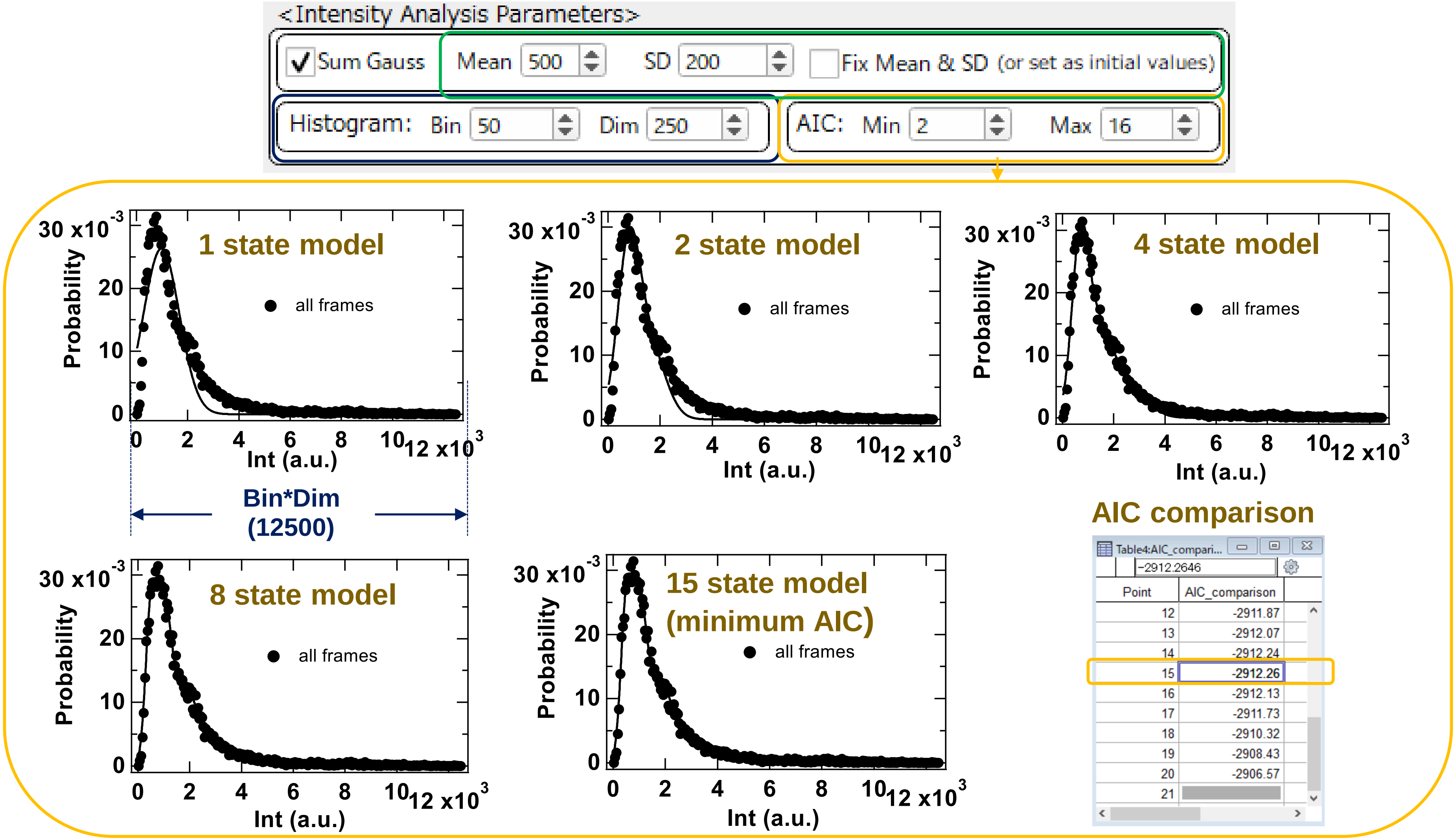


Fitting function of the intensity histogram

$$P(x) = \sum_{n=1}^{N} A_n \exp\left(-\frac{(x - nI)^2}{2n\sigma^2}\right)$$

You can compare Min (1) to Max (20) state fitting models by AIC. Check the fitting results to confirm the Mean and SD values are within the appropriate range as a single-molecule intensity.

The Mean and SD values are fixed to the input values when the checkbox is checked.

### Fig. S12 How to use the “Intensity” macro (HMM format)

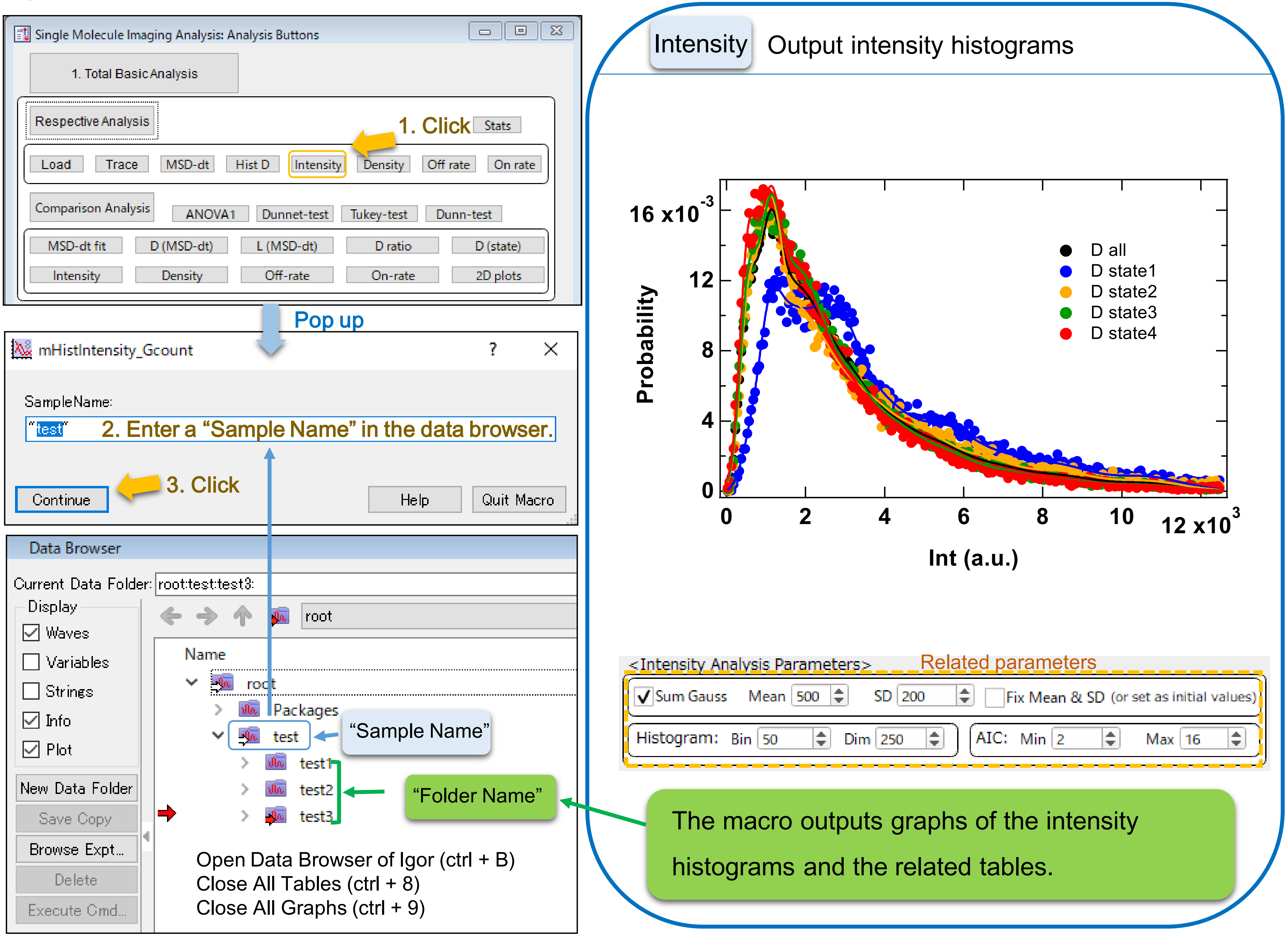

### Fig. S13 How to use the "Density" macro

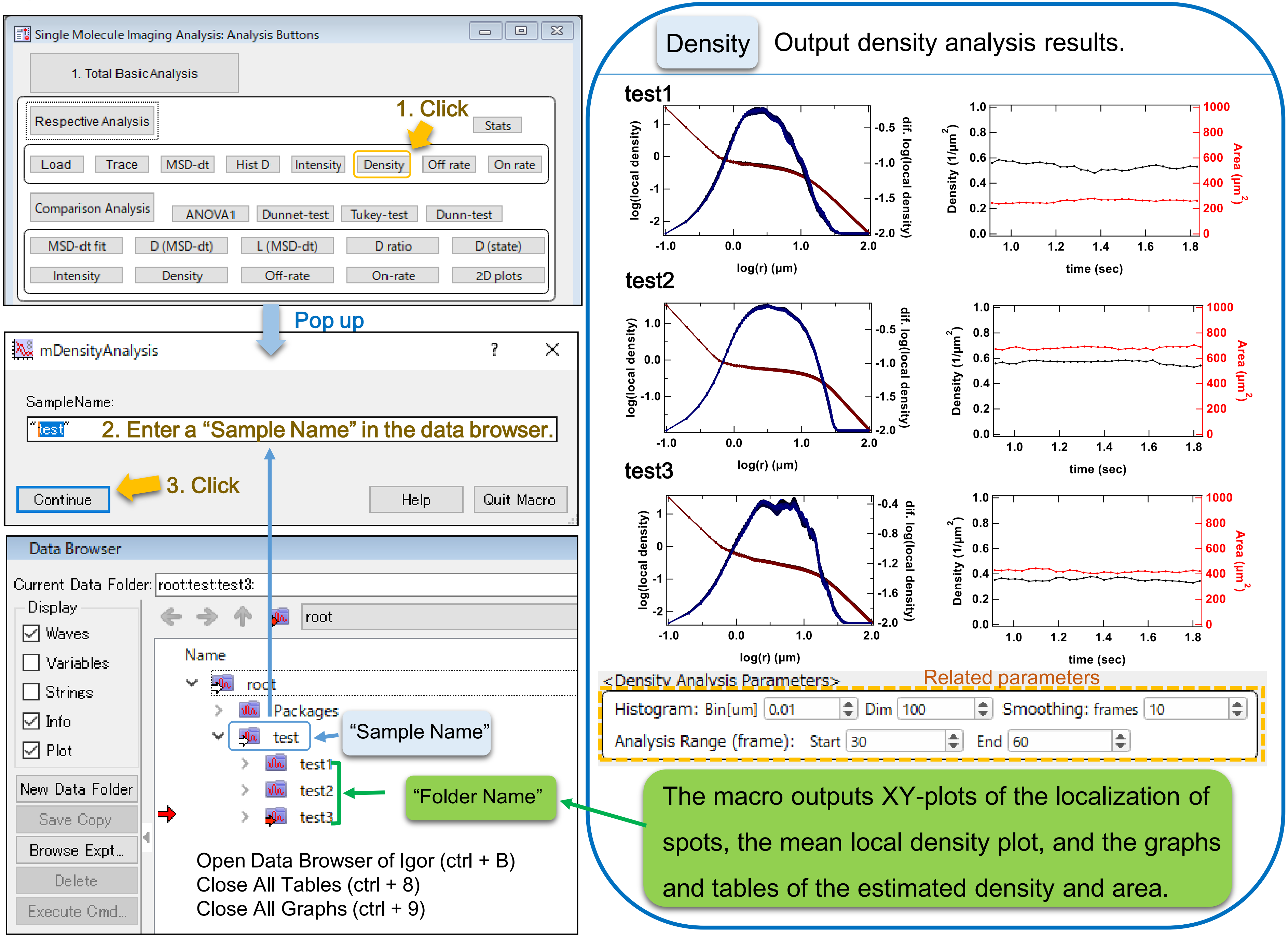

### Fig. S14 Parameter settings of the "Density" macro

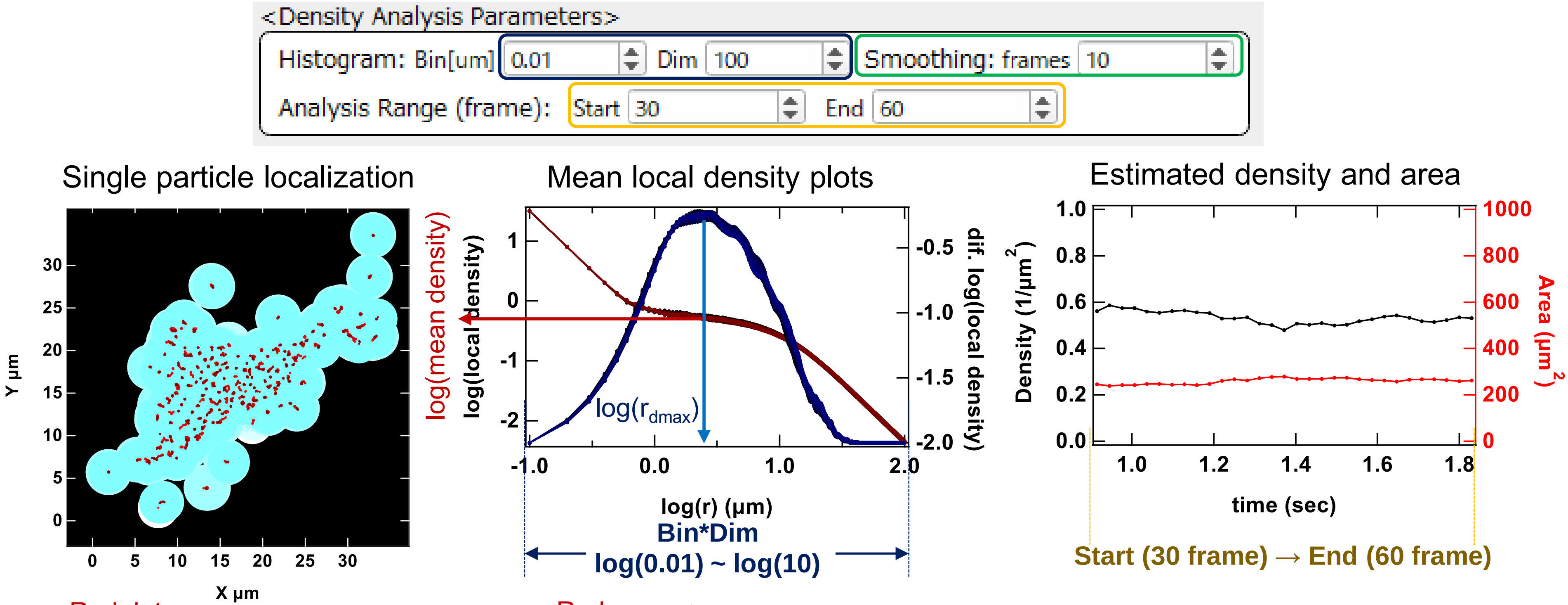


<Red dots>
The particle localization within the selected frames (Start: 30 ~ End: 60) are plotted.

<Blue circles around the red dots>
Circles with radius ($r_{dmax}$) that gives the plateau of the mean local density curve (middle panel) are plotted around each localization.

<Red curves>
The mean local density around in the vicinity of r from each localization within a frame are plotted against r.

<Blue curves>
First-order derivative of the mean local density curves. The local density in the vicinity of $r_{dmax}$, which gives the peak, is adopted as the mean density of particles at each frame.

<Black plots>
The mean density at each frame are plotted against time.

<Red plots>
The area of cell region that was estimated from the mean density and particle number are plotted against time.

#### The mean local density function

$$d_{avg}(\mathrm{r}) = \frac{1}{n}\sum_{i=1}^{n}\frac{N_{Pi}(r)}{\pi r^2}$$

($N_{Pi}(r)$: number of points $N$ within a distance $r$ of point $i$)

A parameter to find peak ($r_{dmax}$) in the middle panel.

Increase it if the peak detection was improper due to the noise.

Fig. S15 How to use the "Off rate" macro

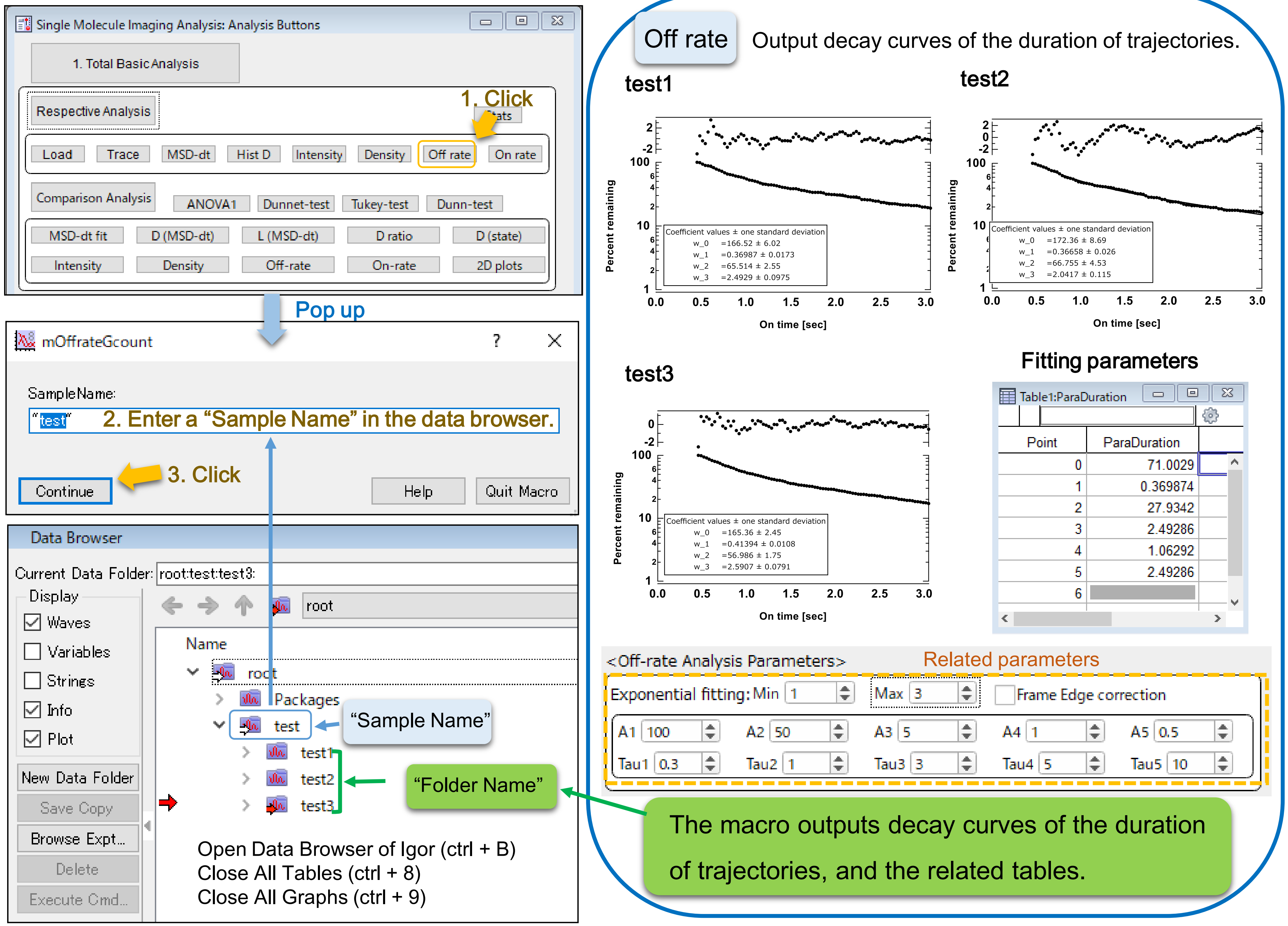

Fig. S16 Parameter settings of the “Off rate” macro

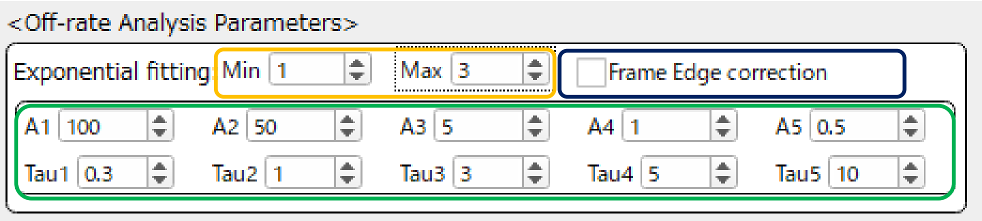


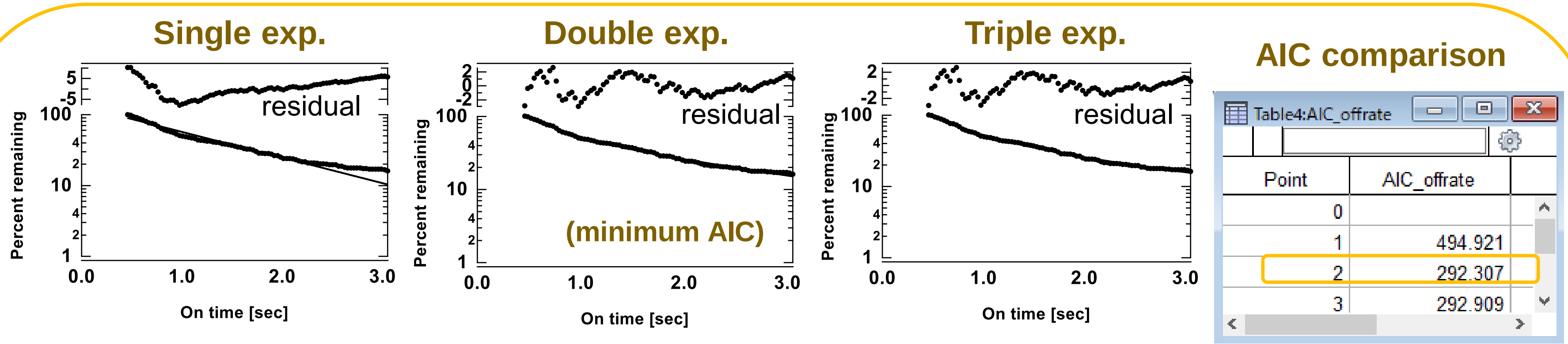


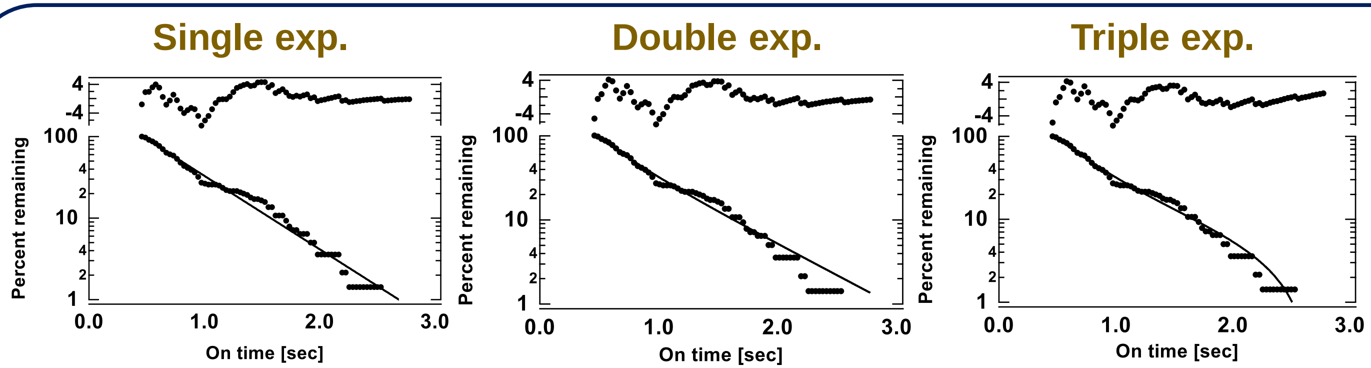


If the check box is checked, the plot is created except for the trajectories that start in the first frame or ends in the last frame.

#### Fitting function of the decay curve

$$P(t) = \sum_{i=1}^{n} A_i \exp\left(\frac{-t}{\tau_i}\right)$$

You can compare Min (1) to Max (5) state fitting models by AIC, but the fitting result is sensitive to the initial values. Check the fitting result of each model if the error message outputs as below .

```
error: singular matrix or other numeric error
```

Fig. S17 How to use the "On rate" macro

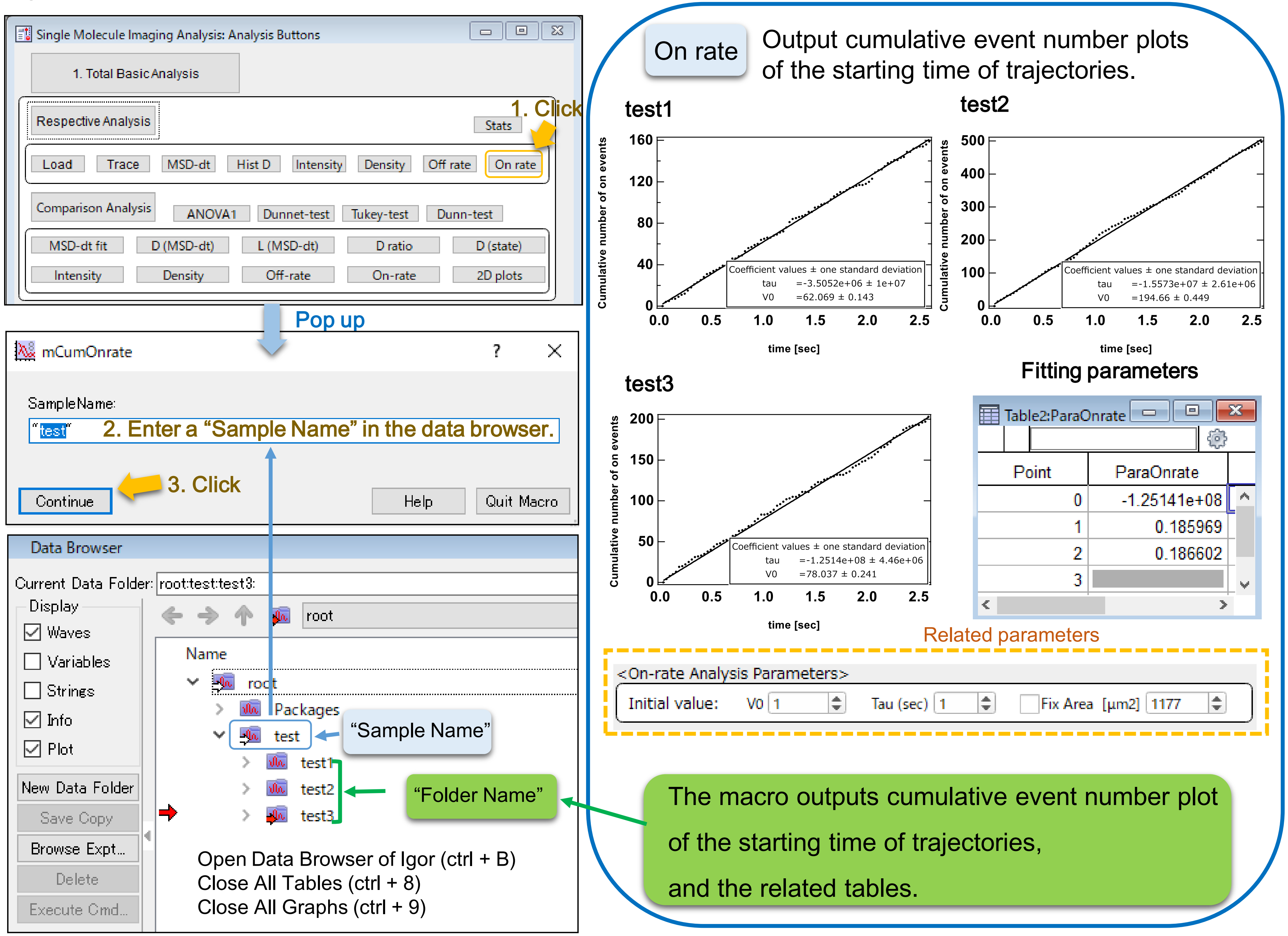

Fig. S18 Parameter settings of the “On rate” macro

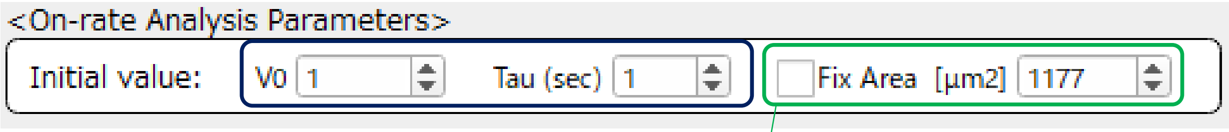


On-rate analysis should be done after the density analysis

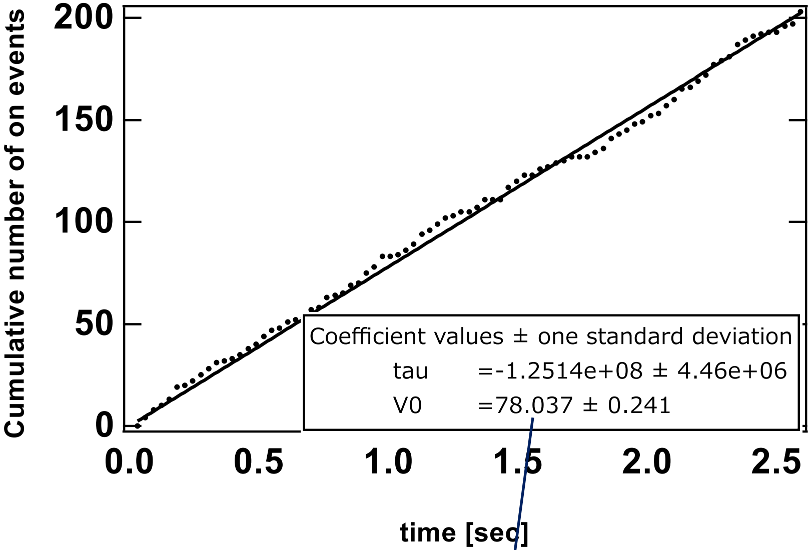


$$\text{on-rate} = V_0 \;/\; \text{Area}_{\text{cell}}$$

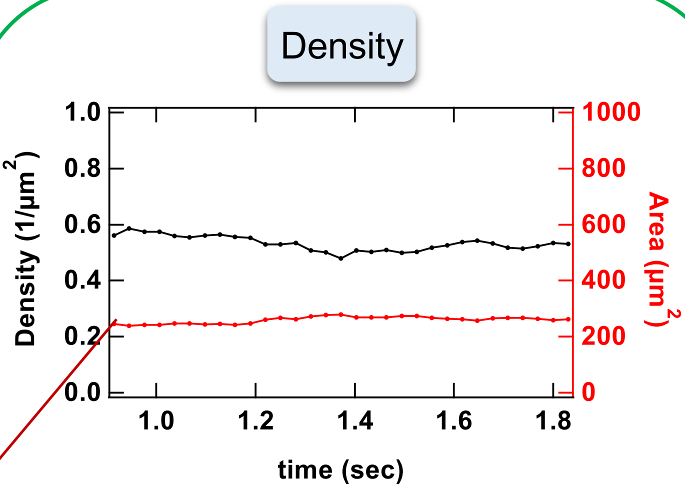


If the check box is unchecked, the $\text{Area}_{\text{cell}}$ estimated from the density analysis is used for the calculation of the on-rate.

If checked, the input value is used for the calculation of the on-rate.

Fitting function of the plot

$$f(t) = V_0\left(1 - \exp\left(\frac{-t}{tau}\right)\right)$$

V0 is the initial slope of the cumulative event number plot.

Tau is required when the cumulative event number plot is not linear due to the photo bleaching.

### Fig. S19 How to use the “Stats” macro

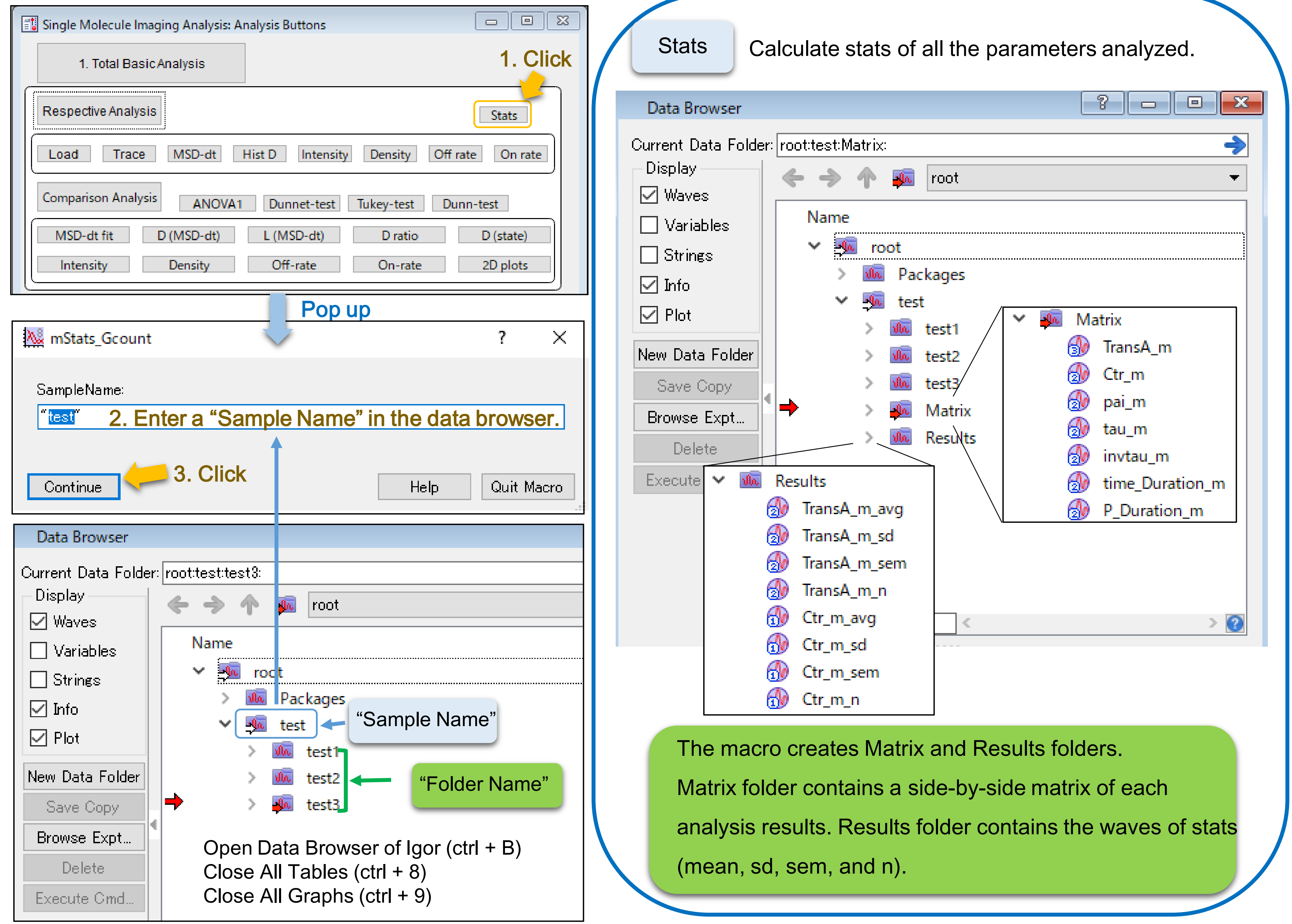

### Fig. S20 Example waves in the Matrix and Results folders

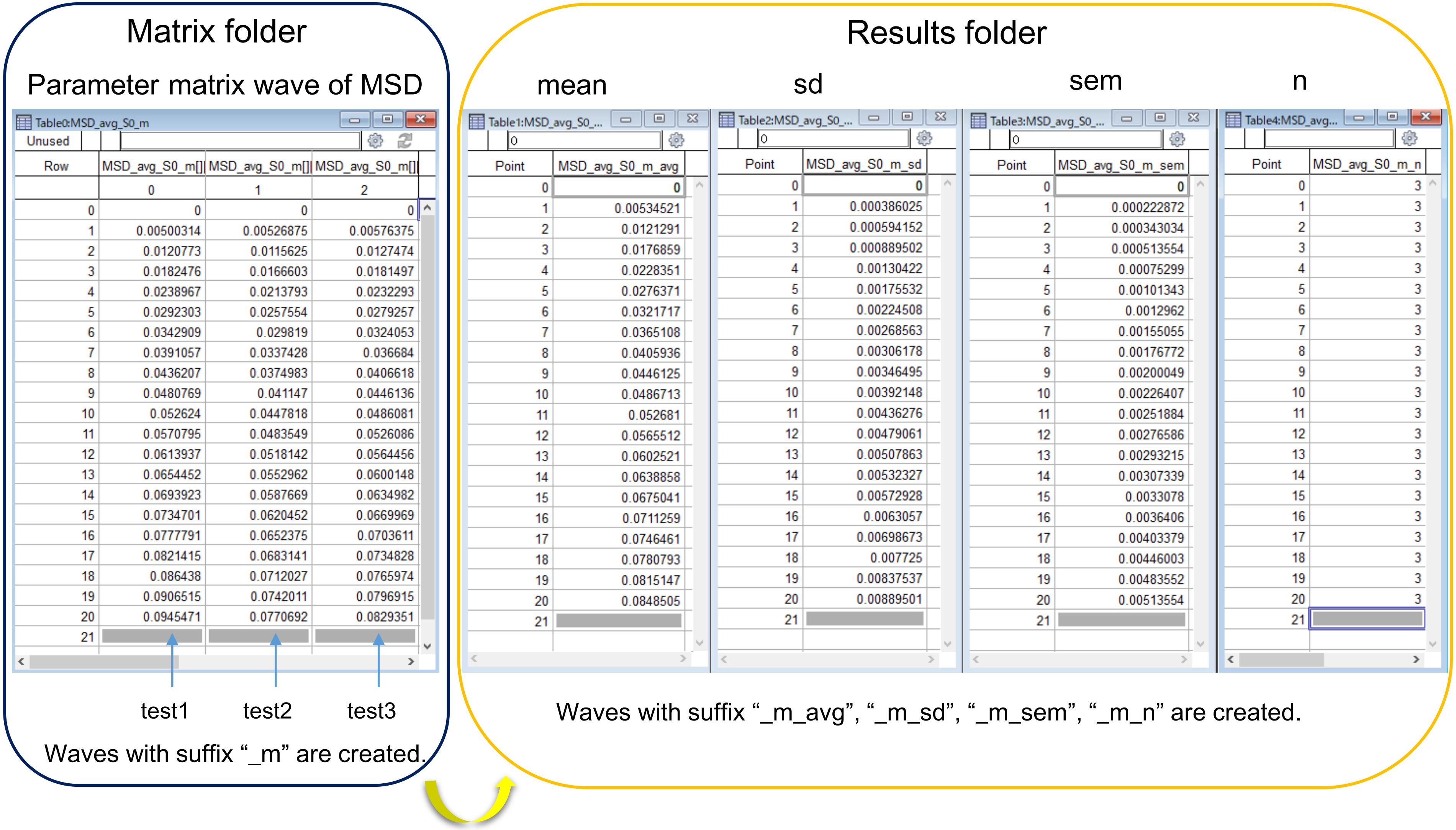


| Row | MSD_avg_S0_m[]( 0 | MSD_avg_S0_m[]( 1 | MSD_avg_S0_m[]( 2 |
|---|---|---|---|
| 0 | 0 | 0 | 0 |
| 1 | 0.00500314 | 0.00526875 | 0.00576375 |
| 2 | 0.0120773 | 0.0115625 | 0.0127474 |
| 3 | 0.0182476 | 0.0166603 | 0.0181497 |
| 4 | 0.0238967 | 0.0213793 | 0.0232293 |
| 5 | 0.0292303 | 0.0257554 | 0.0279257 |
| 6 | 0.0342909 | 0.029819 | 0.0324053 |
| 7 | 0.0391057 | 0.0337428 | 0.036684 |
| 8 | 0.0436207 | 0.0374983 | 0.0406618 |
| 9 | 0.0480769 | 0.041147 | 0.0446136 |
| 10 | 0.052624 | 0.0447818 | 0.0486081 |
| 11 | 0.0570795 | 0.0483549 | 0.0526086 |
| 12 | 0.0613937 | 0.0518142 | 0.0564456 |
| 13 | 0.0654452 | 0.0552962 | 0.0600148 |
| 14 | 0.0693923 | 0.0587669 | 0.0634982 |
| 15 | 0.0734701 | 0.0620452 | 0.0669969 |
| 16 | 0.0777791 | 0.0652375 | 0.0703611 |
| 17 | 0.0821415 | 0.0683141 | 0.0734828 |
| 18 | 0.086438 | 0.0712027 | 0.0765974 |
| 19 | 0.0906515 | 0.0742011 | 0.0796915 |
| 20 | 0.0945471 | 0.0770692 | 0.0829351 |
| 21 | | | |

| Point | MSD_avg_S0_m_avg | MSD_avg_S0_m_sd | MSD_avg_S0_m_sem | MSD_avg_S0_m_n |
|---|---|---|---|---|
| 0 | 0 | 0 | 0 | 3 |
| 1 | 0.00534521 | 0.000386025 | 0.000222872 | 3 |
| 2 | 0.0121291 | 0.000594152 | 0.000343034 | 3 |
| 3 | 0.0176859 | 0.000889502 | 0.000513554 | 3 |
| 4 | 0.0228351 | 0.00130422 | 0.00075299 | 3 |
| 5 | 0.0276371 | 0.00175532 | 0.00101343 | 3 |
| 6 | 0.0321717 | 0.00224508 | 0.0012962 | 3 |
| 7 | 0.0365108 | 0.00268563 | 0.00155055 | 3 |
| 8 | 0.0405936 | 0.00306178 | 0.00176772 | 3 |
| 9 | 0.0446125 | 0.00346495 | 0.00200049 | 3 |
| 10 | 0.0486713 | 0.00392148 | 0.00226407 | 3 |
| 11 | 0.052681 | 0.00436276 | 0.00251884 | 3 |
| 12 | 0.0565512 | 0.00479061 | 0.00276586 | 3 |
| 13 | 0.0602521 | 0.00507863 | 0.00293215 | 3 |
| 14 | 0.0638858 | 0.00532327 | 0.00307339 | 3 |
| 15 | 0.0675041 | 0.00572928 | 0.0033078 | 3 |
| 16 | 0.0711259 | 0.0063057 | 0.0036406 | 3 |
| 17 | 0.0746461 | 0.00698673 | 0.00403379 | 3 |
| 18 | 0.0780793 | 0.007725 | 0.00446003 | 3 |
| 19 | 0.0815147 | 0.00837537 | 0.00483552 | 3 |
| 20 | 0.0848505 | 0.00889501 | 0.00513554 | 3 |
| 21 | | | | |

Note: At least 3 data are required to use the stats macro.

Fig. S21 Work flow of the “Comparison Analysis” and “Multiple comparison test” macros

#### Comparison Analysis

- **MSD-dt**: Output the MSD-Δt plot (mean ± sem of the cells).
- **$D_{MSD\text{-}dt}$**: Output violin/box plot of the diffusion coefficient (D) estimated from the MSD-Δt plots.
- **$L_{MSD\text{-}dt}$**: Output violin/box plot of the confinement length (L) estimated from the MSD-Δt plots.
- **$D_{ratio}$**: Output violin/box plot of the diffusion state ratio estimated from the displacement histogram analysis.
- **$D_{state}$**: Output violin/box plot of the D of each state estimated from the displacement histogram analysis.
- **Intensity**: Output the intensity analysis results.
- **Density**: Output violin/box plot of density analysis results.
- **Off rate**: Output decay curves of the duration of trajectories and violin/box plots of the off rate parameters.
- **On rate**: Output violin/box plot of the on rate parameters.
- **2D plots**: Output 2D plots of the diffusion, intensity, and density parameters of the single-cell.

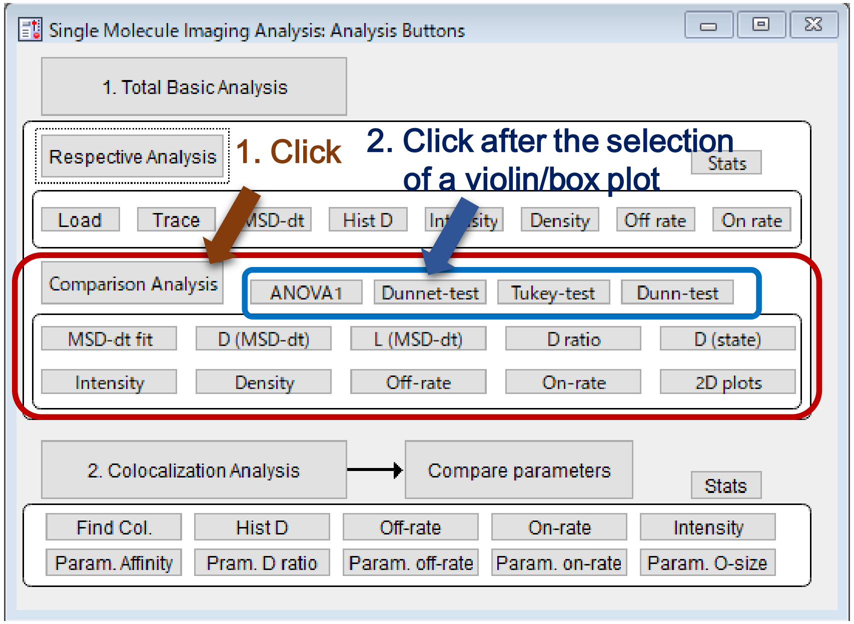


#### Multiple comparison test

- **ANOVA**: Output result table of one-way ANOVA among groups in a selected violin plot.
- **Dunnett**: Output result table of Dunnett's test among groups in a selected violin plot.
- **Tukey**: Output result table of Tukey's test among groups in a selected violin plot.
- **Dunn**: Output result table of Dunn-Holland-Wolfe’s test among groups in a selected violin plot. (Non-parametric)

Fig. S22 Example of the violin/box plots (MSD-Δt plot analysis) and the multiple comparison test

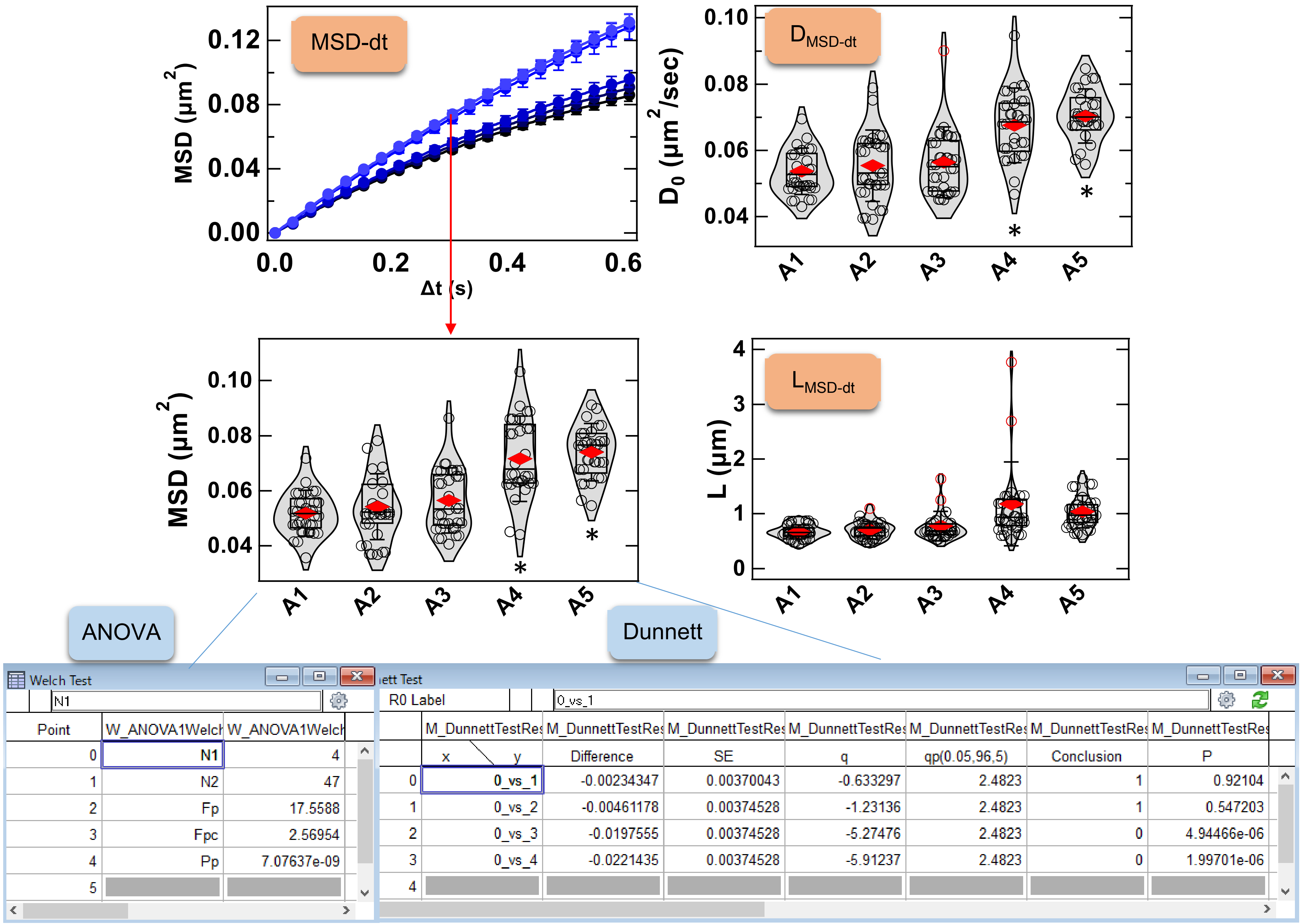


| Point | W_ANOVA1Welch | W_ANOVA1Welch |
|---|---|---|
| 0 | N1 | 4 |
| 1 | N2 | 47 |
| 2 | Fp | 17.5588 |
| 3 | Fpc | 2.56954 |
| 4 | Pp | 7.07637e-09 |
| 5 | | |

| | M_DunnettTestRes | M_DunnettTestRes | M_DunnettTestRes | M_DunnettTestRes | M_DunnettTestRes | M_DunnettTestRes | M_DunnettTestRes |
|---|---|---|---|---|---|---|---|
| | x \ y | Difference | SE | q | qp(0.05,96,5) | Conclusion | P |
| 0 | 0_vs_1 | -0.00234347 | 0.00370043 | -0.633297 | 2.4823 | 1 | 0.92104 |
| 1 | 0_vs_2 | -0.00461178 | 0.00374528 | -1.23136 | 2.4823 | 1 | 0.547203 |
| 2 | 0_vs_3 | -0.0197555 | 0.00374528 | -5.27476 | 2.4823 | 0 | 4.94466e-06 |
| 3 | 0_vs_4 | -0.0221435 | 0.00374528 | -5.91237 | 2.4823 | 0 | 1.99701e-06 |
| 4 | | | | | | | |
